# Predicting range shifts of African apes under global change scenarios

**DOI:** 10.1101/2020.06.25.168815

**Authors:** Joana S. Carvalho, Bruce Graham, Gaёlle Bocksberger, Fiona Maisels, Elizabeth A. Williamson, Serge Wich, Tenekwetche Sop, Bala Amarasekaran, Richard A. Bergl, Christophe Boesch, Hedwige Boesch, Terry M. Brncic, Bartelijntje Buys, Rebecca Chancellor, Emmanuel Danquah, Osiris A. Doumbé, Anh Galat-Luong, Jessica Ganas, Sylvain Gatti, Andrea Ghiurghi, Annemarie Goedmakers, Nicolas Granier, Dismas Hakizimana, Barbara Haurez, Josephine Head, Ilka Herbinger, Annika Hillers, Sorrel Jones, Eno-Nku Manasseh, Maureen S. McCarthy, Bethan J. Morgan, Stuart Nixon, Louis Nkembi, Emmanuelle Normand, Sarah H. Olson, Leon Payne, Charles-Albert Petre, Alex K. Piel, Lilian Pintea, Andrew J. Plumptre, Aaron Rundus, Adeline Serckx, Fiona A. Stewart, Jacqueline Sunderland-Groves, Nikki Tagg, Angelique Todd, Ashley Vosper, José Francisco Carminatti Wenceslau, Erin G. Wessling, Jacob Willie, Hjalmar Kühl

## Abstract

**Aim:** Modelling African great ape distribution has until now focused on current or past conditions, whilst future scenarios remain scarcely explored. Using an ensemble forecasting approach, we predicted changes in taxon-specific distribution under future scenarios of climate, land-use and human population changes.

**Location:** Sub-Saharan Africa

**Methods:** We compiled occurrence data on African ape populations from the IUCN A.P.E.S. database and extracted relevant human-, climate- and habitat-related predictors representing current and future (2050) conditions to predict taxon-specific distribution under a best- and a worst-case scenario, using ensemble forecasting. Given the large effect on model predictions, we further tested algorithm sensitivity by considering *default* and *non-default* modelling options. The latter included interactions between predictors and polynomial terms in correlative algorithms.

**Results:** The future distributions of gorilla and bonobo populations are likely to be directly determined by climate-related variables. In contrast, future chimpanzee distribution is influenced mostly by anthropogenic variables. Both our modelling approaches produced similar model accuracy, although a slight difference in the magnitude of range change was found for *Gorilla beringei beringei, G. gorilla diehli*, and *Pan troglodytes schweinfurthii*. On average, a decline of 50% of the geographic range (*non-default*; or 55% *default*) is expected under the best scenario if no dispersal occurs (57% *non-default* or 58% *default* in worst scenario). However, new areas of suitable habitat are predicted to become available for most taxa if dispersal occurs (81% or 103% best, 93% or 91% worst, *non-default* and *default*, respectively), except for *G. b. beringei*.

**Main Conclusions:** Despite the uncertainty in predicting the precise proportion of suitable habitat by 2050, both modelling approaches predict large range losses for all African apes. Thus, conservation planners urgently need to integrate land-use planning and simultaneously support conservation and climate change mitigation measures at all decision-making levels both in range countries and abroad.

## INTRODUCTION

Currently, a major conservation challenge is to assess the potential future effects of climate and land-use changes on species distributions, typically through the use of species distribution models (SDMs), and usually under a range of future environmental scenarios. SDMs are widely used to predict and map species’ ecological niches through time and space (Elith & Leathwick, 2009; Guillera-Arroita et al., 2015; Hao, Elith, Guillera-Arroita, & Lahoz-Monfort, 2019). Importantly, SDMs can inform spatial prioritization decisions for conservation and management actions, such as identification of strategic locations for new conservation or survey sites, and predicting future distributions (Araújo & New, 2007; Elith & Leathwick, 2009; Guillera-Arroita et al., 2015).

Changes in climate and land use are among the main global threats to biodiversity, therefore, how the synergistic interactions between these drivers impact species is an important area of research (Oliver & Morecroft, 2014). Newbold et al. (2018) assessed the single and combined effects of future climate and land-use change on local vertebrate biodiversity. They found that climate change is likely to be the principal driver of species distribution change in coming decades, equalling or surpassing the potential effects of land-use change by 2070. Because human population growth is already an extinction threat to many species (McKee, Chambers, & Guseman, 2013), it is important to determine how human distribution will impact future species presence (Jones & O’Neill, 2016).

Many primates are facing imminent extinction, due to extensive habitat loss and fragmentation, land-use change, global commodity growth and trade, and hunting (Estrada et al., 2018). Climate change is a delocalised, multi-faceted driver to add to the list. It exposes many species, especially forest-dwelling primates, to climatically unsuitable conditions (Carvalho et al., 2019). Primates have relatively limited dispersal abilities for their body size, slow reproduction, low population densities, dietary requirements and poor thermoregulation, and a predicted reduction of up to 86% of the Neotropical primate ranges with >3°C warming is likely to constrain their dispersal, resulting in elevated risks of extinction (Carvalho et al., 2019).

All African great apes (hereafter African apes) are classified either as Endangered (mountain gorillas *G. b. beringei*, bonobos *Pan paniscus*, Nigeria-Cameroon chimpanzees *P. t. ellioti*, eastern chimpanzees *P. t. schweinfurthii* and central chimpanzees *P. t. troglodytes*) or Critically Endangered (Grauer’s gorillas *G. b. graueri*, Cross River gorillas *G. g. diehli*, western lowland gorillas *G. g. gorilla* and western chimpanzees *P. t. verus*) on the IUCN Red List of Threatened Species (www.iucnredlist.org) and are regarded as flagship species for conservation. African apes have faced dramatic changes in suitable environmental conditions over the past 20 years (Junker et al., 2012) as well as large population losses (Kuehl et al., 2017; Plumptre et al., 2016; Strindberg et al., 2018) caused by human activities and/or infectious epidemics (Walsh et al., 2003). Many African apes live in areas that are suitable for agricultural expansion and 58.7% of oil-palm concessions currently overlap with African ape ranges (Wich et al., 2014). Moreover, massive development corridors (Heinicke et al., 2019) and mining activities (Howard, 2019) in their geographic ranges are projected to expand considerably, and to disrupt ape habitat connectivity and accelerate habitat loss.

Most African apes occur outside protected areas (Freeman, Roehrdanz, & Peterson, 2018; Heinicke et al., 2019; Strindberg et al., 2018). Importantly, protected areas will not be exempt from climate change (Araújo, Alagador, Cabeza, Nogués-Bravo, & Thuiller, 2011), and shifts in species ranges as predicted by future scenarios would certainly determine the degree of species representation within protected areas. Improving the effectiveness of conservation efforts in protected areas as well as habitat connectivity would allow apes to disperse to new climatically suitable areas, and favour ape population survival in the long term.

A few studies have examined the future effects of climate change on African apes (Lehmann, Korstjens, & Dunbar, 2010; Thorne et al., 2013). Lehmann et al. (2010) investigated how climate change is likely to influence survival and reported that chimpanzees may lose 10% of their range, and gorillas 75% by year 2100. Given the small range of mountain gorillas and their highly restricted occurrence in mountain refuges as a result of human encroachment and the geographic barrier of the Rift Valley, one would expect them to be particularly susceptible to global warming and extinction (Thorne et al., 2013). However, in general, how future synergetic interactions among climate, land-use and human population changes will affect African apes and their habitat has been largely unexplored.

Here we combine data on projected climate, land-use and human population changes to model African ape distribution for the year 2050. We use the most comprehensive database on ape populations available, the IUCN SSC Ape Populations, Environments and Surveys database (A.P.E.S.) to predict the distribution of great apes on the African continent under best- and worst-case scenarios. We subsequently consider an ensemble forecasting approach to reduce the uncertainty among different models and future scenarios (Araújo & New, 2007; Thuiller, 2004) and estimate the proportional change in range size in 2050 relative to current estimated range sizes for African apes.

## METHODS

### African ape data

We compiled information on African ape occurrence held in the IUCN SSC A.P.E.S. database, a repository that includes a remarkable amount of information on population status, threats and conservation for several hundred sites (Heinicke et al., 2019; Kuehl, Williamson, Sanz, Morgan, & Boesch, 2007). We extracted all occurrence data, which are georeferenced point data of direct sightings and great ape sign (mostly night nests) collected over 20 years (1998-2017, see Appendix S1 in Supporting Information, Table S1.1). We obtained a total of 62,469 presence records across all African ape taxa (occurrence data for each species in Table S1.1).

Although these data may be spatially biased as sampling effort is unevenly spread over the ape range, presence-only data are commonly the most available and hence most frequently used in SDMs (Phillips et al., 2009). The (sub)species (hereafter taxon) occurrence data we used were collected during systematic site-based wildlife and human impact surveys, which were generally based on some prior knowledge of occurrence, often in or close to protected areas, FSC-certified and other logging concessions, and from habituated populations, factors that can distort an SDM (Phillips et al., 2009). Different approaches have been applied to account for biased datasets: random background, bias background, geographic thinning/filtering, and environmental filtering (Aiello-Lammens, Boria, Radosavljevic, Vilela, & Anderson, 2015; Fourcade, Engler, Rödder, & Secondi, 2014; Phillips et al., 2009; Varela, Anderson, García-Valdés, & Fernández-González, 2014). Thus, we considered all approaches, and for the bias background distances to roads, protected areas and villages were included. We favoured the approach with the best performance by visually inspecting the greatest overlap between taxon occurrence and each sampling bias (Fig. S1.1). Given that the geographic thinning approach performed best for all taxa, we integrated it into the SDMs for sampling bias correction (Fig. S1.1, Table S1.1).

Model algorithms require presence and absence data, so we generated a set of 10,000 pseudo-absence occurrences (Guillera-Arroita et al., 2015; Phillips et al., 2009) in the range of each taxon, except for *G. b. beringei*. Only 1,000 background occurrences were created for mountain gorillas due to their small range.

We delineated taxon-specific study regions to avoid unrealistic geographical predictions (Anderson & Gonzalez, 2011). For this, we created buffers bounding IUCN range polygons (IUCN, 2018) and included all occurrence data for each taxon (Table S1.1) (Jantz, Pintea, Nackoney, & Hansen, 2016; Junker et al., 2012; Thorne et al., 2013). Whenever the buffer overpassed a known geographic barrier to ape dispersal (e.g. major rivers), we disregarded that area.

### Predictor variables

We selected predictor variables based on their importance for African ape ecology, whilst guaranteeing data availability for current and future conditions (2050) under best- and worst-case scenarios and minimizing correlation between variables. We compiled altitude and climatic variables (*N*=19) for the present and future conditions from Worldclim (periods of 1950-2000 and 2050, respectively; Table S1.1, (Hijmans, Cameron, Parra, Jones, & Jarvis, 2005)). For future predictions, we chose a best-case scenario (i.e. high mitigation scenario, CCSM4 RCP 4.5) and a worst-case scenario (i.e. low mitigation scenario, HadGEM-ES RCP 8.5) (for more details see (Carvalho et al., 2019)). Land-use/cover data for current conditions and 2050 projections were compiled from the Land-use Harmonization Project (period of 1500-2100, Table S1.1, (Chini, Hurtt, & Frolking, 2014; Hurtt et al., 2011)). This dataset represents a set of land-use change and emission scenarios for studies of human impact on the past and future global carbon-climate system. Again, we considered a best-case scenario (MiniCam RCP 4.5) and a worst-case scenario (MESSAGE RCP 8.5) (Carvalho et al., 2019). We focused on the land-use states that best represent biomes where great apes can be found: primary (i.e. natural vegetation (either forest or non-forest) undisturbed by humans), secondary (i.e. natural vegetation previously disturbed by agriculture or wood harvesting), and cropland.

We based human population scenarios on a new set of future societal development scenarios, namely Shared Socioeconomic Pathways (SSP) (Table S1.1) (Jones & O’Neill, 2016). These future scenarios are based on both qualitative narratives of future development and quantitative projections of key elements such as human population growth at the national level, educational composition, urbanization and economic growth. These data are available from 2010 to 2100 for urban and rural population. We used two future scenarios, SSP1 and SSP3, given that they represent best- and worst-case scenarios, respectively.

We also considered distances to roads, villages and rivers as they are known to influence the distribution of African apes (Table S1.1) (Carvalho, Marques, & Vicente, 2013). We extracted data on protected areas in each taxon distribution from the most comprehensive global database on terrestrial and marine protected areas, the World Database on Protected Areas (Table S1.1).

Firstly, we extracted all variables for the extent of the range of each taxon, resampled onto a 5km x 5km equal-area grid and projected them into the WGS 1984 geographic coordinate system. Secondly, we used Spearman rank correlations to select a subset of least correlated variables to minimize multicollinearity (Brun et al., 2019). For this, we used a graphical representation of the correlation values between variables to identify five least correlated variables for each species to avoid overfitting in model predictions (Fig. S1.2) (Thorne et al., 2013). We performed data analyses using the software R version 3.6.1 (R Development Core Team, 2019) and ArcMap version 10.4.1 (ESRI, 2011).

### SDM performance and ensemble forecasting

We predicted future African ape distributions using an ensemble forecasting approach (i.e. combining predictions from individual models into an ensemble as implemented in the ‘biomod2’ package in R (version 3.3-7)) (Thuiller, Georges, & Engler, 2016). We selected two correlative algorithms, generalised linear model (GLM) and generalised additive model (GAM), and three machine-learning techniques, Maxent, random forest (RF) and artificial neural networks (ANN) to build predictive SDMs for each species. These algorithms have been shown to perform well in previous SDMs (Elith et al., 2006; Thuiller, Lafourcade, Engler, & Araújo, 2009). As the choice of configuration settings of individual modelling algorithms has potentially considerable impacts on predicted distributions (Hallgren, Santana, Low-Choy, Zhao, & Mackey, 2019), we contrasted results under two modelling approaches by 1) using the default tuning options of algorithms (i.e. *default*), and 2) changing the configuration settings of algorithms (i.e. *non-default*). The latter included changes in the correlative algorithms, particularly defining interactions between predictors as well as polynomial terms, to better represent assumptions about the ecological niche of each taxon (see R code in Appendix S2).

For the present time period only, we assessed the predictive performance of each model through cross validation using a bootstrap approach, i.e. partitioning of the presence data, using 80% of presences, randomly selected, for model calibration and 20% for evaluation, and repeating this procedure 5 times (Thuiller et al., 2009). We evaluated the performance of each model by the ‘true skill statistic’ metric (TSS) (Allouche, Tsoar, & Kadmon, 2006). TSS is an accuracy measure that accounts both for omission errors (i.e. the percentage of true presences predicted as absences are minimised) and commission errors (i.e. the percentage of true absences predicted as presences are minimised), is unaffected by prevalence, and ranges from −1 to 1, with a prediction accuracy considered similar to ‘random’ when ≤0, ‘poor’ in the range 0.2-0.5, ‘useful’ in the range 0.6-0.8, and ‘good’ to ‘excellent’ when >0.8 (Allouche et al., 2006).

Ensemble forecasting has been widely employed to reduce the uncertainties associated with using a single algorithm, and is a useful method to account for uncertainties of extrapolation of species-environment relationships outside the environments sampled by the species data (Araújo & New, 2007; Hao et al., 2019; Thuiller, Guéguen, Renaud, Karger, & Zimmermann, 2019; Thuiller et al., 2009). We chose to apply the weighted mean ensemble method, which scales predictions of different models by weights based on some measure of predictive performance (Araújo & New, 2007; Thuiller et al., 2009). We included only individual models that reached very ‘good’ predictive accuracies (TSS>0.8) in ensemble models to map the current and future habitat suitability predicted for each taxon (Thuiller et al., 2019). For each modelling approach, we repeated the modelling five times (cross-validation) and given the five modelling algorithms and the three repetitions for variable importance (see below), we obtained an ensemble of 75 predicted distributions for each species for each time period (present and 2050) and future scenarios (best- and worst-case scenarios).

### Relative importance of predictors

For each taxon and modelling approach, we calculated the importance of each predictor by correlating the fitted values of the full models with those from the model in which the values of the predictor were randomly permuted. We repeated this procedure three times (default settings of ‘biomod2’ were used) and used the average Pearson’s correlation to measure variable importance. A high correlation between the values from the full and permuted models indicates that the variable has a low importance, contributing poorly to the model. We then ranked each variable value based on the correlation coefficients and reversed its relative importance and scaled from 0 to 1, the more influential variables for the model representing those with a higher relative importance (Thuiller et al., 2009).

### Species range change

We estimated the proportional change in range size, in 2050 compared to the present, for each taxon by subtracting the future prediction ensemble output from the SDMs for the best- and worst-case scenarios from that under current conditions. We considered continuous predictive outputs from ensemble models as they provide richer information over outputs classified into binary maps, particularly when the purpose of our study is to inform spatial prioritization decisions for conservation and management actions (Guillera-Arroita et al., 2015).

We subsequently identified areas of range loss (i.e. sites where the species is present at the moment but is likely to be absent in the future), gain (i.e., sites where the species is absent at the moment but is likely to be present in the future), and stability (i.e., sites where the species is potentially present at the moment and is likely to be present in the future). For this, we considered range change under two contrasting dispersal scenarios: 1) full dispersal, which assumes that the species can disperse to new suitable areas in the future; and 2) no dispersal, which assumes that the species will be unable to disperse and only the overlap between present and future distributions will be the expected suitable habitat for the species (Thomas et al., 2004).

## RESULTS

In general, both modelling approaches (*default* and *non-default*) resulted in similar predictive accuracy of individual and ensemble models and in the relative importance of the variable predictors (Fig. 1, S1.3-4). Predictive accuracy of the individual models based on TSS was ‘useful’ to ‘excellent’, depending on the algorithm (Fig. S1.3). On average, RF models performed best relative to ANN and GLM models which performed worst at predicting species distributions (Fig. S1.3a). Importantly, with TSS scores >0.9 ensemble models had excellent predictive accuracy and clearly outperformed individual models (Fig. S1.3b).

**Figure 1.**
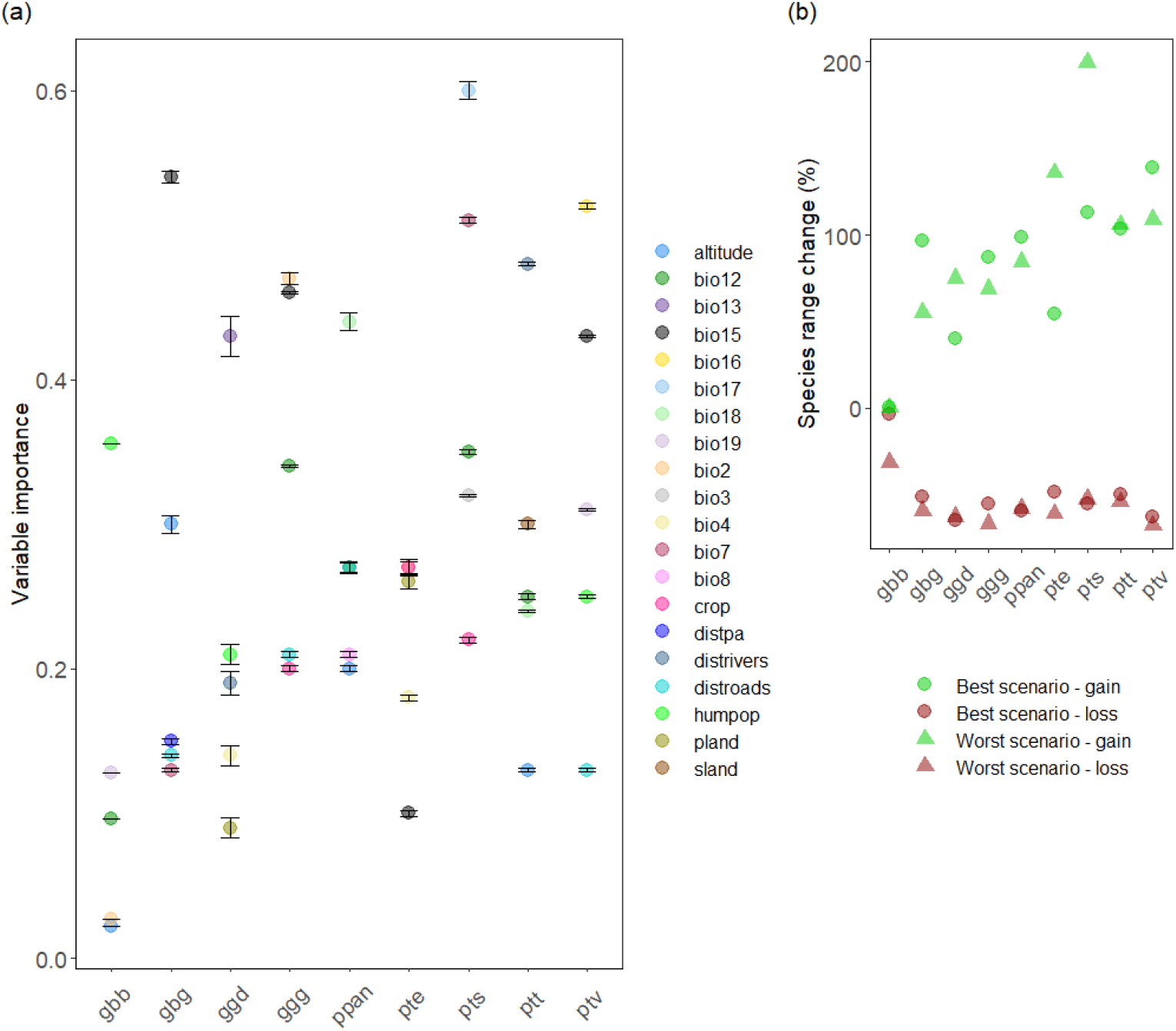
Results for the modelling approach where algorithm settings were changed for each African ape species (i.e. *non-default*). (a) Variable importance (mean and standard deviation (SD) of the correlation values) for the ensemble models, and (b) Predicted percentage change in African ape ranges by 2050 under the best- and the worst-case scenario, assuming no dispersal (loss) and dispersal (gain) scenarios. Variable predictor abbreviations: bio12 – annual precipitation, bio13 – precipitation of wettest month, bio15 – Seasonal variation of precipitation, bio16 – precipitation of wettest quarter, bio17 – precipitation of driest quarter, bio18 – precipitation of warmest quarter, bio19 – precipitation of coldest quarter, bio2 – mean diurnal range, bio3 – isothermality, bio4 – temperature seasonality, bio7 – temperature annual range, bio8 – mean temperature of wettest quarter, crop – cropland, distpa – distance to protected areas, distrivers – distances to rivers, distroads–distances to roads, humpop – human population density, pland–primary land, sland – secondary land.

Our ensemble models indicated that the current distribution of *G. b. beringei, P. t. ellioti* and *P. t. troglodytes* is strongly determined by anthropogenic variables, in contrast to the greater influence of climate-related variables on the distribution of *G. b. graueri, G. g. diehli, G. g. gorilla, P. paniscus, P. t. schweinfurthii* and *P. t. verus* (Fig. 1, S1.4-6). Our models indicate that the future distributions of most gorillas and bonobos will be more heavily influenced by climate-related variables, contrasting with the greater influence of anthropogenic variables in shaping the chimpanzee distribution (Fig. S1.5-6).

On average, the same trend in species range change was estimated for both *default* and *non-default* modelling approaches (Fig. 2, 3, S1.7-8). However, the *non-default* approach predicted a slightly lower range loss (50% or 57% under the best and worst scenarios, respectively) and lower range gain (81% or 93% under the best and worst scenarios, respectively) than the *default* approach (loss: 55% or 58%, gain: 103% or 91%, under the best and worst scenarios, respectively) (Fig. 3, S1.8). Moreover, differences were found in the magnitude of change for some taxa, particularly for *G. b. beringei, G. g. diehli* and *P. t. schweinfurthii*. To simplify, only results from the *non-default* approach are presented below (details for the *default* approach are provided in Appendix S1).

**Figure 2.**
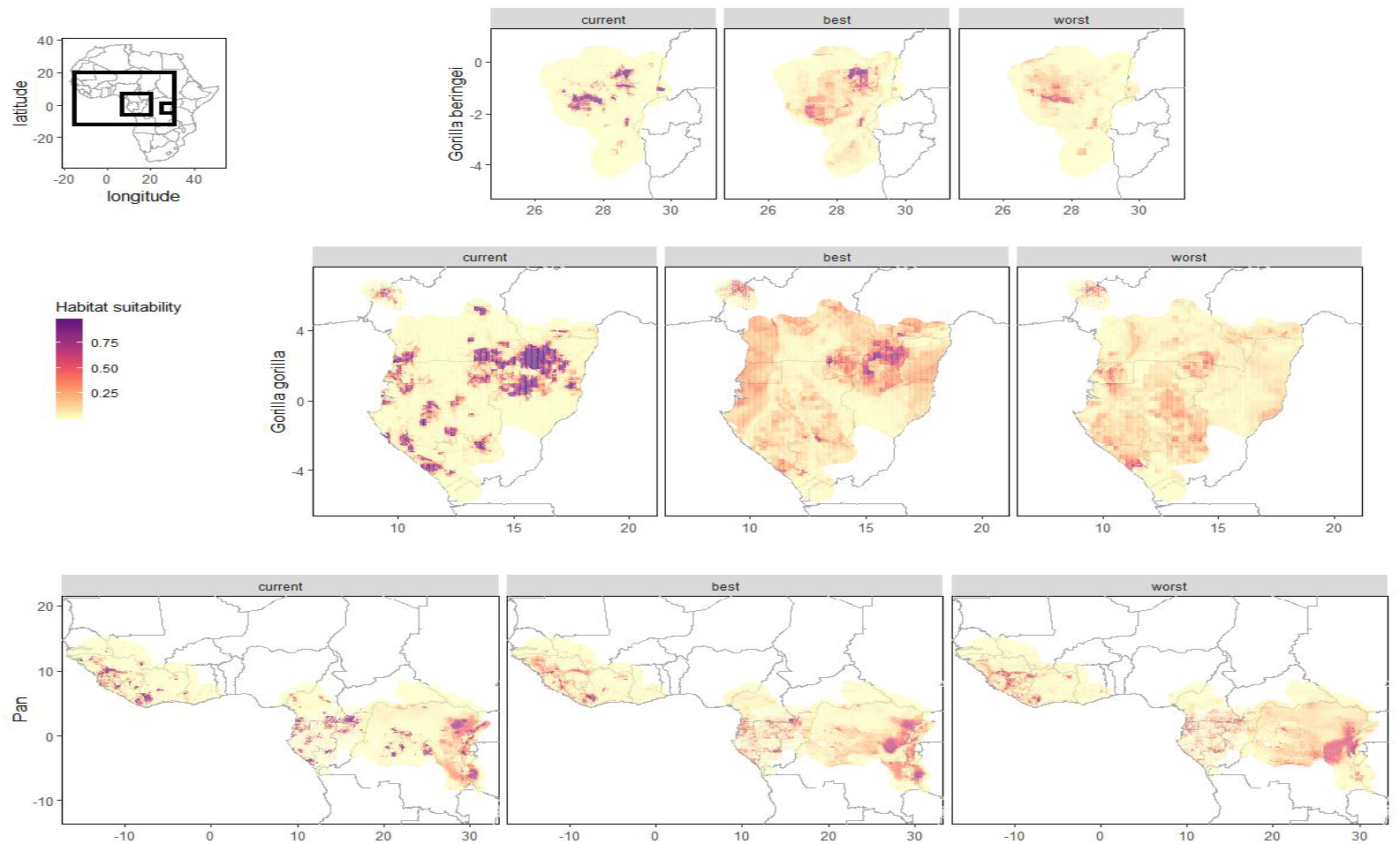
Ensemble forecasting of the current and future (best- and worst-case scenarios) habitat suitability for *Gorilla beringei* spp, *Gorilla gorilla* spp and *Pan* spp based on weighted mean and the true skill statistics (TSS) for the *non-default* approach. Note that only results for the no dispersal scenario are presented here.

**Figure 3.**
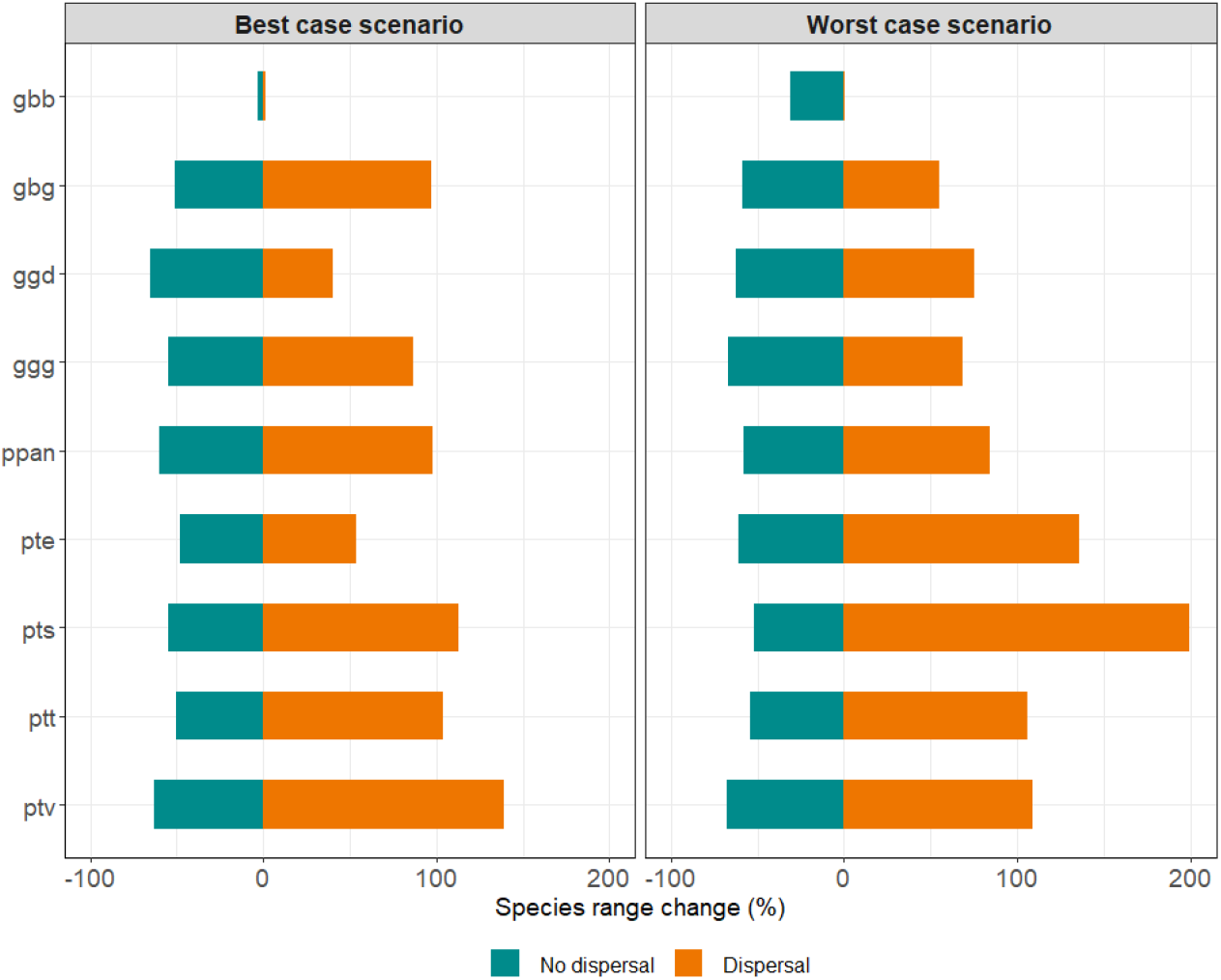
Predicted percentage change in African ape ranges by 2050 under the best- and the worst-case scenario, assuming either no dispersal (loss) and dispersal (gain) for the *non-default* approach.

### *Gorilla beringei beringei* (mountain gorilla)

Annual precipitation contributed most in the correlative models, whereas human population density was the strongest determinant of mountain gorilla distribution in machine-learning and ensemble models (Fig. S1.4). This taxon is confined to artificial ‘island’ areas in a sea of agriculture where annual precipitation (1,200-1,500 mm), diurnal temperature range (11-12 °C), human population (3,000-10,000 people/km^2^), and elevation (1,500-2,200 m) are high, and precipitation in the coldest quarter is low (180-440 mm) (Fig. S1.5-6). Precipitation variables will increase and diurnal temperature range will decrease by 2050 under the best scenario (Fig. S1.5-6). However, the opposite is predicted under the worst scenario. Human population growth is predicted to occur under both future scenarios.

Range stability is predicted to occur under the best scenario, but one-third of current suitable habitat is predicted to be lost under the worst scenario (Fig. 3, S1.7). Moreover, both future scenarios agree that dispersal is unlikely given that no new suitable habitat will become available for mountain gorillas.

### *Gorilla beringei graueri* (Grauer’s gorilla)

Seasonal variation in precipitation (precipitation seasonality) was the most important variable in both individual and ensemble models in predicting the distribution of Grauer’s gorillas (Fig. S1.4). This taxon is predicted to persist where seasonal variation of precipitation is low (25-40 mm), diurnal temperature range (12-13 °C) and altitude (500-1500 m) are high, close to roads (<6 km) and far from protected areas (<30 km) (Fig. S1.5-6). Both climatic variables are expected to decrease under the best scenario or increase under the worst scenario (Fig. S1.5-6).

If no dispersal occurs, this taxon is predicted to lose half of its current habitat under both scenarios (Fig. 3, S1.7). However, if dispersal occurs, a 97% range gain is predicted under the best scenario, but only 55% under the worst scenario.

### *Gorilla gorilla diehli* (Cross River gorilla)

Precipitation in the wettest month was the most important predictor of Cross River gorilla distribution in both individual and ensemble models (Fig. S1.4). High precipitation during the wettest month (340-400 mm) and seasonal variation in temperature (9-10 °C), very low human population (<2,500 people/km^2^), a large distance to main rivers (>400 km) and presence of large tracts of primary land (>95%) are suitable conditions for the distribution of this species (Fig. S1.5-6). Precipitation in the wettest month is predicted to increase and seasonal variation in temperature to decrease under the best scenario, whereas the opposite pattern is predicted under the worst scenario (Fig. S1.5-6). According to both future scenarios, human population is predicted to increase, and primary land is predicted to be completely lost.

If no dispersal occurs, more than two-thirds of the current range is predicted to disappear under both future scenarios (Fig. 3, S1.7). In contrast, if Cross River gorillas could disperse, range gains of 40% or 75% are predicted under the best and worst scenarios, respectively.

### *Gorilla gorilla gorilla* (western lowland gorilla)

All climatic variables were important predictors of western lowland gorilla distribution, particularly seasonal variation of precipitation and diurnal temperature range (Fig. S1.4). Areas characterised by low seasonal variation in precipitation (50-70 °C), high annual precipitation (1,600-1,800 mm) and diurnal temperature range (9-11 °C), absence of cropland and close to roads (0-15 km) harbour suitable conditions for the persistence of this subspecies (Fig. S1.5-6). According to both future scenarios, precipitation variables will not change, but decrease in diurnal temperature is predicted to occur. Cropland is predicted to increase under the worst scenario.

Assuming no dispersal, loss of more than half the suitable area is predicted under both future scenarios (Fig. 3, S1.7). With dispersal, however, a large increase in suitable habitat under both future scenarios is predicted (87% best, 69% worst).

### *Pan paniscus* (bonobo)

Precipitation in the warmest quarter is an important predictor of bonobo distribution in both individual and ensemble models (Fig. S1.4). Favourable environmental conditions shaping this species distribution are high precipitation (450-500 mm) and temperature (24-25 °C) of the warmest quarter and annual precipitation (1,750-1,950 mm), mid altitudes (350-450 m) and close proximity to roads (<6 km) (Fig. S1.5). All climatic variables are predicted to increase in the future (Fig. S1.5-6).

Under both future scenarios, more than half of the area of suitable habitat is predicted to be lost if no dispersal occurs (Fig. 3, S1.7). Suitable habitat is predicted to expand to new areas and, if bonobos disperse, range gains of 98% or 84% are predicted under the best and worst scenarios, respectively.

### *Pan troglodytes ellioti* (Nigeria-Cameroon chimpanzee)

Cropland and primary land were the best predictors in GLM and Maxent models, in contrast to diurnal temperature range, which was the variable with the highest importance in GAM and RF models as well as in the ensemble models (Fig. S1.4). Areas with a low proportion of cropland (<7%), very high proportion of primary land (>90%), high diurnal temperature range (9-11 °C), pronounced seasonal variation of both precipitation (67-75 mm) and temperature (8-11 °C) offer suitable conditions for Nigeria-Cameroon chimpanzees (Fig. S1.5). An increase in cropland and a large reduction in primary land are predicted under the worst scenario. In contrast, no cropland expansion and a smaller decrease in primary land are expected under the best scenario. Under both future scenarios, seasonal variation of precipitation and temperature are predicted to increase and diurnal temperature range to decrease (Fig. S1.5-6).

If no dispersal occurs, almost half of the area of suitable habitat is predicted to be lost under both future scenarios (Fig. 3, S1.7). However, if dispersal occurs, substantial range gains are predicted to occur, particularly under the worst scenario (137% vs. 83% best scenario).

### *Pan troglodytes schweinfurthii* (eastern chimpanzee)

Precipitation of driest quarter was an important predictor in most individual models, except for GLM and ANN models, where annual precipitation performed best (Fig. S1.4). However, all predictors ranked equally in importance in ensemble models. Eastern chimpanzees encounter suitable conditions where precipitation of the driest quarter is low (30-190 mm), annual precipitation (1,250-1,750 mm), isothermality (7-8 °C) and diurnal temperature range (13-17 °C) are high, and cropland is rare (<5%) (Fig. S1.5-6). Under the best scenario, climatic variables are unlikely to change, but a decrease in isothermality is predicted under the worst scenario. The latter also predicts expansion of cropland by 2050.

According to both future scenarios, suitable habitat is predicted to be confined to the core of the species’ current distribution (Fig. 3, S1.7). In terms of range change, half of suitable habitat is predicted to be lost under both future scenarios if no dispersal occurs. In contrast, if dispersal occurs, range expansion of 113% or 199% into new areas is expected under the best and worst scenarios, respectively.

### *Pan troglodytes troglodytes* (central chimpanzee)

Secondary land and distances to main rivers were the predictors of greatest importance in individual and ensemble models, except for GLM models, where only secondary land was a slightly better predictor (Fig. S1.4). Suitable environmental conditions for central chimpanzees are characterised by a high percentage of secondary land (12.5-75%), large distance to main rivers (>450 km), high annual precipitation (1,600-1,800 mm) and precipitation in the warmest quarter (400-500 mm), and mid-altitudes (300-600 m) (Fig. S1.5-6). According to both future scenarios, climatic variables are predicted to remain unchanged, but expansion of secondary land is expected by 2050.

A reduction of half the current area of suitable habitat is expected under both future scenarios if no dispersal occurs (Fig. 3, S1.7). Predictions of range gains for central chimpanzees suggest that 104% or 106% of suitable habitat will become available in new areas under the best and future scenarios, respectively.

### *Pan troglodytes verus* (western chimpanzee)

Precipitation in the wettest quarter was the most important variable in individual and ensemble models (Fig. S1.4). Current suitable conditions for western chimpanzees are very high precipitation of the wettest quarter (600-900 mm), high seasonal variation of precipitation (60-110 mm) and precipitation of coldest quarter (100-600 mm), low human population (<5,000 people/km^2^) and low proximity to roads (<2 km) (Fig. S1.5-6). A greater increase in both seasonal variation in precipitation and precipitation of the coldest quarter is predicted under the worst scenario.

Western chimpanzees have the widest geographic distribution among African apes, however, loss of more than half their present range is predicted under both future scenarios if no dispersal occurs (Fig. 3, S1.7). On the other hand, high range gains are anticipated under both future scenarios if there is dispersal (best: 139%, worst: 109%).

## DISCUSSION

This is the first study to combine climate, land-use and human population changes in an ensemble forecasting approach to predict African ape distribution by 2050. Customizing the model specifications relative to the default settings in the correlative algorithms revealed uncertainty in predicting the amount of suitable habitat available by 2050. However, both approaches agreed that all African ape taxa are likely to experience marked range losses (best: 55% *default* or 50% *non-default;* worst: 58% *default* or 57% *non-default*), and that these reductions may not be as severe if dispersal to new areas of suitable habitat occurs (best: 103% *default* or 81% *non-default;* worst: 91% *default* or 93% *non-default*).

A previous study quantified changes in suitable environmental conditions for African apes between 1990 and 2000, and found that the greatest proportional reductions occurred for gorillas (*G. g. diehli*, range loss of 59%; *G. b. graueri*, 52%; *G. g. gorilla*, 32%) and bonobos (*P. paniscus*, 29%) compared with chimpanzees (*P. t. troglodytes*, 17%; *P. t. verus*, 11%) (Junker et al., 2012). Lehmann and colleagues (2010) employed a mechanistic approach to investigate how climate change under a worst scenario would influence African ape survival and reported that chimpanzees might lose 10% of current range and gorillas 75%. Our study concurs with these results for most gorilla taxa, but more than half of suitable habitat area for chimpanzees is predicted to be lost under both future scenarios if no dispersal occurs. However, our full dispersal scenario predicts gains of suitable conditions in new areas under both future scenarios for all taxa, except mountain gorillas.

The variables that best predicted current distribution were taxon-specific: anthropogenic variables were key predictors in most models for mountain gorillas, Nigeria-Cameroon and central chimpanzees, and climate-related variables for Grauer’s, Cross River and western lowland gorillas, bonobos, eastern and western chimpanzees. However, future scenarios suggest that the distribution of most gorillas and bonobos is predicted to be determined by changes in climatic variables, and that of chimpanzees by land-use changes. Moreover, the forecast variation in both climatic and human population variables can explain the predicted habitat stability for mountain gorillas. Interestingly, habitat gains predicted for Cross River gorillas, Nigeria-Cameroon and eastern chimpanzees under the worst scenario are likely to be explained by stability of climatic variables and changes in anthropogenic variables by 2050.

Despite the importance of climatic variables in determining African ape time budgets, and consequently their effects on distributions, gorillas and chimpanzees are more sensitive to variations in temperature than in precipitation and they persist better in habitats with lower monthly temperature variation (Lehmann et al., 2010). Moreover, gorillas are predicted to be affected more than chimpanzees given the more restricted behavioural flexibility of gorillas to cope with temperature variation (Lehmann et al., 2010). Our study suggests that diurnal and seasonal variability influence the distribution of most gorillas, yet only play a similar role for Nigeria-Cameroon and eastern chimpanzees. Additionally, annual precipitation, and particularly its distribution over the wet and dry seasons, affects the distribution of most gorillas and chimpanzees, as well as bonobos. These results are indirect evidence of the marked influence of temperature and precipitation on species ecological niche with regard to dehydration and thermoregulation (Wessling et al., 2018). The influence of climate change on great ape ecology has been poorly explored compared with anthropogenic disturbance; nonetheless, we can expect temperature extremes to exacerbate habitat loss for African apes and consequently pose serious threats to species persistence, as they are anticipated to impact other non-primate species (Araújo et al., 2011; Newbold, 2018). By integrating future climate and land-use changes as well as human population scenarios, our predictions provide strong evidence for synergistic interactions among these global drivers constraining species distributions.

Strindberg and colleagues (2018) found that western lowland gorillas and central chimpanzees, two sympatric taxa with 97% range overlap, mostly occur outside protected areas, and argued for *‘reinforcement of anti-poaching efforts both inside and outside protected areas (particularly where habitat quality is high and human impact is low), diligent disease control measures (including training, advocacy, and research into Ebola virus disease), and the preservation of high-quality habitat through integrated land-use planning and implementation of best practices by the extractive and agricultural industries’*. Our study suggests that both taxa will find environmentally suitable areas outside their current range, in line with our predictions for most African apes. Thus, outcomes from our SDMs should be integrated with a habitat connectivity analysis to optimize conservation land-use planning and identify priority areas for these species (Freeman et al., 2018; Jones et al., 2018). This is extremely important given that African protected areas are distant from each other (Santini, Saura, & Rondinini, 2016) and particularly because great apes have a low dispersal capacity due to their slow reproduction, low population densities, dietary requirements and poor thermoregulation. It will be important to ensure objective assessments of human pressures and habitat conditions in potential protected areas to avert species extinctions in the long term (Jones et al., 2018).

Taxon-specific frameworks of environmental and socio-economic trends (Estrada et al., 2018; Strindberg et al., 2018; Tranquilli et al., 2014) should be considered at all major decision-making levels in range countries and abroad to (1) improve the management and recovery of degraded habitat in protected areas and off-reserve management to increase resilience, (2) increase the size of protected areas, establish additional protected areas and ensure habitat connectivity, (3) incorporate climate change into land-use planning and propose mitigation measures on the conservation agenda for African apes and sympatric wildlife, and (4) implement more effective law enforcement in ape ranges, supplemented by environmental education, community development, tourism programmes and research (IUCN & ICCN, 2012; IUCN, 2014; Tranquilli et al., 2014, 2012). Public-private partnerships have proven highly effective across the forest and savanna zones, where an NGO or other organisation takes on management responsibility for a given site over one or more decades (Scholte et al., 2018).

### Limitations of distribution models

Modelling species responses to global environmental changes carries many uncertainties (Araújo & New, 2007; Thuiller et al., 2019). Using two algorithm approaches, two future scenarios, two dispersal scenarios, an ensemble forecasting and including only a few but highly important predictors of the distribution of African apes, should have reduced uncertainties in our distribution models (Brun et al., 2019; Thorne et al., 2013). A recent study proposed that SDMs include historical records to produce better predictions of range shifts rather than relying on contemporary records alone (Faurby & Araújo, 2018). This is important for large vertebrates given the direct effects of anthropogenic disturbances on their distribution, and many ranges being far from equilibrium under current environmental conditions (Faurby & Araújo, 2018). The species occurrences we used are from field sites and may represent realized niches as currently African apes may be absent from areas of potential distribution, but unfortunately, reliable information about historical ranges is not available. Thus, it is possible that our predictions of current African ape distribution have underestimated the diversity of niches suitable for these species, which consequently may have limited our model predictions under future conditions.

Mining concessions and granted mining claims are increasing dramatically across Africa, particularly threatening large ape populations in Guinea, Gabon and Liberia (Howard, 2019). It will be important to model the influence of this threat on future African ape distributions once appropriate spatial data sets become available.

Our results corroborate the other recent studies showing that African ape populations and their habitats are declining dramatically (Freeman et al., 2018; Hickey et al., 2013; Kuehl et al., 2017; Lehmann et al., 2010; Plumptre et al., 2016; Strindberg et al., 2018; Thorne et al., 2013), thus our findings should also be used to guide the prioritization of conservation efforts for these flagship species to avoid irreversible losses.

### Conclusions

Given that the existing network of protected areas is inadequate for ensuring the long-term conservation of African apes (Strindberg et al., 2018), we support the argument that effective conservation strategies require taxon-specific conservation planning that focuses on existing and proposed protected areas, the creation and/or management of which can be informed by our habitat suitability models. Additionally, efforts to maintain connectivity between the habitats predicted to be suitable in the future will be crucial for the survival of African apes. For a species to shift to new climatically suitable areas, the availability of such habitat will need to be ensured through effective protection by antipoaching teams whether in a protected area or not, the enforcement of industrial extractive industry conservation regulations (for example, as currently practiced in logging concessions under FSC Standards), and by national land-use planning that ensures that the areas of current and future high conservation value are not transformed into agricultural plantations or fragmented by roads. As an example, this country-wide approach has been undertaken in Gabon, where planning for the development of agriculture, road and rail links, and mineral extraction has been informed by wildlife and vegetation data in order to locate these activities in areas that are already degraded, and to avoid closed-canopy old-growth and remote forests (Government of Gabon, 2012; Strindberg et al., 2018). This will be an effective way of promoting habitat connectivity to maintain African ape populations as well as sympatric wildlife.

## Acknowledgments

We are grateful to the governments and national authorities for research permissions, and field staff for logistical support and guidance during data collection. We would like to thank all organizations for sharing African ape survey data, particularly Antwerp Zoo Society, Chimbo Foundation, Conservation Society of Sierra Leone, Environment and Rural Development Foundation, Fauna and Flora International, Frankfurt Zoological Society, Jane Goodall Institute, Rio Tinto, Royal Society for the Protection of Birds, San Diego Zoo Global, Sekakoh Organisation, Sierra Rutile Limited, Tacugama Chimpanzee Sanctuary, The Biodiversity Consultancy, The Born Free Foundation, West African Primate Conservation Action, Wild Chimpanzee Foundation, Wildlife Conservation Society, WWF-CAR, and World Wide Fund for Nature. A special thanks to numerous individuals for data sharing: Alastair McNeilage, Abdulay Barrie, Anna Binczik, Bejnamin Barca, Catherine Tayleur, Cleve Hicks, Damien Caillaud, David H. Humminbird, Emma Stokes, Takeshi Furuichi, Gay Reinartz, Genevieve Campbell, Hilde Vanleeuwe, Jennifer Moustgaard, Joel Gamys, John Hart, Jremy Lindsell, Julian Blanc, Lilian Pintea, Mary Molokwu, Nakashima Yoshihiro, Noelle Kumpel, Nzooh Zacharie, Patrick Guislain, Paul K. N’Goran, Roger Fotso, Sebatien Regnaut, and Stephanie Latour. JSC was funded by a University of Stirling Research Programme Fellowship.

## Data Accessibility Statement

The African ape data sets are available via the IUCN SSC A.P.E.S. database (http://apesportal.eva.mpg.de).

## Biosketch

Joana Carvalho is a postdoctoral researcher in the Faculty of Science, Liverpool John Moores University, UK, who has been working in the field of primate ecology and conservation. Her current research focuses on evaluating effects of future climate change and land-use/cover change on primates, through species distribution modelling, scenario analyses, and landscape connectivity analyses. Additionally, she has been working on density/distribution models for Tanzania chimpanzees.

## Notes

### Competing Interest Statement

The authors have declared no competing interest.

