## Appendix 1 for "Predicting range shifts of African apes under global change scenarios"

### APPENDIX S1

Table S1.1. Information about the African ape data from the IUCN SSC A.P.E.S. database. Also, the size (km) of buffers bounding IUCN range polygons and the distance (km) between points in geographical thinning layer (bias correction) as well as the predictor variables used in SDMs and corresponding sources are shown.

| Species | Subspecies | Temporal coverage | No. occurrences | Evidences* | Buffer | Bias layer** | Predictor variables*** | Source |
|---|---|---|---|---|---|---|---|---|
| *Gorilla beringei* | *beringei* | 2002 & 2006 | 266 | DO, Du, Ne, Tr | 20 | 1 | Annual precipitation<br>Precipitation coldest quarter<br>Diurnal temperature range<br>Human population<br>Altitude | www.worldclim.org/CMIP5<br>www.worldclim.org<br>www.worldclim.org<br>www.cgd.ucar.edu<br>www.worldclim.org |
| | *graueri* | 2002→2014 | 1,869 | Du, FR, FP, Ne, Vo | 50 | 5 | Seasonal variation precipitation<br>Annual temperature range<br>Altitude<br>Distances to PAs<br>Distances to roads | www.worldclim.org<br>www.worldclim.org<br>www.worldclim.org<br>www.protectedplanet.net<br>www.sedac.ciesin.columbia.edu |
| *Gorilla gorilla* | *diehli* | 1998→2013 | 909 | DO, Du, FR, FP, Ne, Tr, RP, Vo | 50 | 1 | Precipitation wettest month<br>Seasonal variation temperature<br>Primary land<br>Distances to rivers<br>Human population | www.worldclim.org<br>www.worldclim.org<br>www.daac.ornl.gov<br>www.hydrosheds.org<br>www.cgd.ucar.edu |
| | *gorilla* | 2001→2016 | 18,723 | DO, Du, FR, FP, Ne, Tr, Vo | 50 | 10 | Annual precipitation<br>Seasonal variation precipitation<br>Diurnal temperature range<br>Distances to roads<br>Cropland | www.worldclim.org<br>www.worldclim.org<br>www.worldclim.org<br>www.sedac.ciesin.columbia.edu<br>www.daac.ornl.gov |

| | | | | | | | | |
|---|---|---|---|---|---|---|---|---|
| *Pan paniscus* | | 2004→2012 | 4,683 | DO, Du, FR, FP, Ne, Tr, Vo | 50 | 5 | Annual precipitation<br>Precipitation warmest quarter<br>Temperature wettest quarter<br>Altitude<br>Distances to roads | www.worldclim.org<br>www.worldclim.org<br>www.worldclim.org<br>www.worldclim.org<br>www.sedac.ciesin.columbia.edu |
| | *ellioti* | 1998→2015 | 1,222 | DO, Du, FR, FP, Ne, Pl, Tr, RP, Vo | 50 | 5 | Seasonal variation precipitation<br>Seasonal variation temperature<br>Diurnal temperature range<br>Primary land<br>Cropland | www.worldclim.org<br>www.worldclim.org<br>www.worldclim.org<br>www.daac.ornl.gov<br>www.daac.ornl.gov |
| | *schweinfurthii* | 1999→2014 | 12,294 | DO, Du, FR, FP, Ne, Vo | 70 | 10 | Annual precipitation<br>Precipitation driest quarter<br>Annual temperature range<br>Isothermality<br>Cropland | www.worldclim.org<br>www.worldclim.org<br>www.worldclim.org<br>www.worldclim.org<br>www.daac.ornl.gov |
| *Pan troglodytes* | *troglodytes* | 1998→2015 | 13,020 | DO, Du, FR, FP, Ne, Tr, Vo | 50 | 10 | Annual precipitation<br>Precipitation warmest quarter<br>Altitude<br>Distances to rivers<br>Secondary land | www.worldclim.org<br>www.worldclim.org<br>www.worldclim.org<br>www.hydrosheds.org<br>www.daac.ornl.gov |
| | *verus* | 1998→2017 | 14,166 | DO, Du, FR, FP, Ne, Tr, Vo | 200 | 10 | Seasonal variation precipitation<br>Precipitation wettest quarter<br>Precipitation coldest quarter<br>Distances to roads<br>Human population | www.worldclim.org<br>www.worldclim.org<br>www.worldclim.org<br>www.sedac.ciesin.columbia.edu<br>www.cgd.ucar.edu |

[*] DO - Direct observation, Du – Dung, FR - Feeding remains, FP – Footprint, Ne – Nest, Pl- Playground, RP - Resting place, Tr - Trail/passage, Vo – Vocalization; [**] Geographical thinning performed best for all species (see methods, Fig. S1); [***] See methods for detailed information about how these variables were chosen.

*Gorilla beringei beringei*

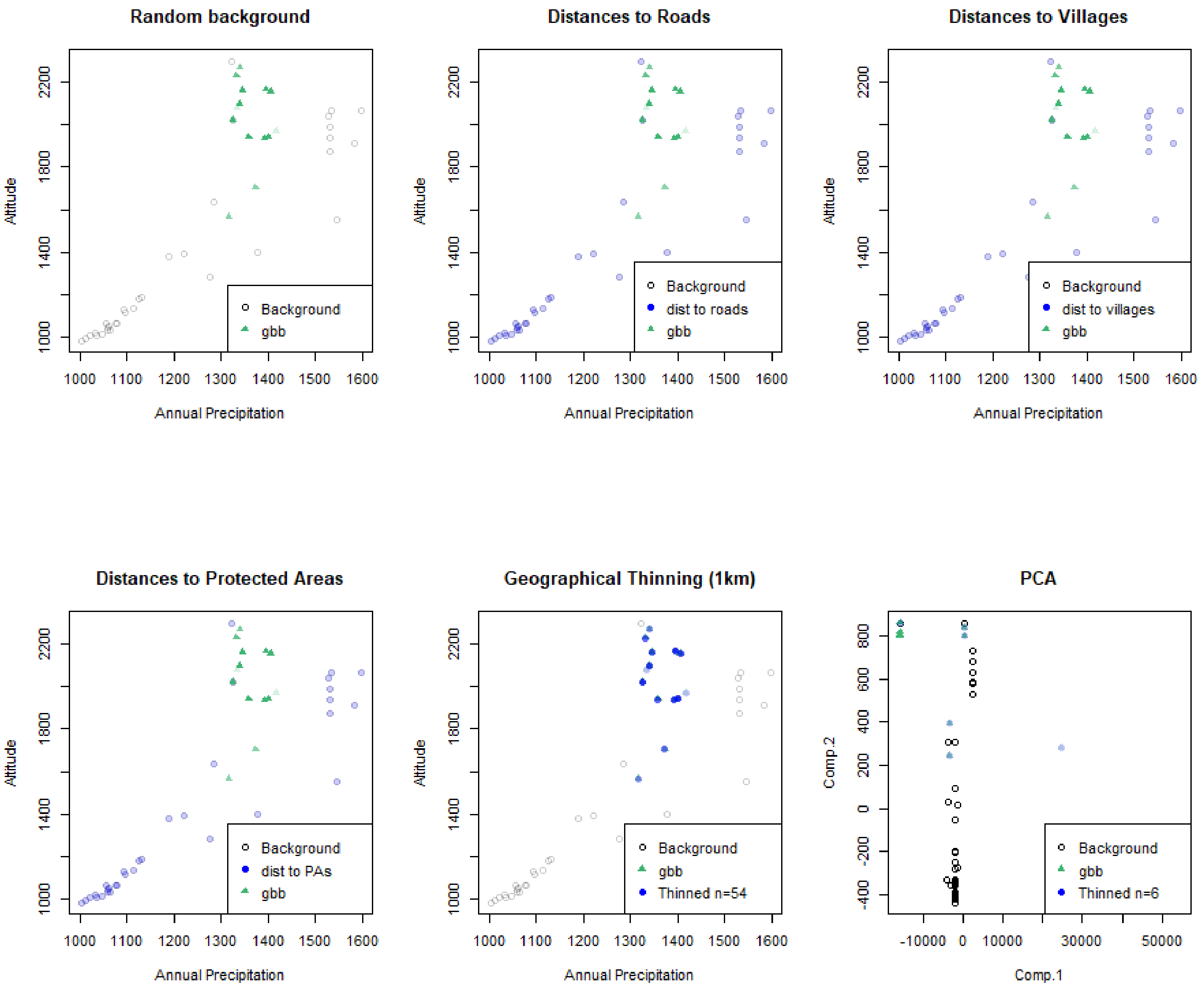

*Gorilla beringei graueri*

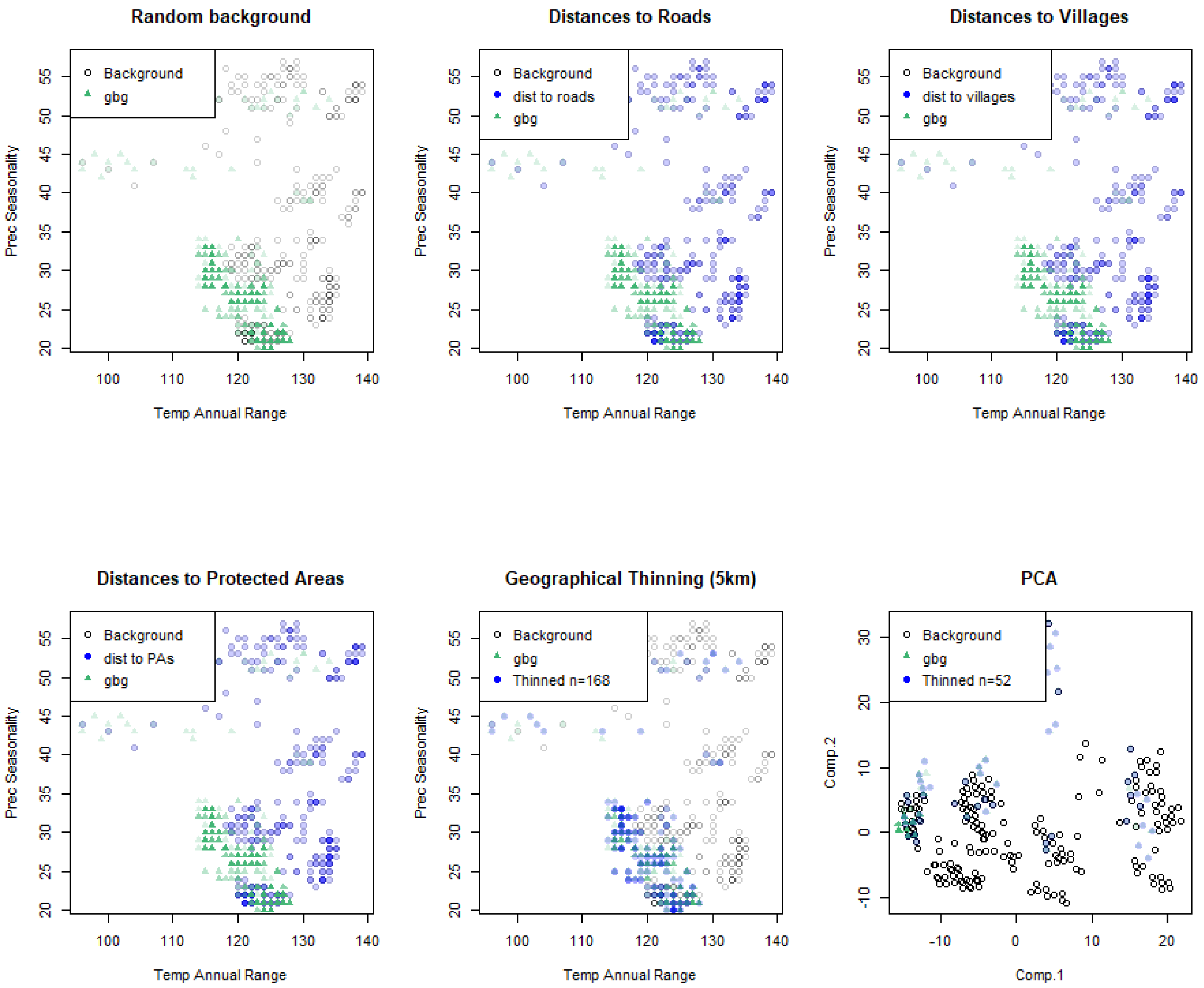

*Gorilla gorilla diehli*

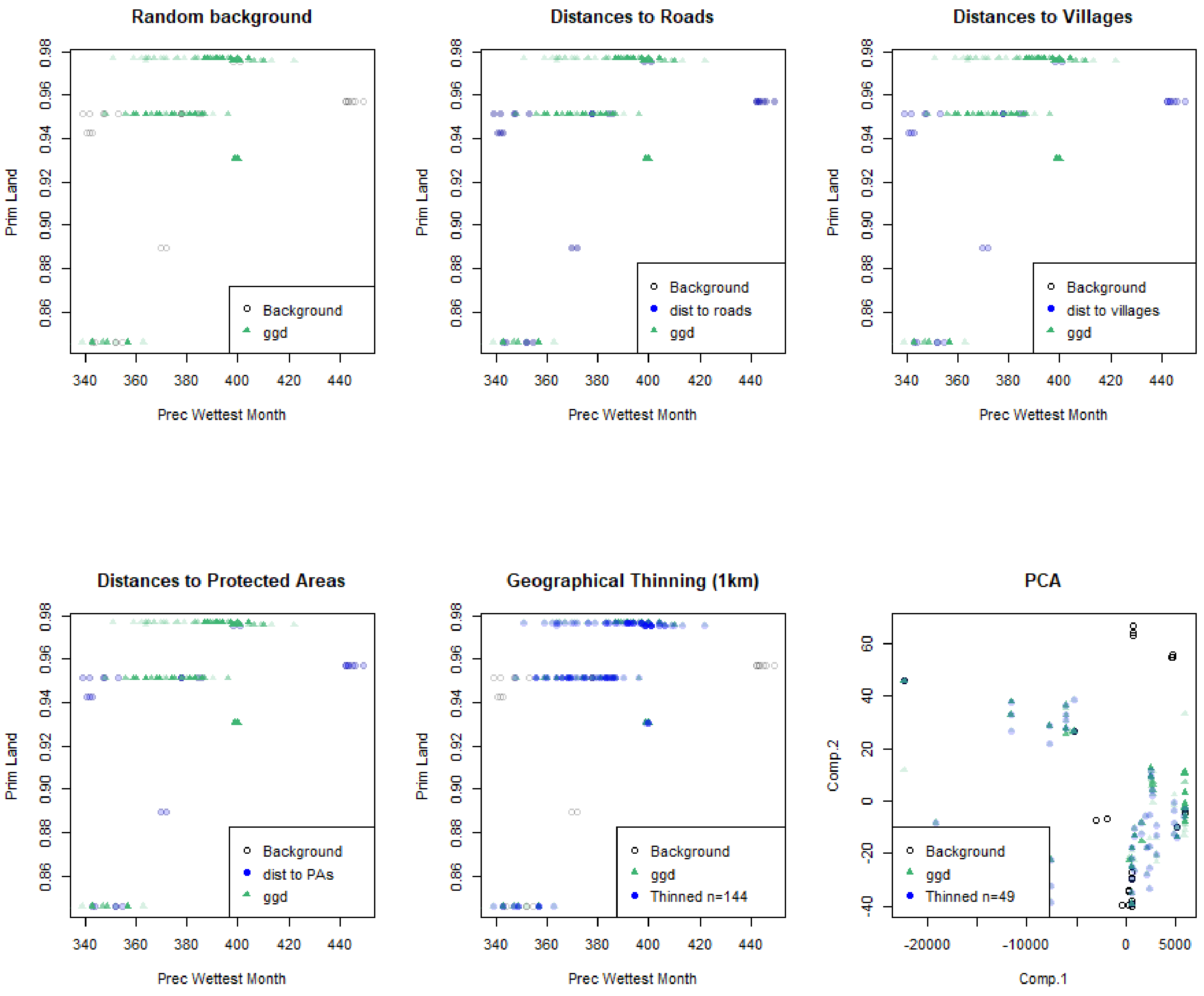

*Gorilla gorilla gorilla*

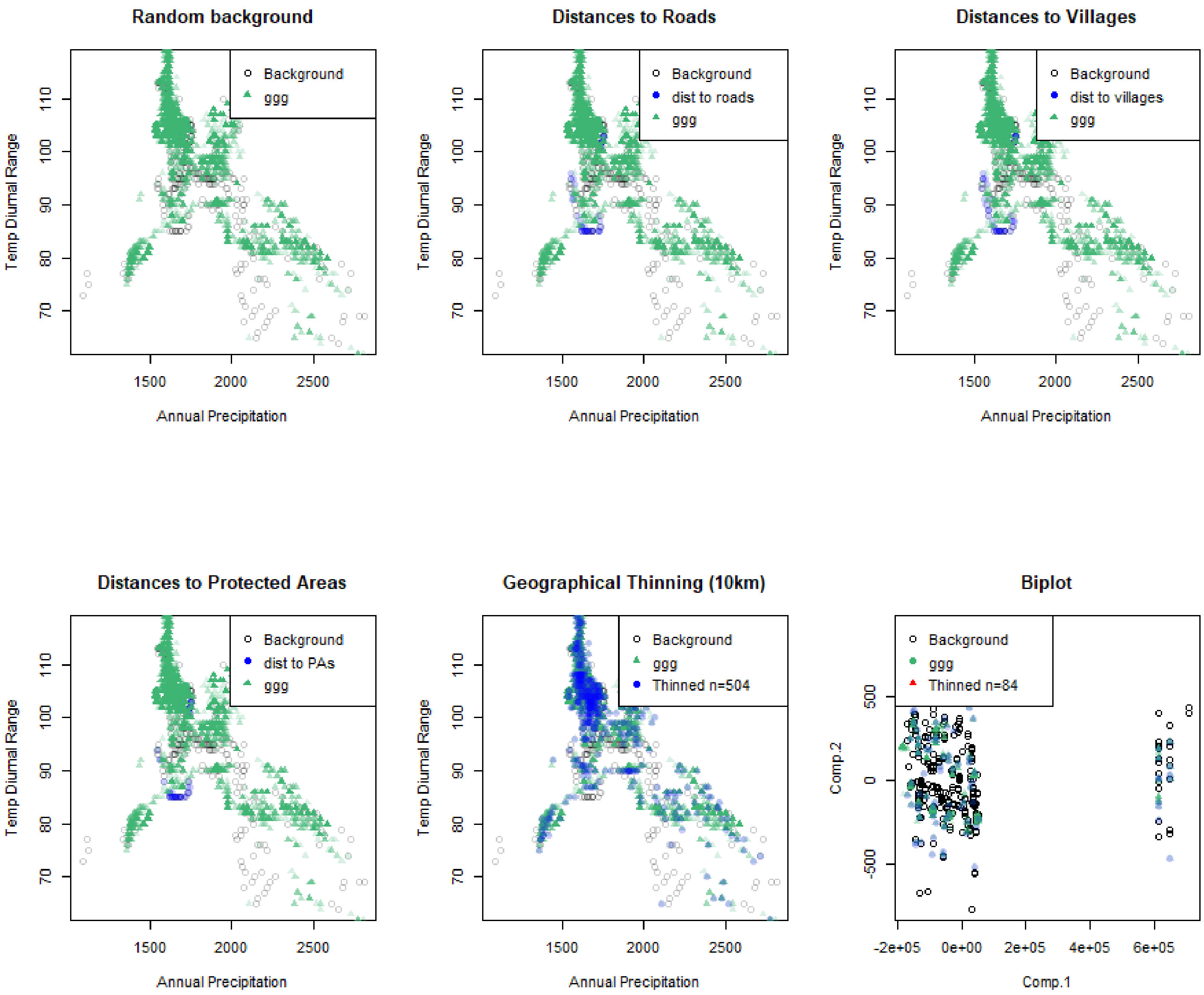

*Pan paniscus*

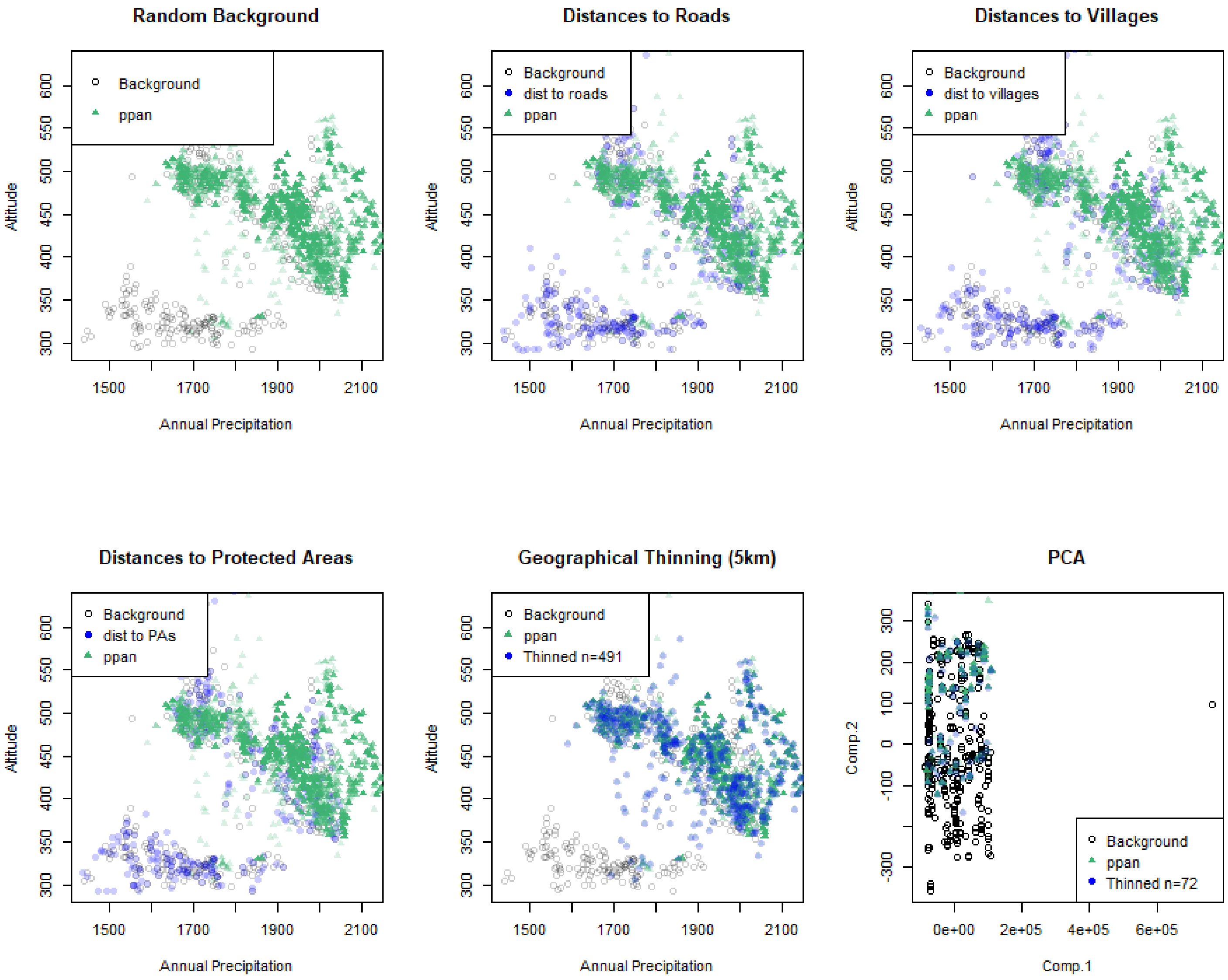

*Pan troglodytes ellioti*

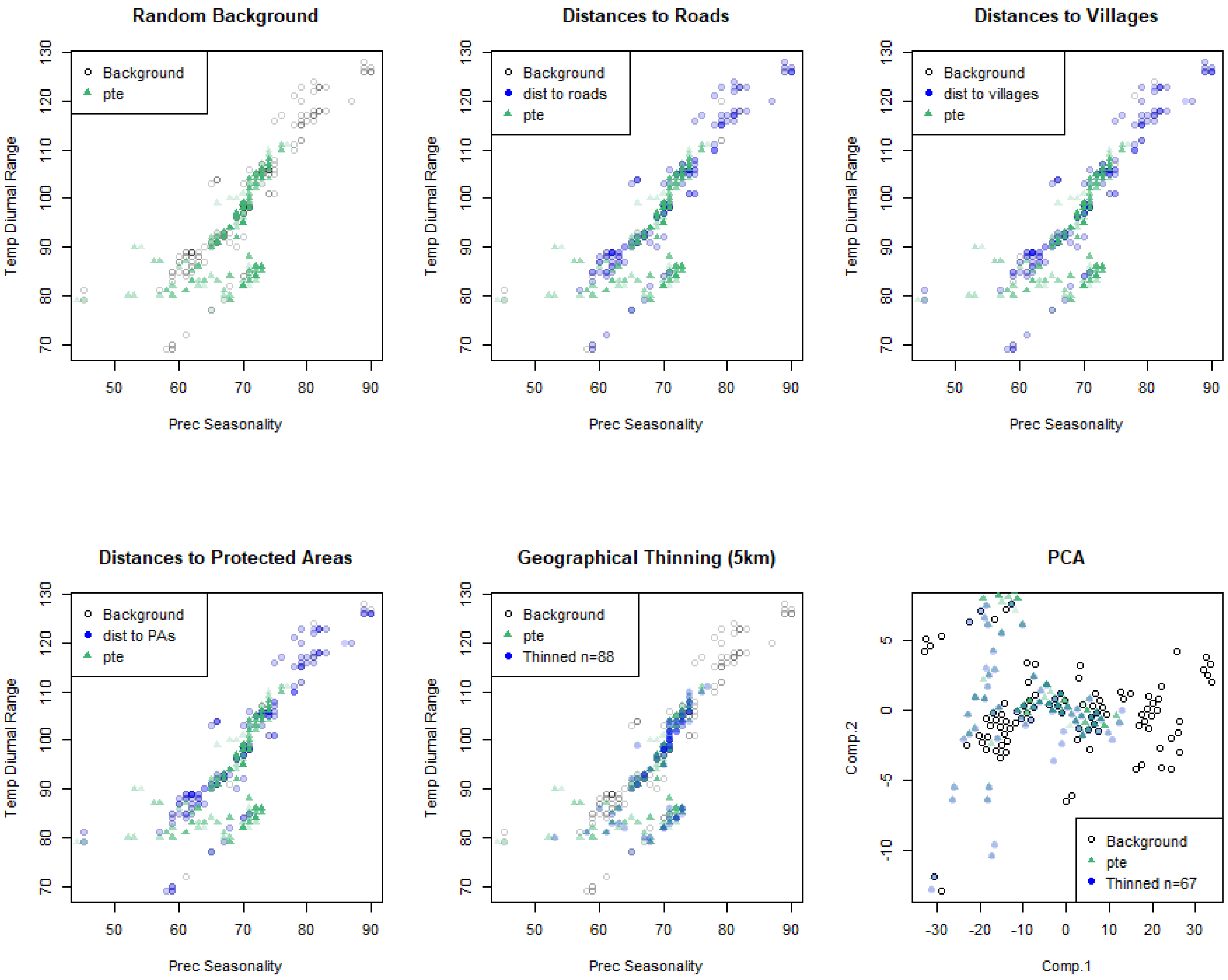

*Pan troglodytes schweinfurthii*

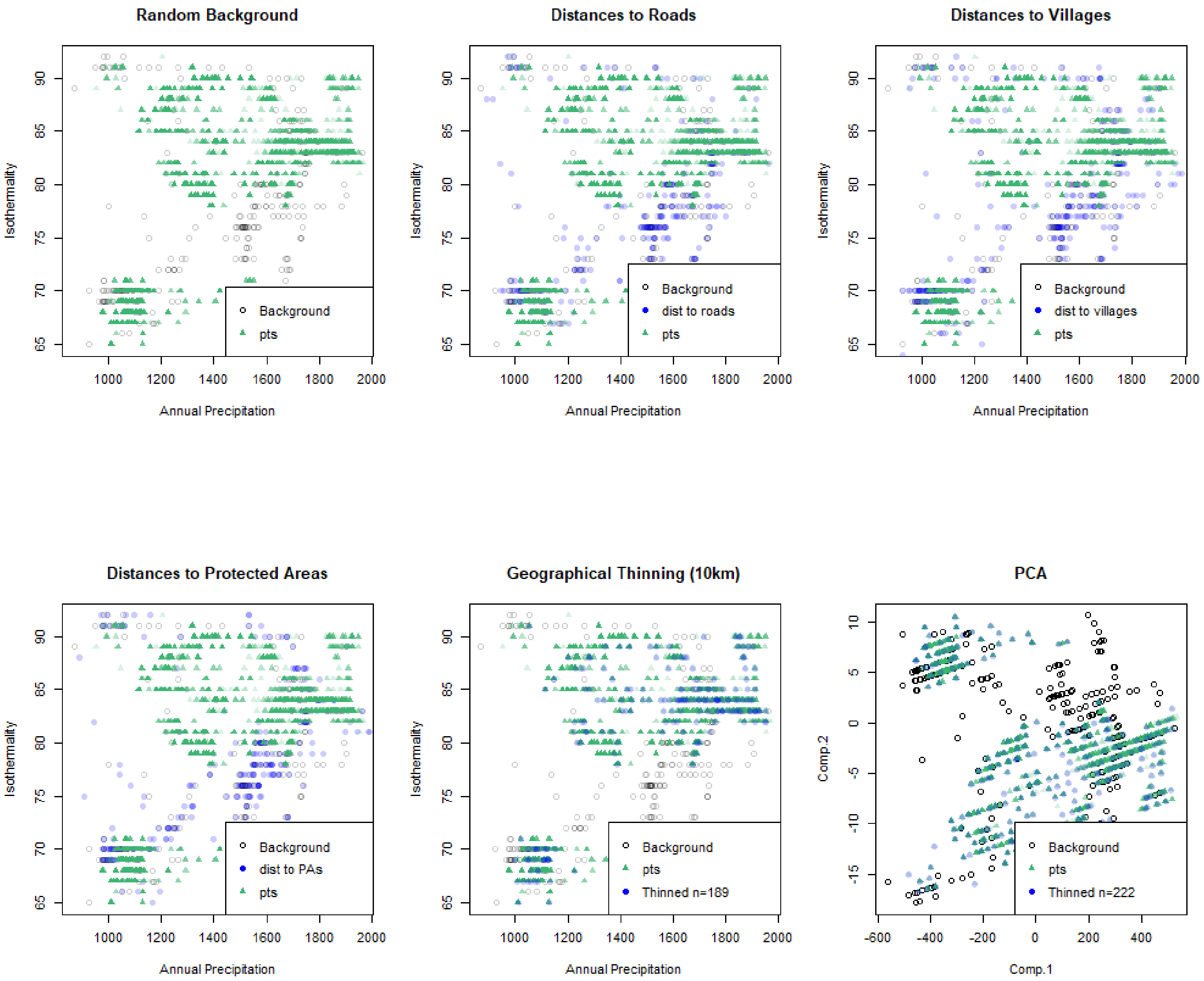

*Pan troglodytes troglodytes*

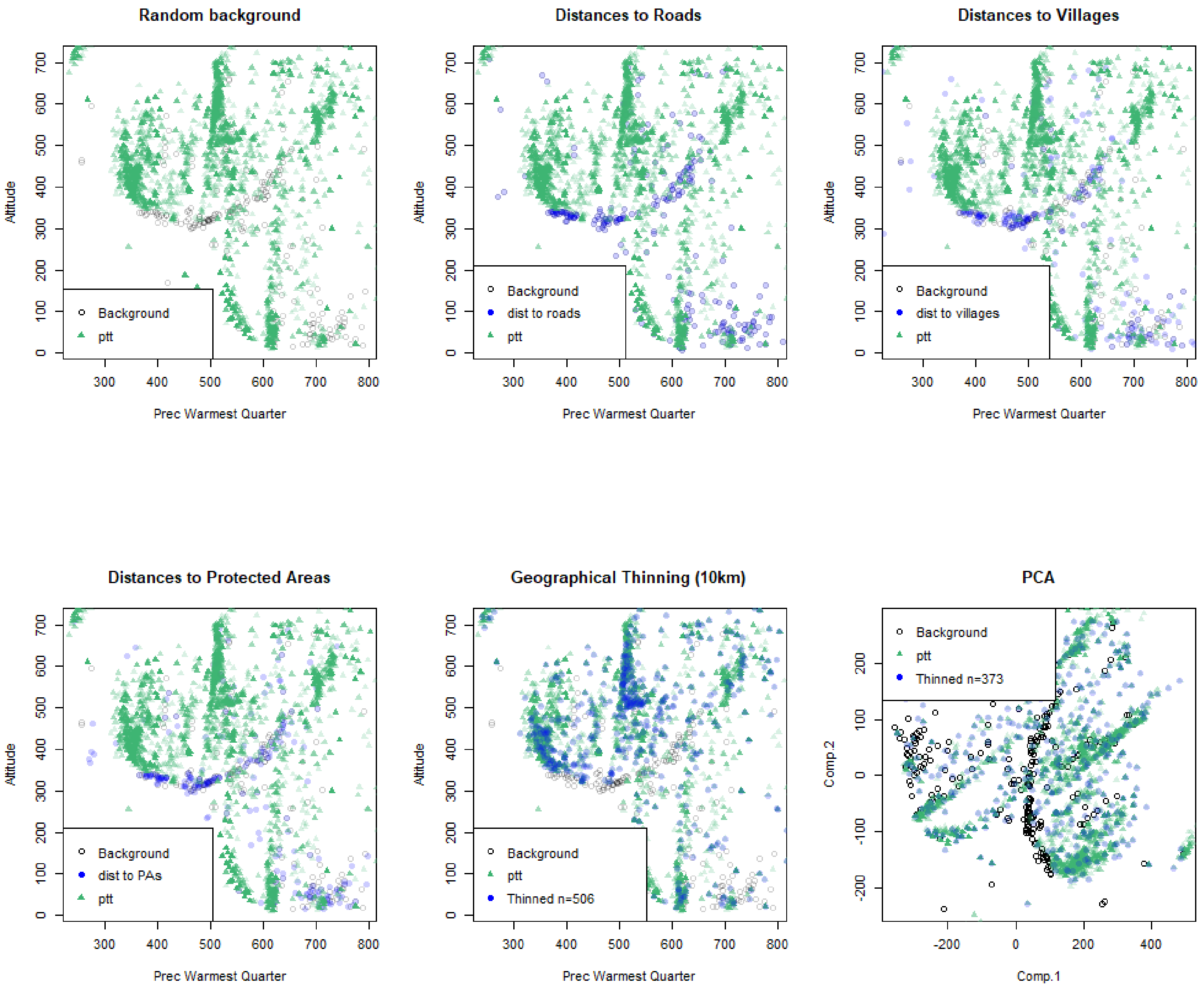

*Pan troglodytes verus*

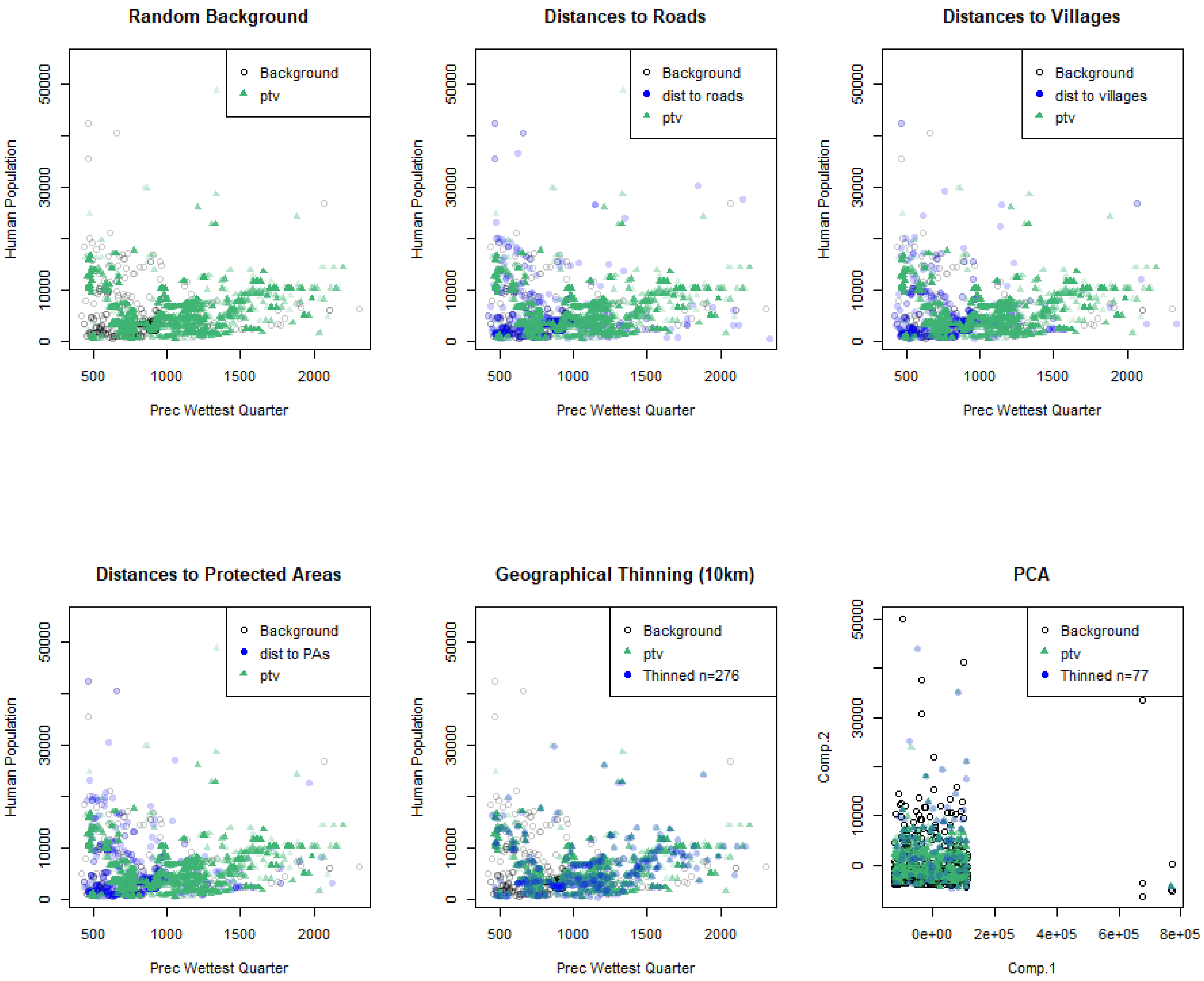


Figure S1.1. Plots of the sampling bias based on different methods of bias correction for each taxon. For each taxon, only two of the most important predictors are plotted (see Fig. 1, S4) and the best approach to sampling bias correction was selected based on the greatest overlap between the predictor values of sampling bias over the species occurrences. Given that geographic thinning approach performed best for all taxa, it was integrated into the SDMs for sampling bias correction.

White circles – random background data, blue circles – sampling bias data, and green triangles – taxon evidences. See methods section for more details and Table S1 for variable predictor names. See more examples at https://drive.google.com/file/d/14XLhdZKEx0OPN-_ACWNxAtEY_4AWZhPN/view.

*Gorilla beringei beringei* (gbb)

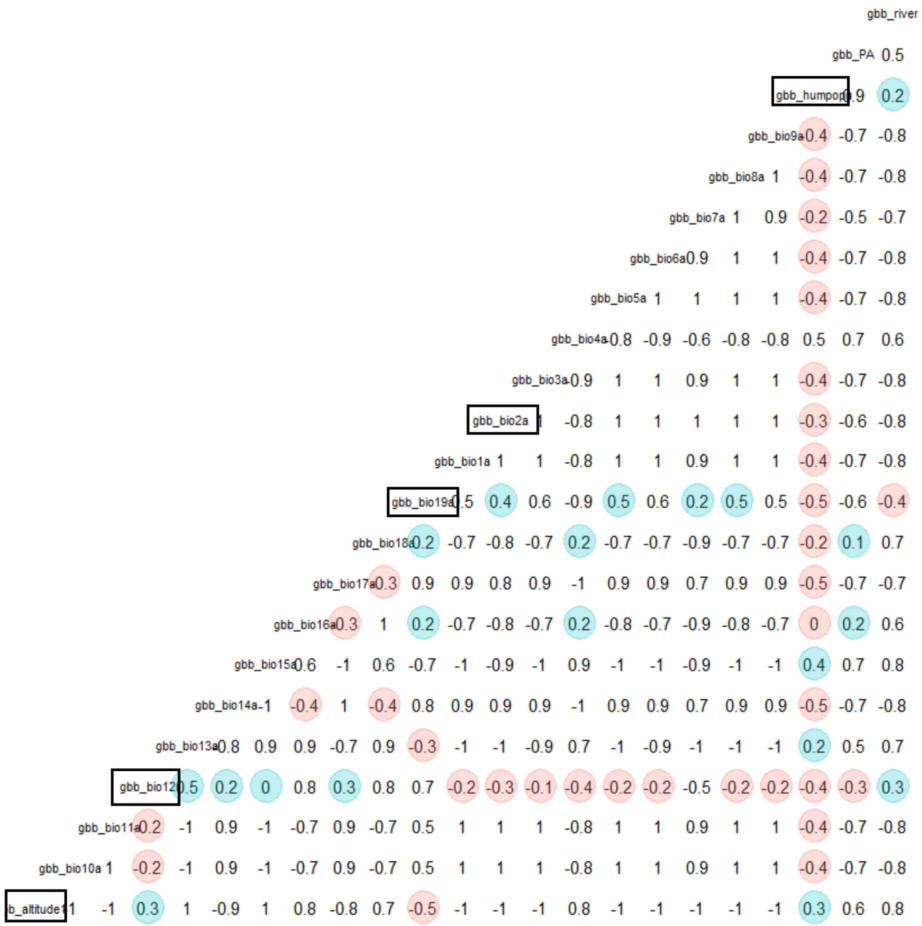


*Gorilla beringei graueri* (gbg)

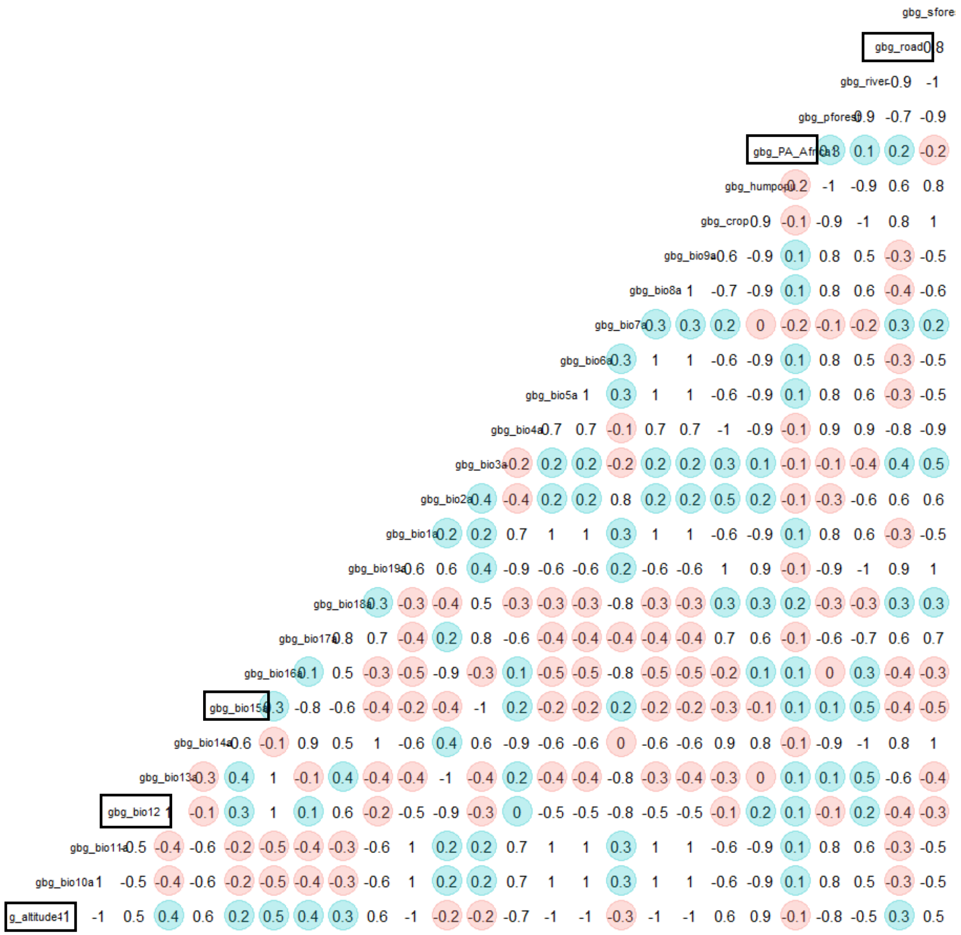

### *Gorilla gorilla diehli* (ggd)

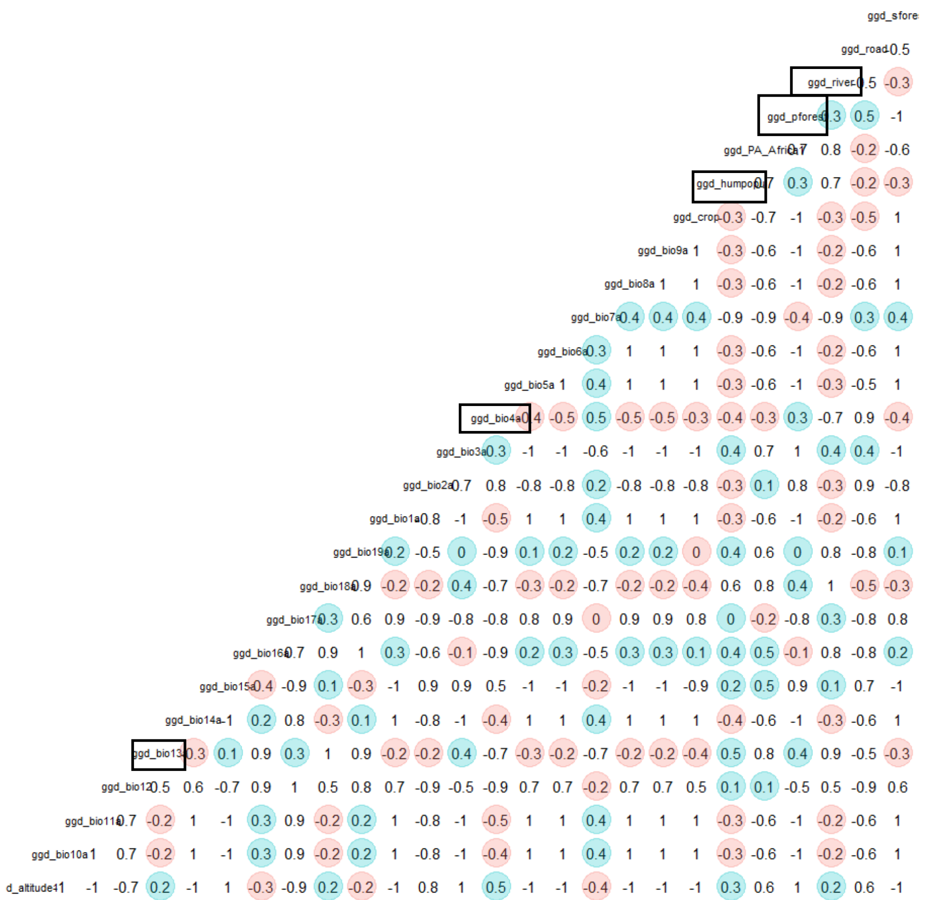


### *Gorilla gorilla gorilla* (ggg)

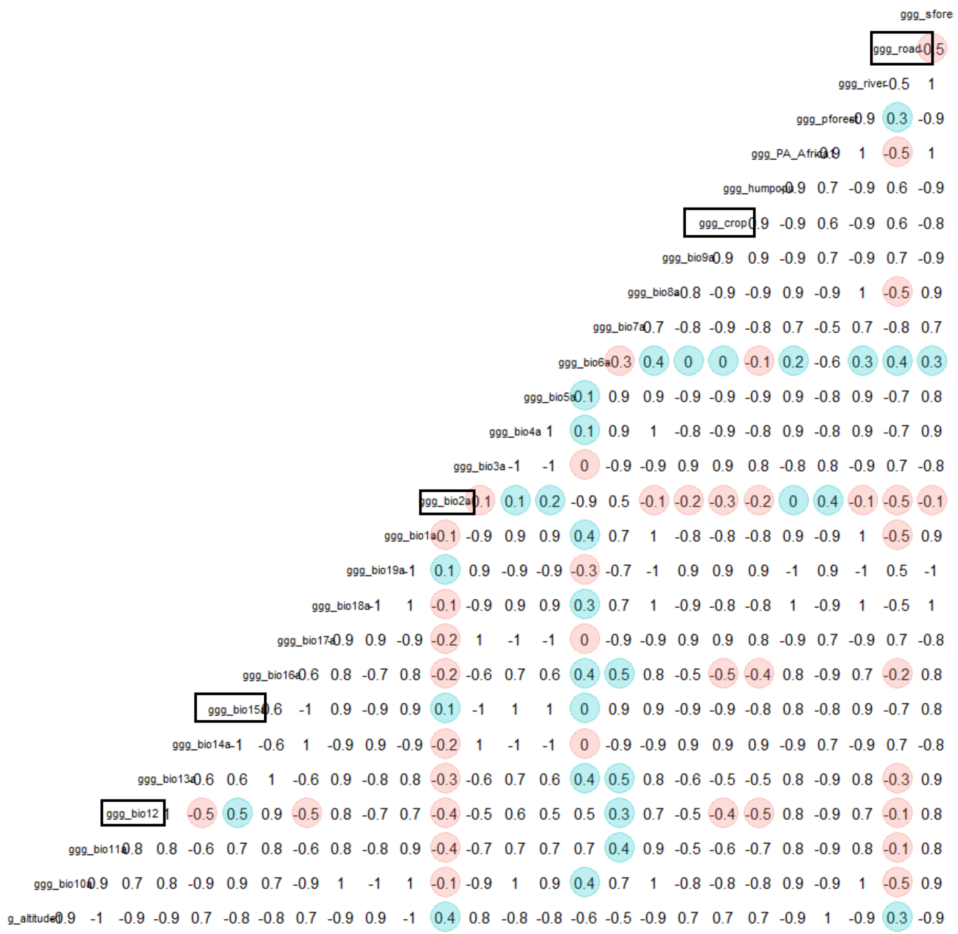

*Pan paniscus* (ppan)

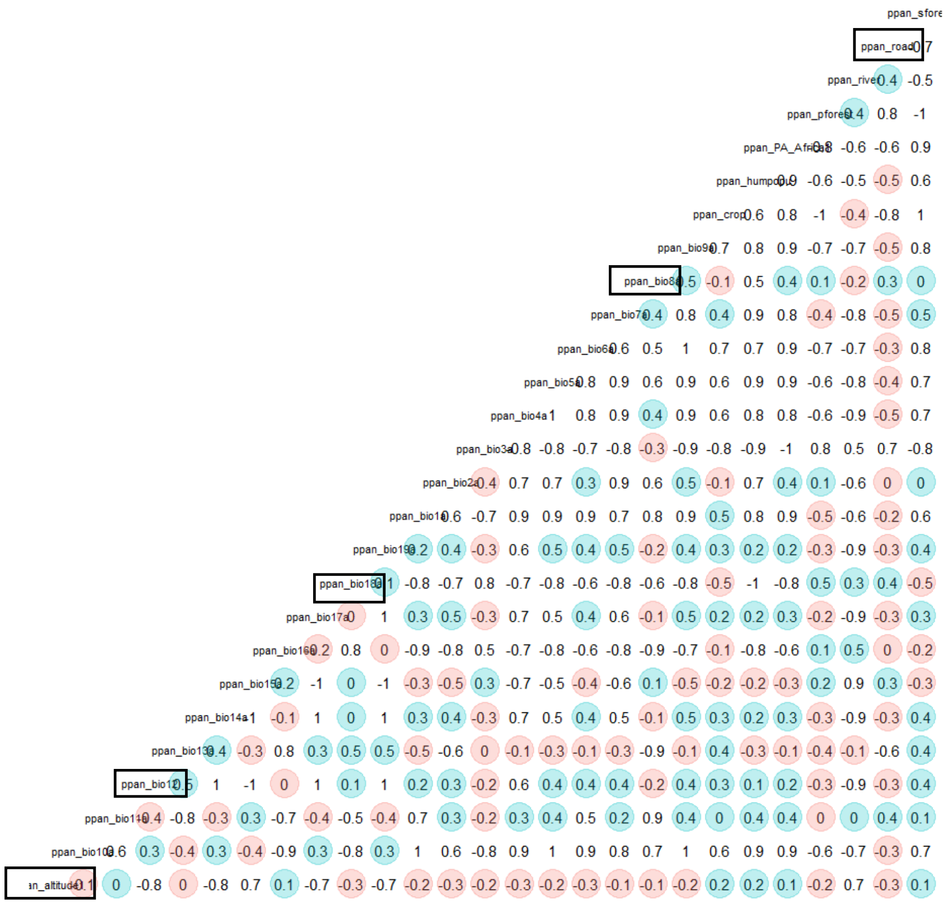

*Pan troglodytes ellioti* (pte)

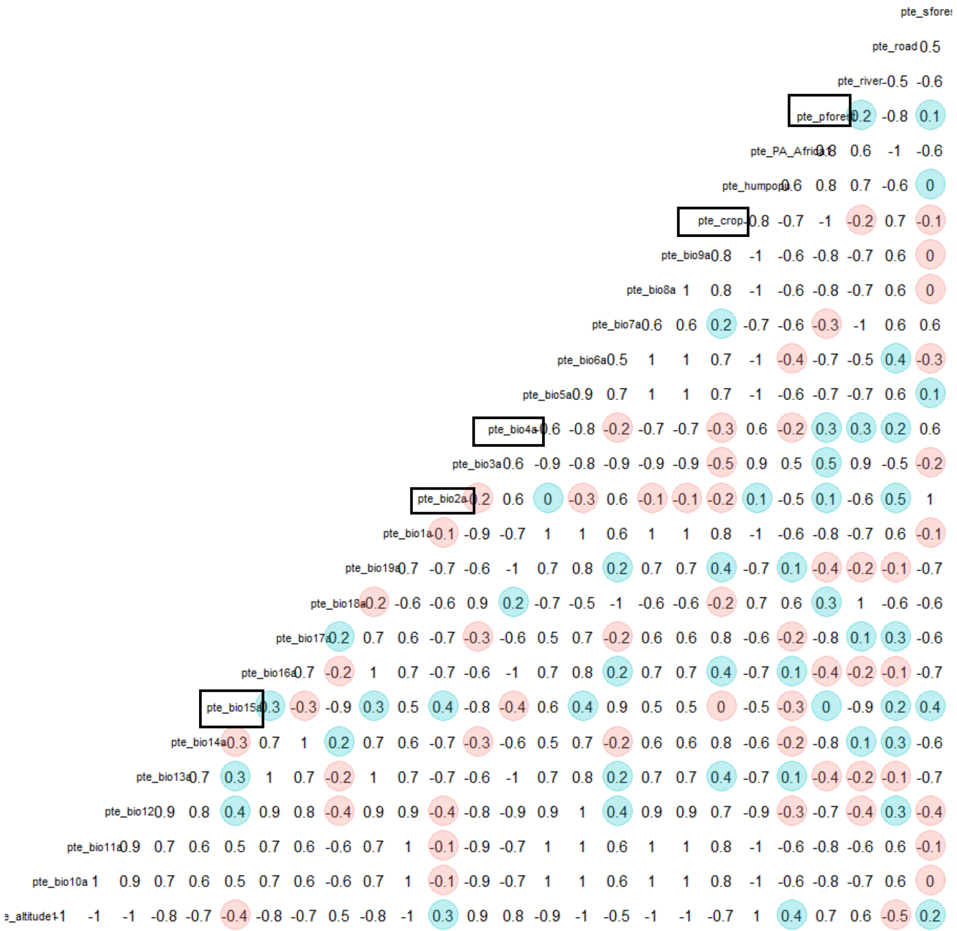

*Pan troglodytes schweinfurthii* (pts)

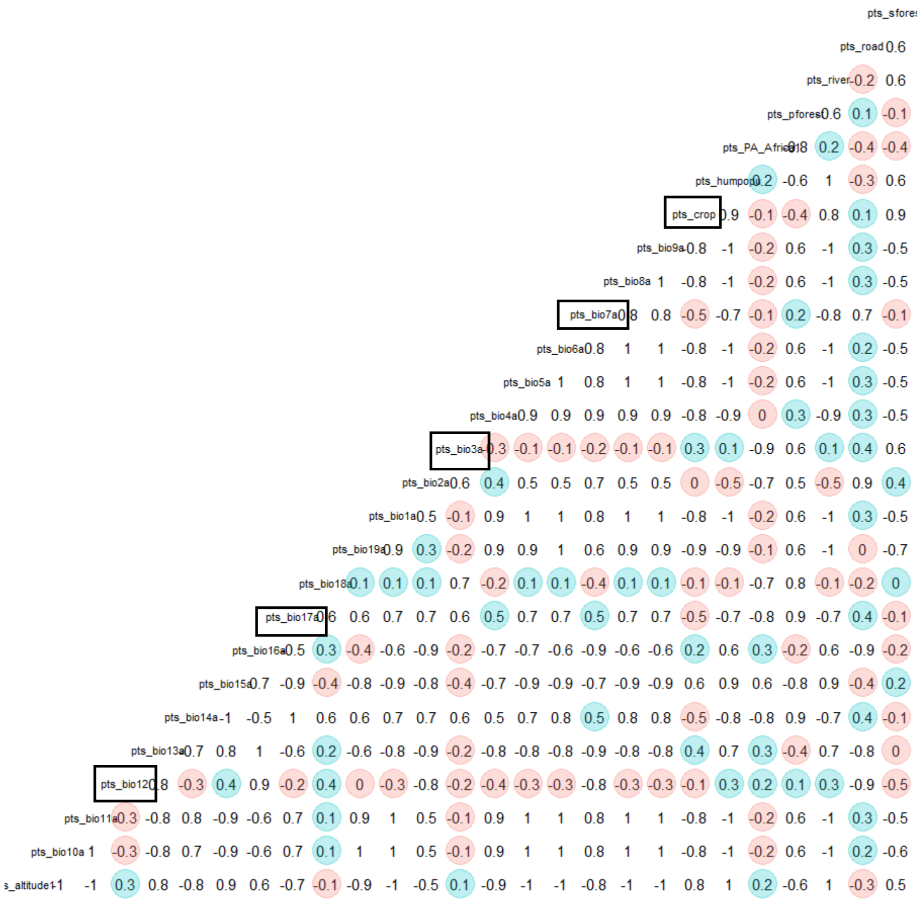


*Pan troglodytes troglodytes* (ptt)

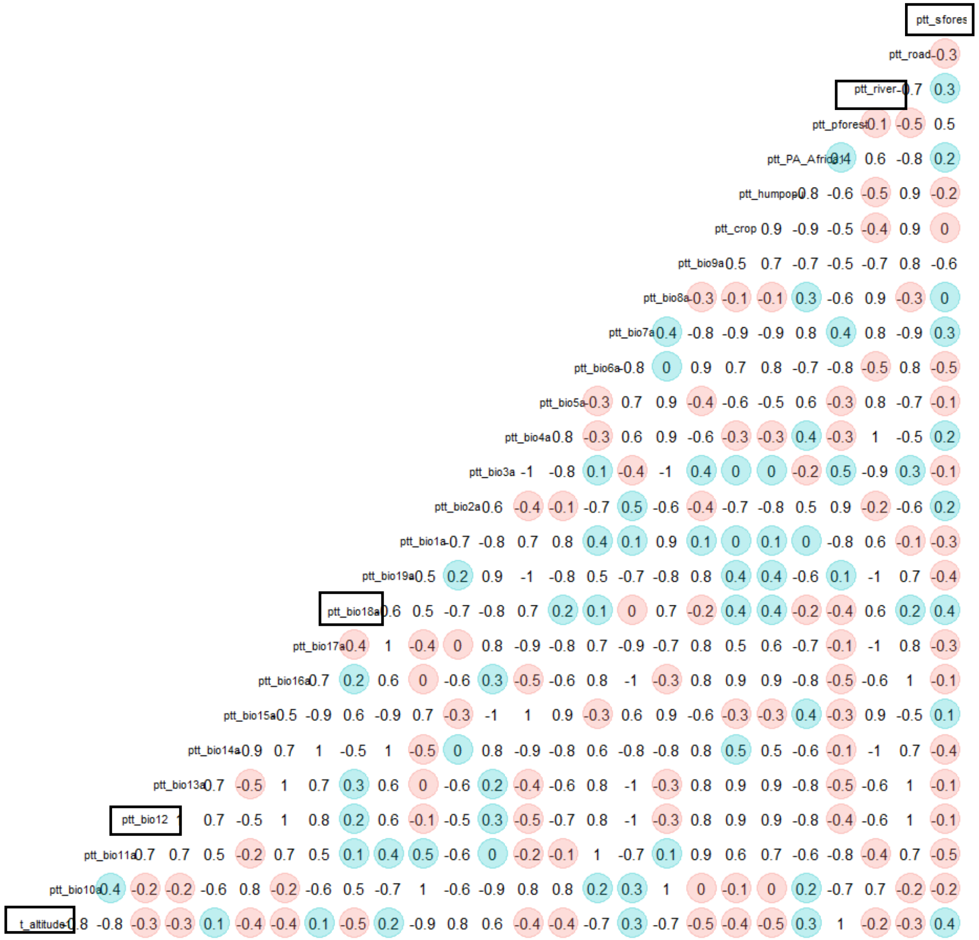

*Pan troglodytes verus* (ptv)

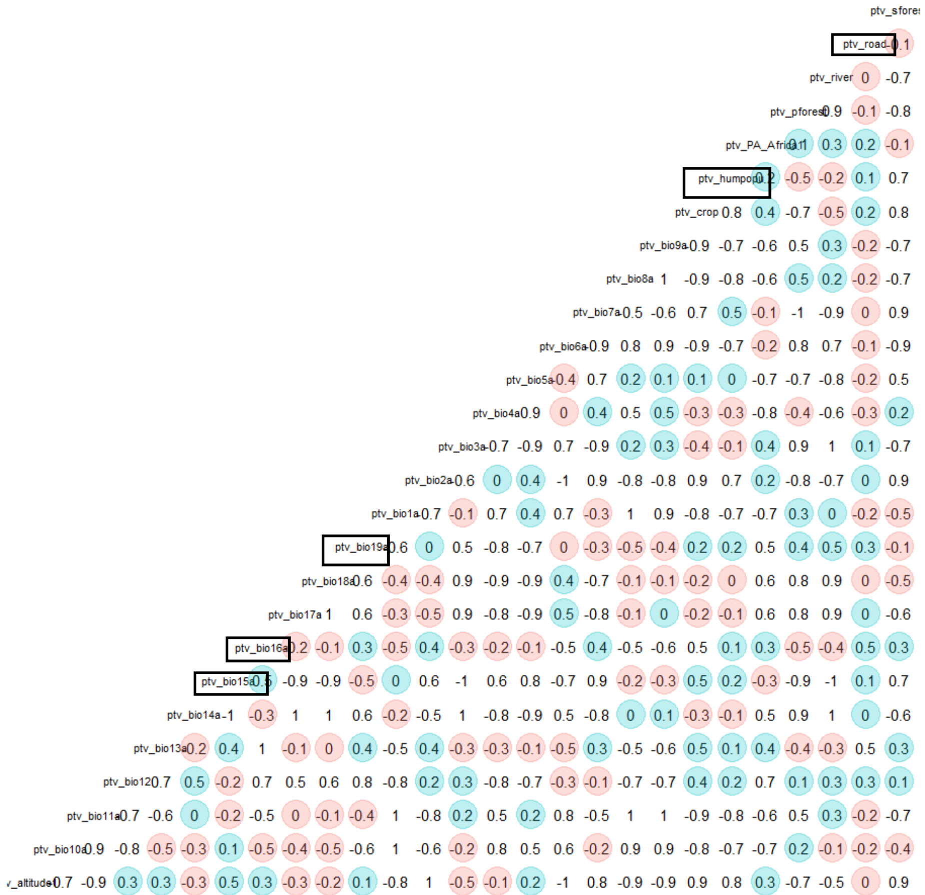


List of abbreviations:

bio1a – annual mean temperature
bio2a – mean diurnal range
bio3a – isothermality
bio4a – temperature seasonality
bio5a – maximum temperature of warmest month
bio6a – minimum temperature of coldest month
bio7a – temperature annual range
bio8a – temperature of wettest quarter
bio9a – mean temperature of driest quarter
bio10a – temperature of warmest quarter
bio11a – temperature of the coldest quarter
bio12a – annual precipitation
bio13a – precipitation of wettest month
bio14a – precipitation of driest month
bio15a – Seasonal variation of precipitation
bio16a – precipitation of wettest quarter
bio17a – precipitation of driest quarter
bio18a – precipitation of warmest quarter
bio19a – precipitation of coldest quarter
crop – cropland
humpopu – human population
PA_Africa – distances to protected areas
Pforest – primary land
River – distances to rivers
Road – distances to roads
Sforest – secondary land

Figure S1.2. Results of Spearman correlations between all predictor variables for each taxon. Only pairwise correlations less than 0.5 are highlighted (blue – positive correlations, red – negative correlations) to aid in the selection of a subset of five least correlated variables (black square). For this, variables were ranked based on the number of pairwise correlations <0.5, selecting those with the highest number as being least correlated.

a)

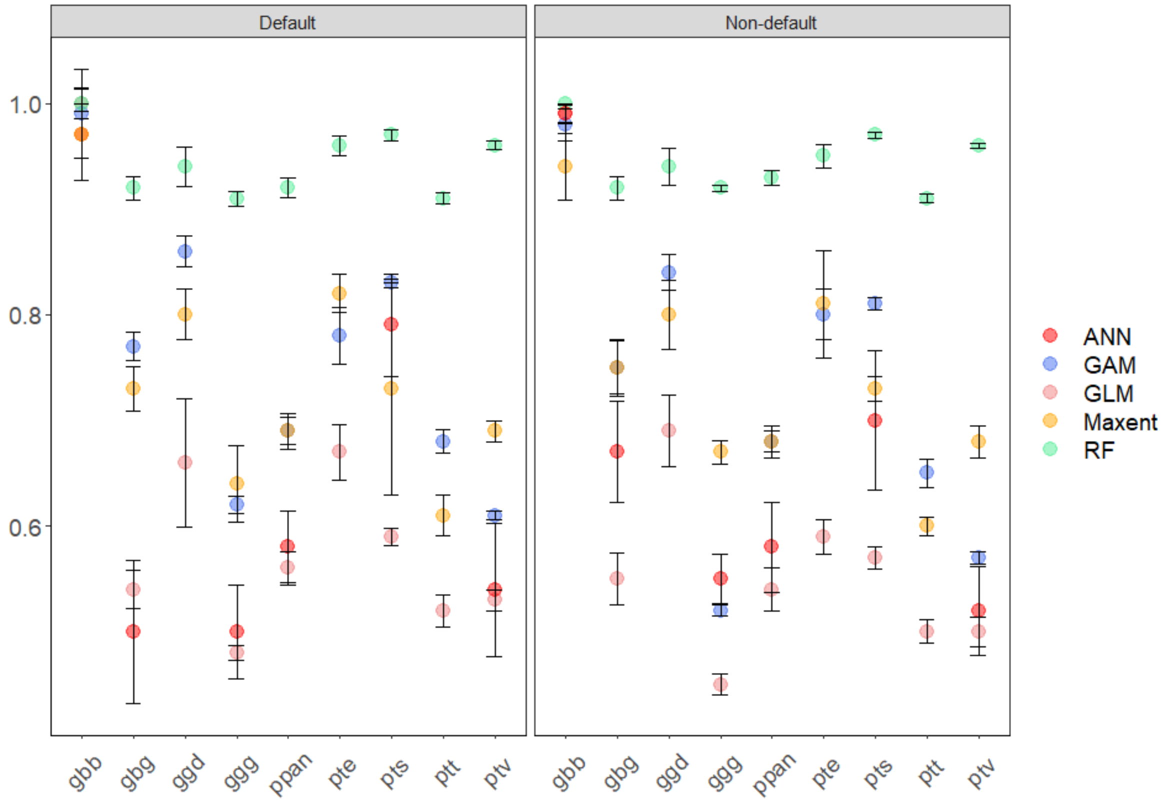


b)

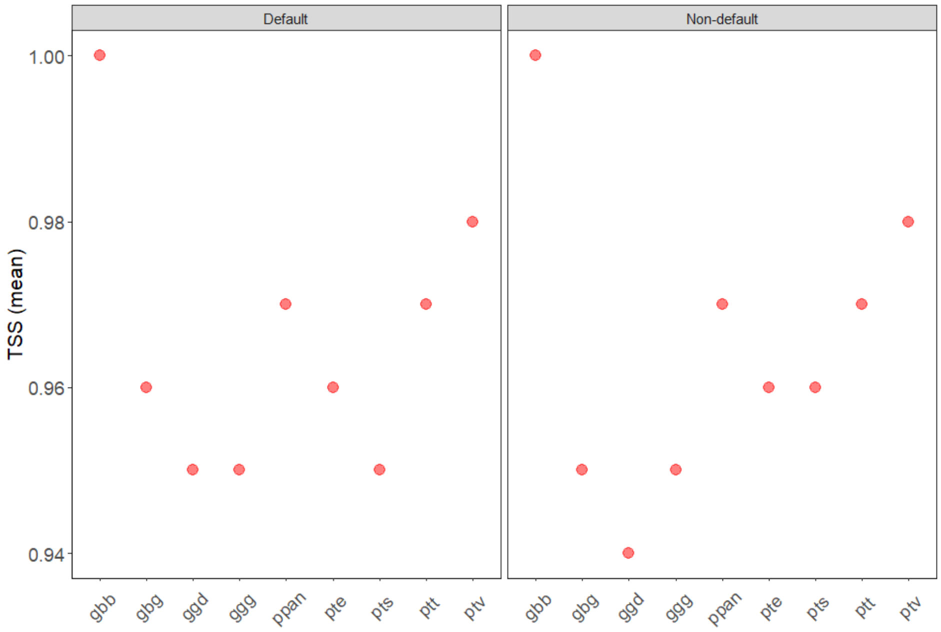


Figure S1.3. Predictive accuracy of individual (a) and ensemble (b) models for both modelling approaches. The mean TSS values and respective standard deviation (SD) are provided for each algorithm evaluated.

a)

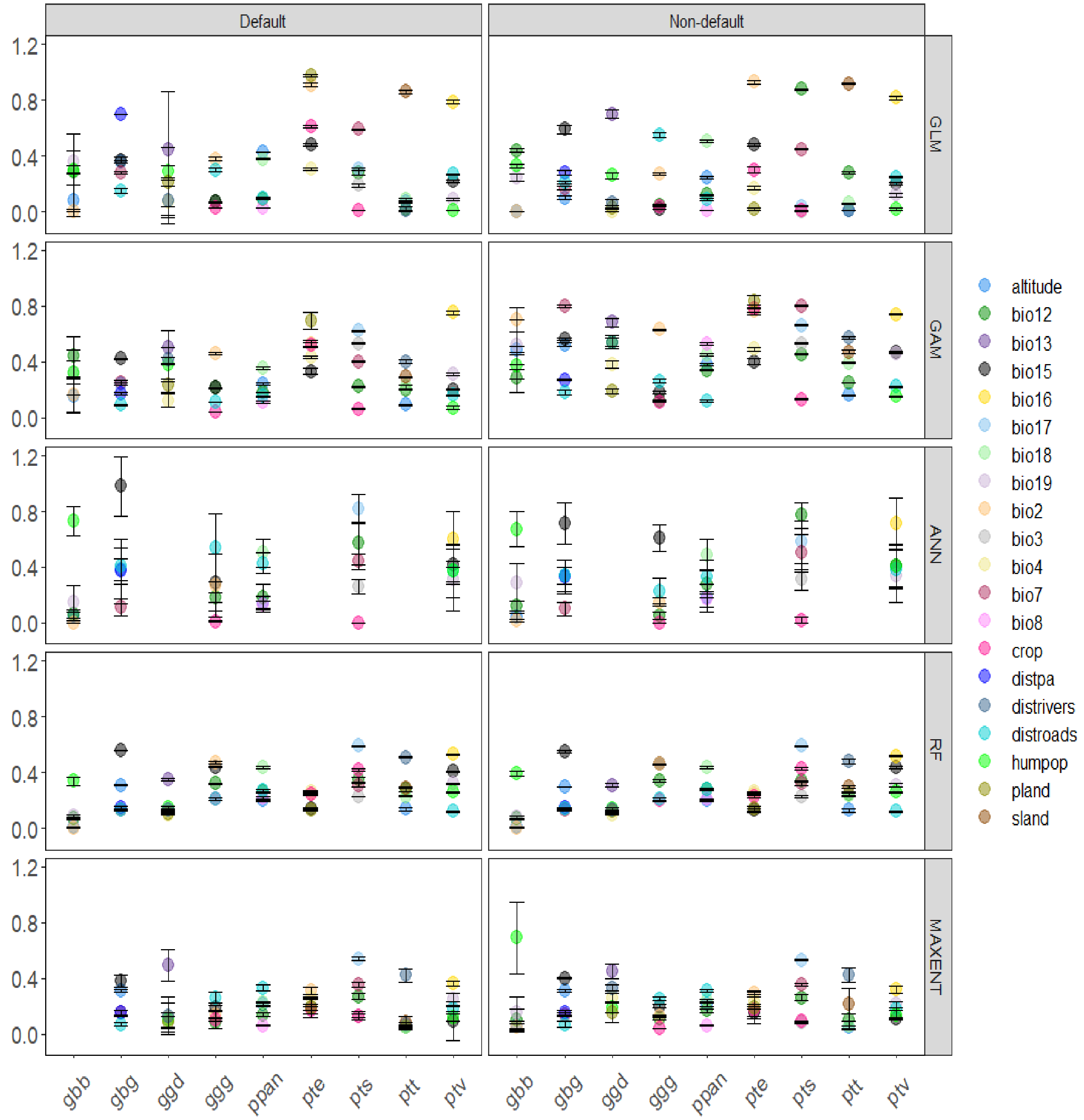
Default
Non-default
GLM
GAM
ANN
RF
MAXENT
1.2
0.8
0.4
0.0
gbb
gbg
ggd
ggg
ppan
pte
pts
ptt
ptv
altitude
bio12
bio13
bio15
bio16
bio17
bio18
bio19
bio2
bio3
bio4
bio7
bio8
crop
distpa
distrivers
distroads
humpop
pland
sland

b)

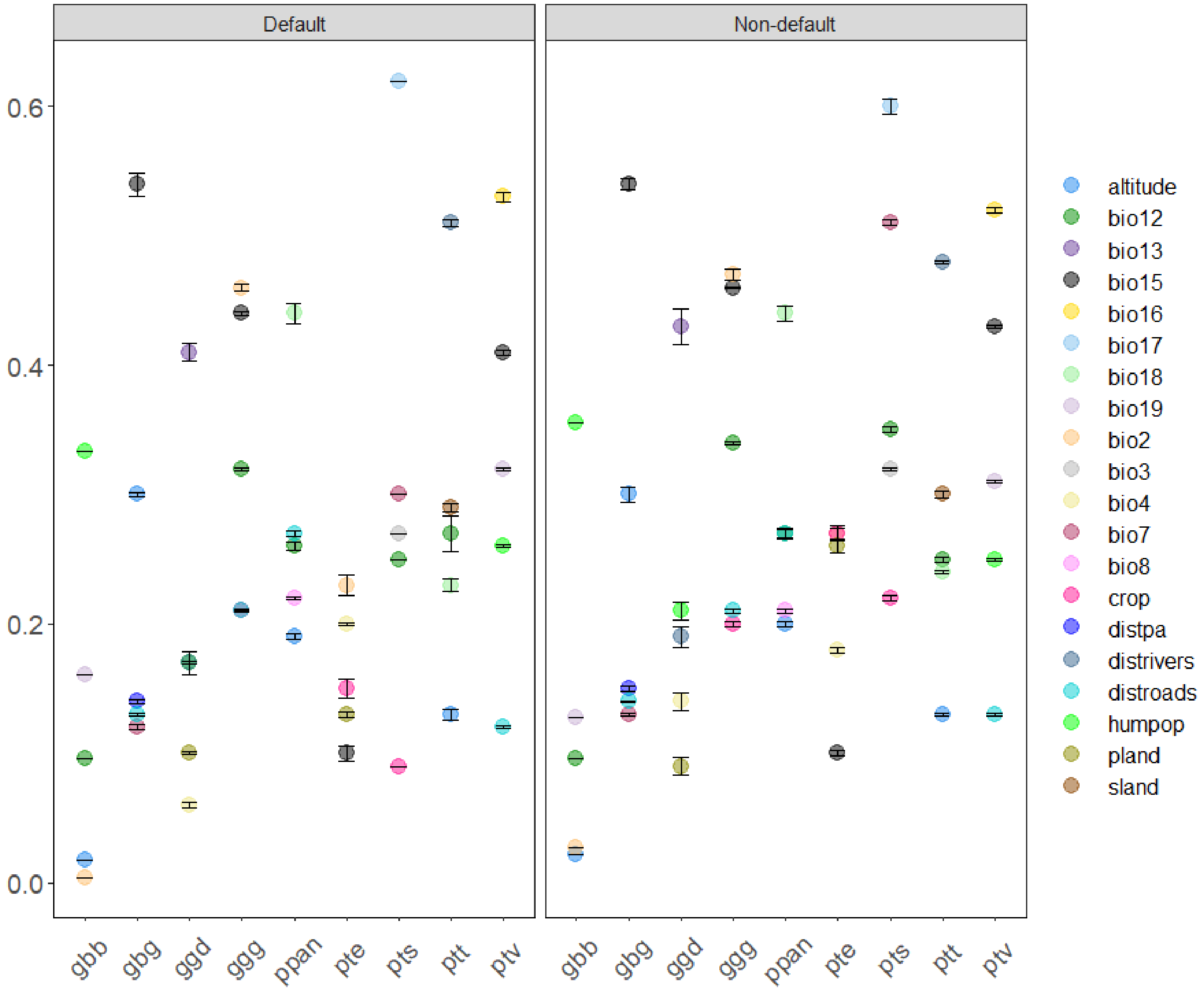


Figure S1.4. Variable importance (mean and standard deviation (SD) of the correlation values) for the individual (a) and ensemble (b) models given two modelling approaches. For the ensemble models, only models with TSS>0.8 were considered and scores of the evaluation metric weighted mean are given. Variable predictor abbreviations: bio12 – annual precipitation, bio13 – precipitation of wettest month, bio15 – Seasonal variation of precipitation, bio16 – precipitation of wettest quarter, bio17 – precipitation of driest quarter, bio18 – precipitation of warmest quarter, bio19 – precipitation of coldest quarter, bio2 – mean diurnal range, bio3 – isothermality, bio4 – temperature seasonality, bio7 – temperature annual range, bio8 – mean temperature of wettest quarter, crop – cropland, distpa – distance to protected areas, distrivers – distances to rivers, distroads– distances to roads, humpop – human population density, pland– primary land, sland – secondary land.

*Gorilla beringei beringei*

### Current

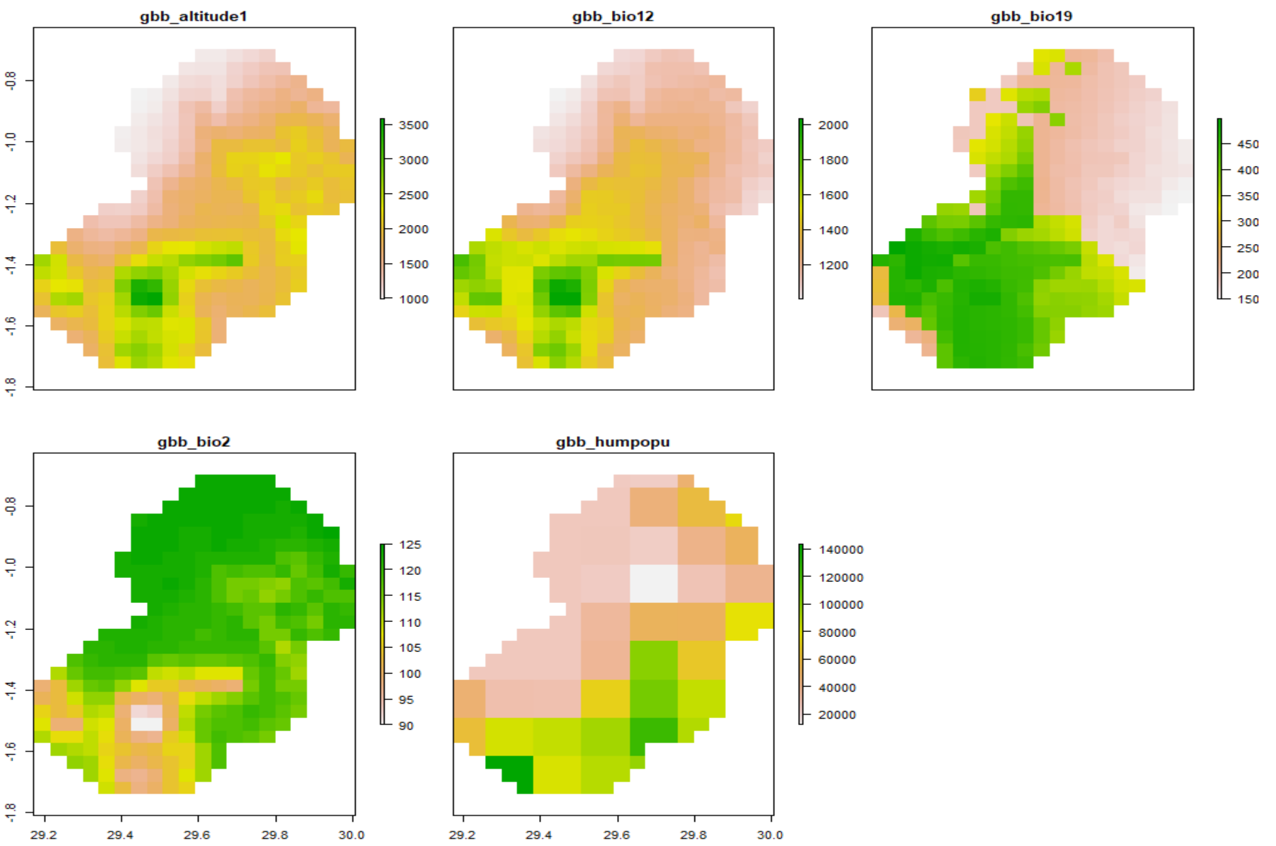


### Best scenario

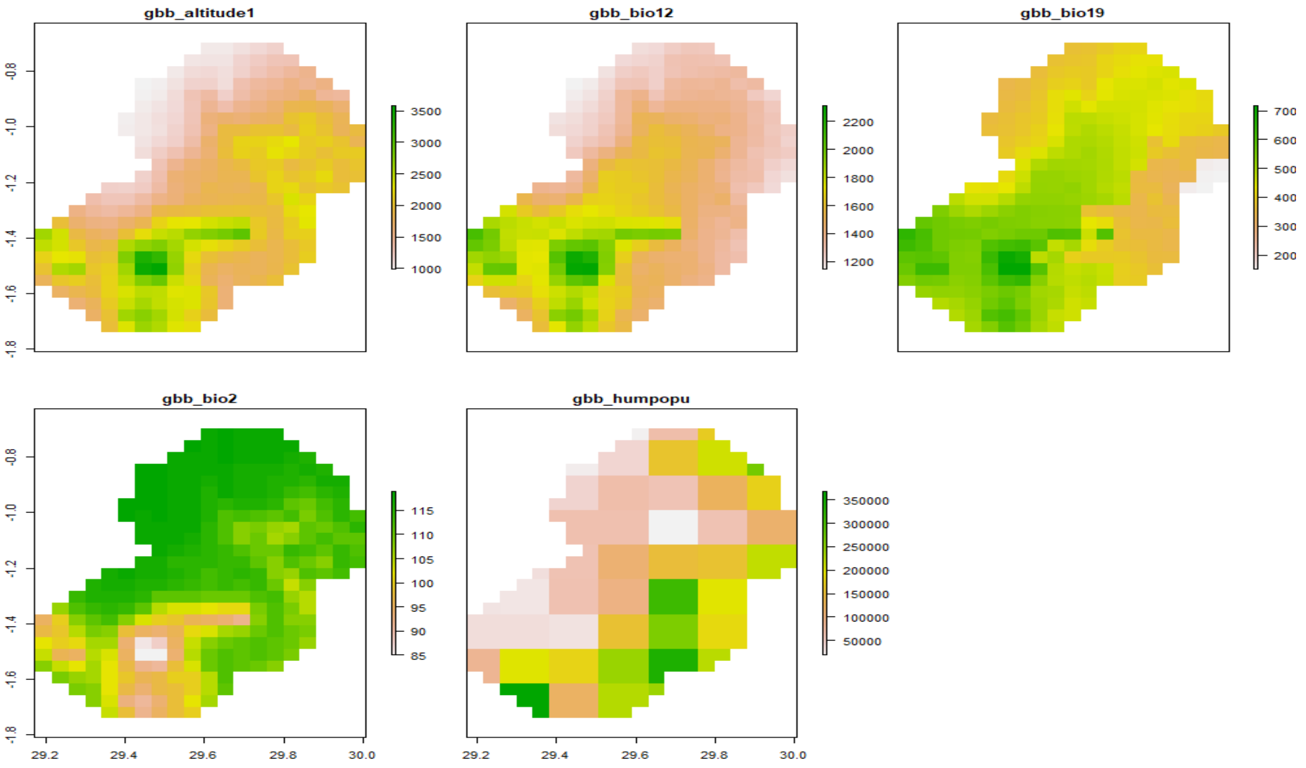

### Worst scenario

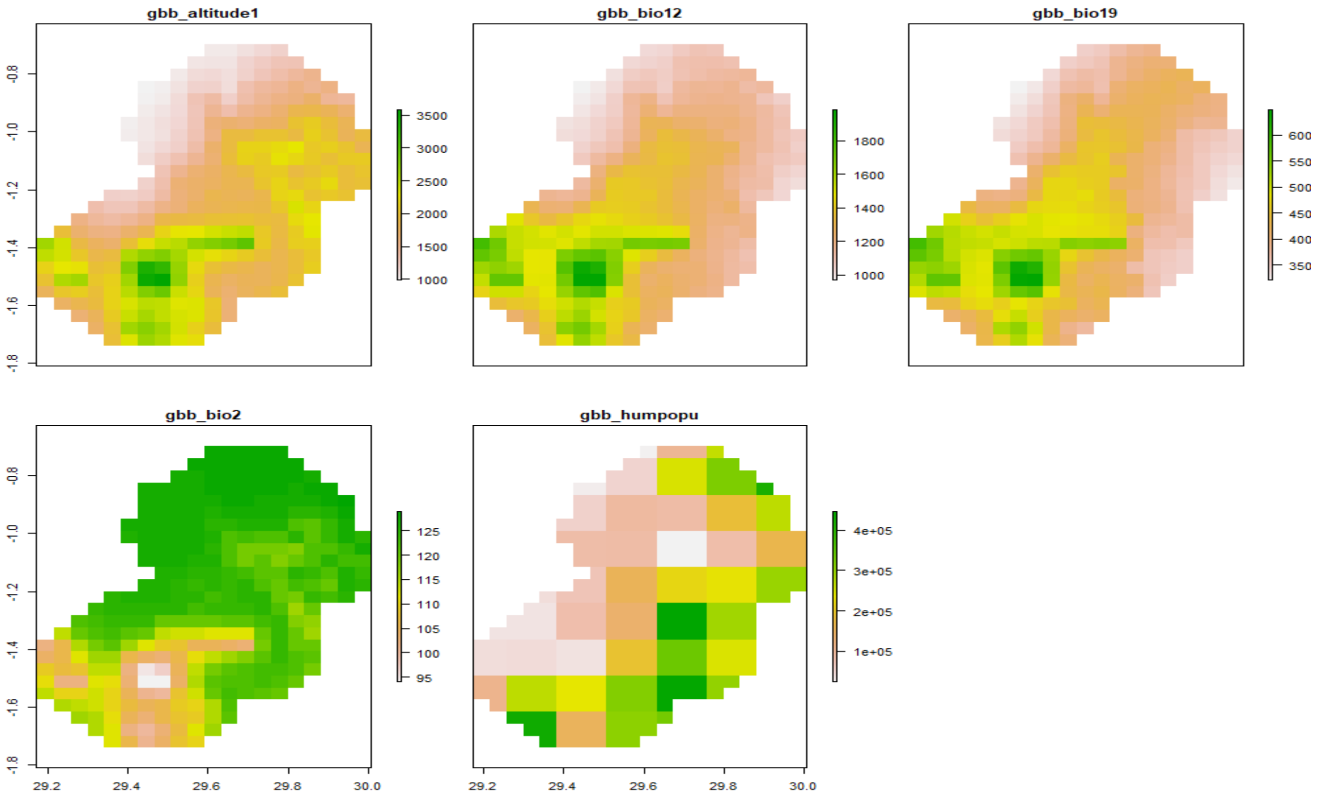


*Gorilla beringei graueri*

### Current

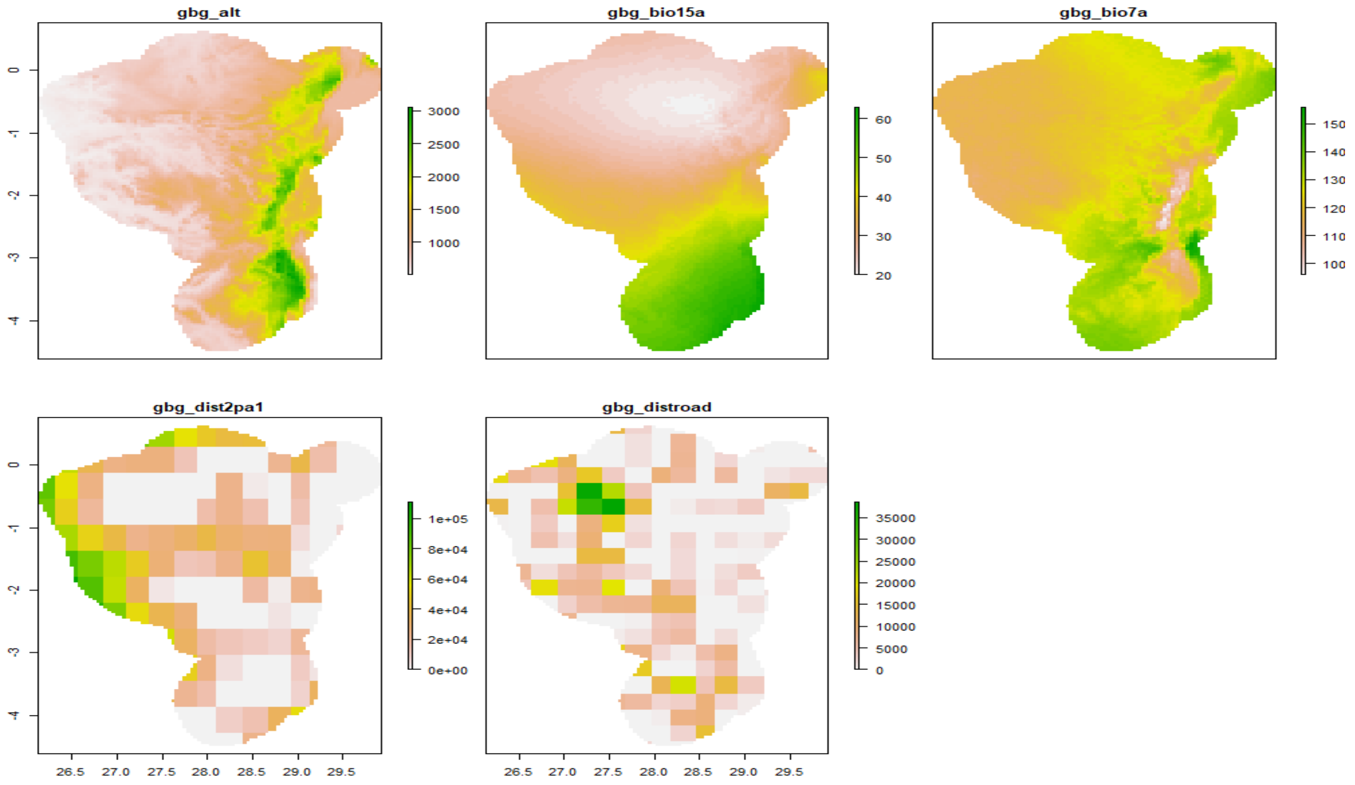

### Best scenario

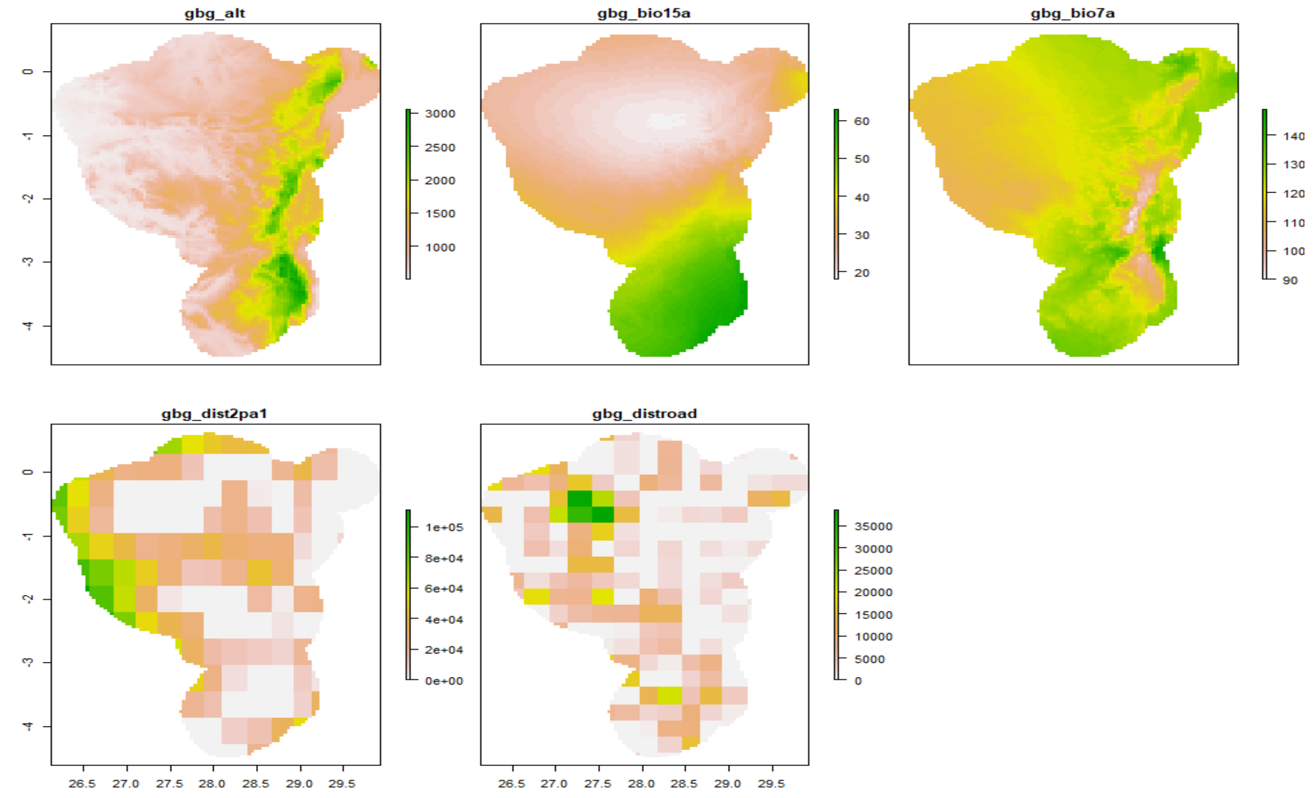


### Worst scenario

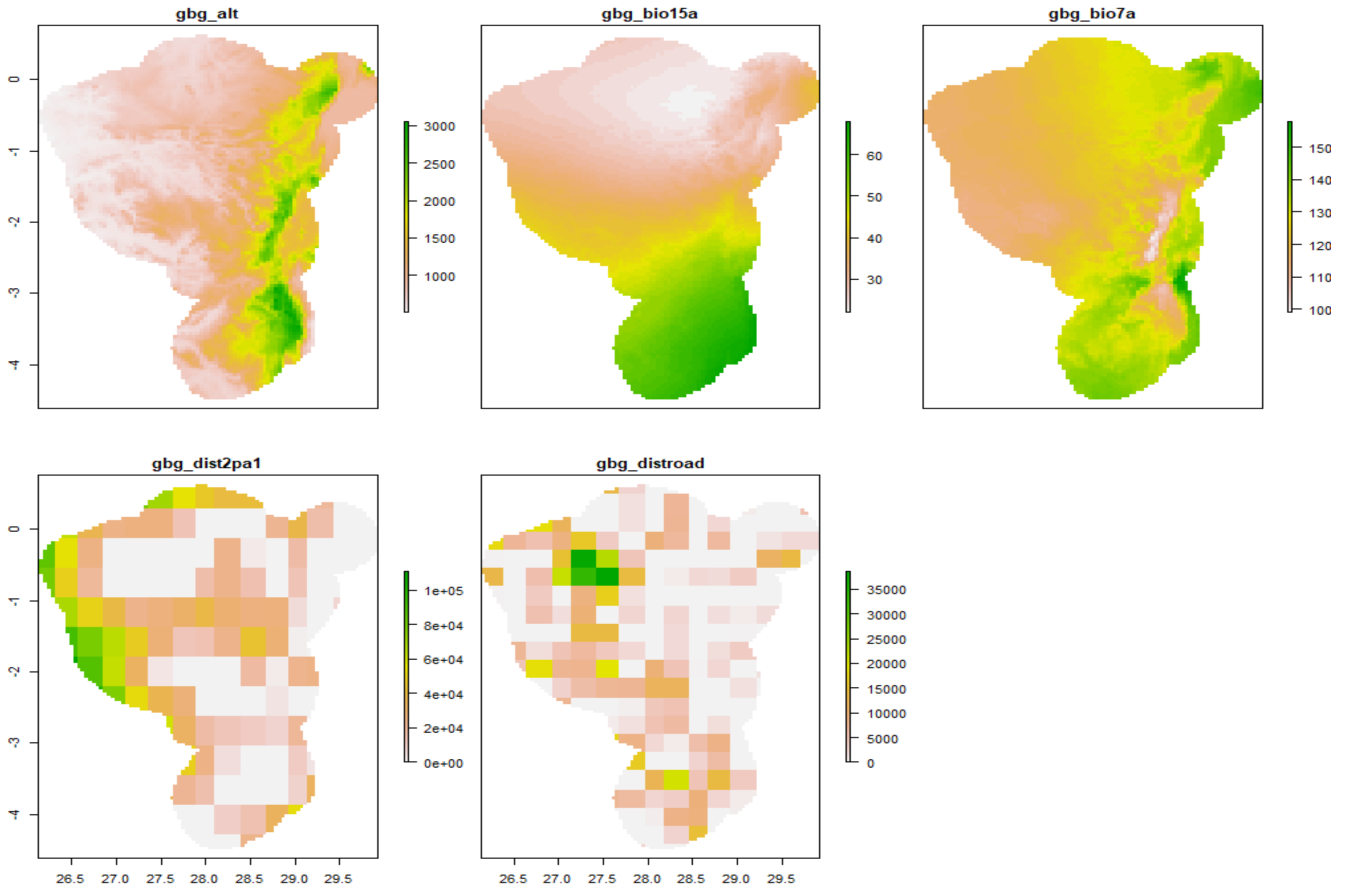

*Gorilla gorilla diehli*

**Current**

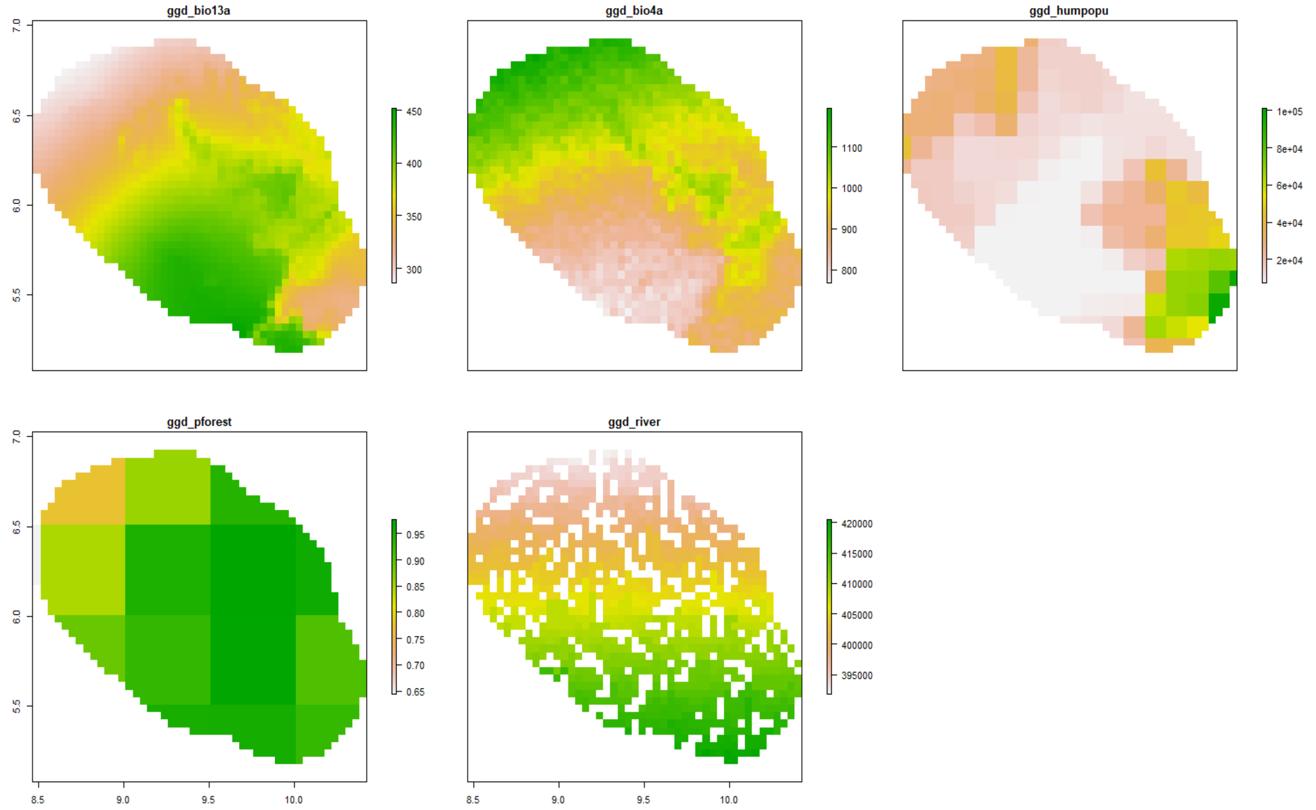


**Best scenario**

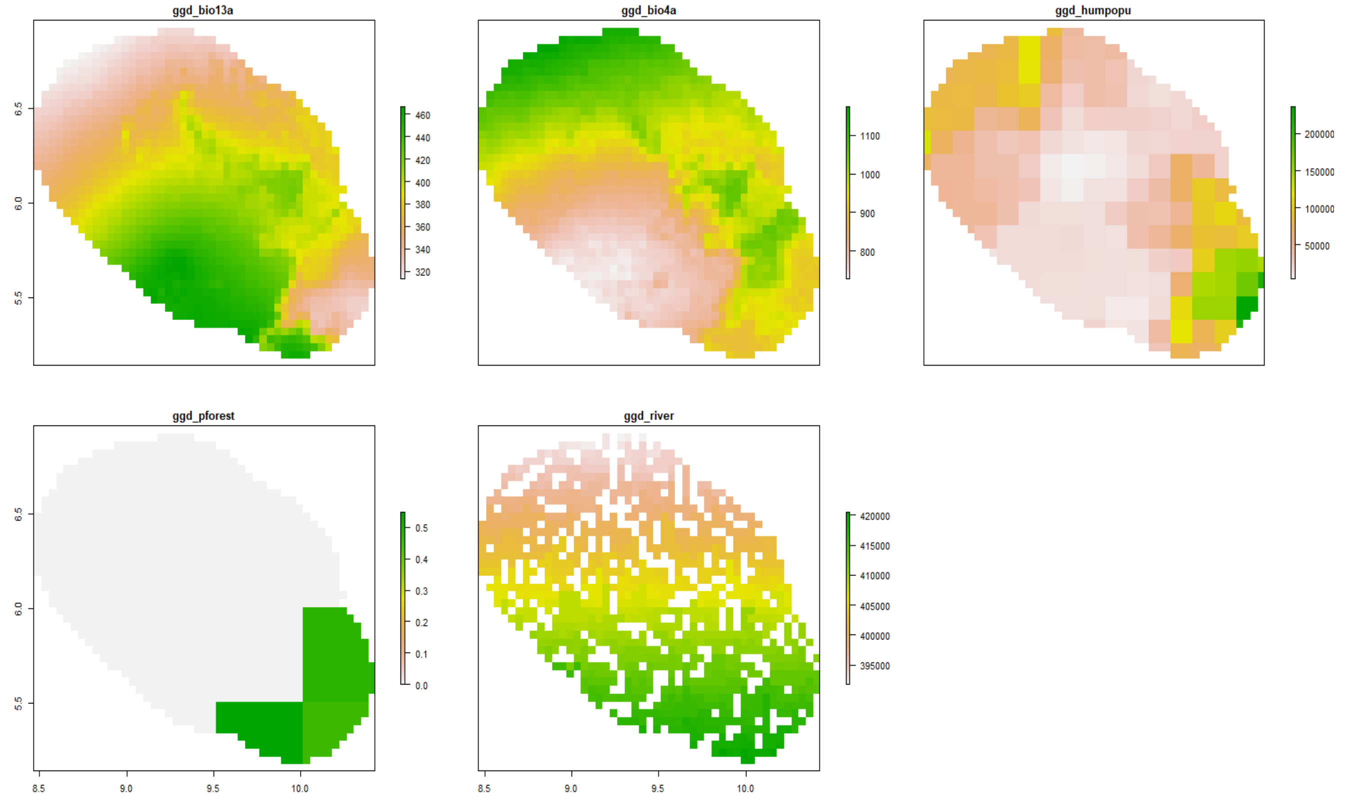

**Worst scenario**

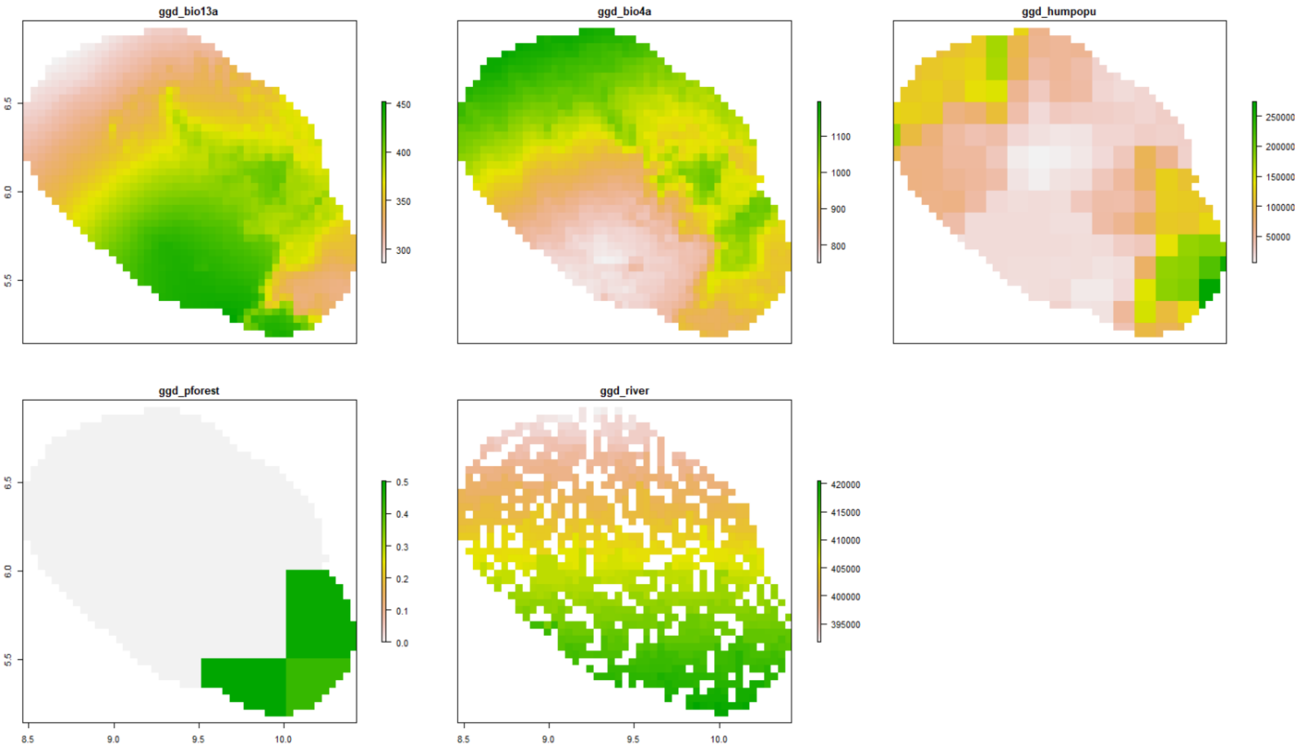


*Gorilla gorilla gorilla*

**Current**

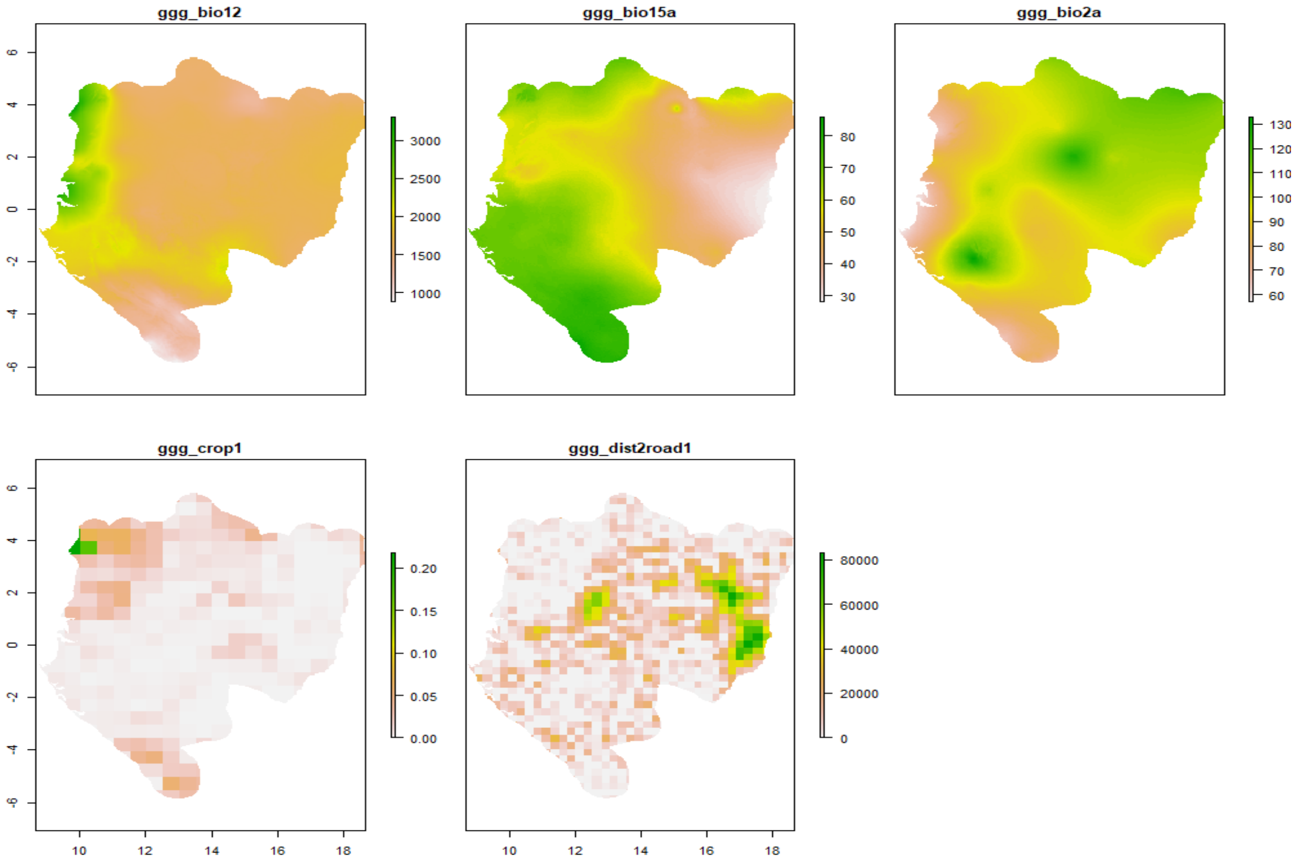

### Best scenario

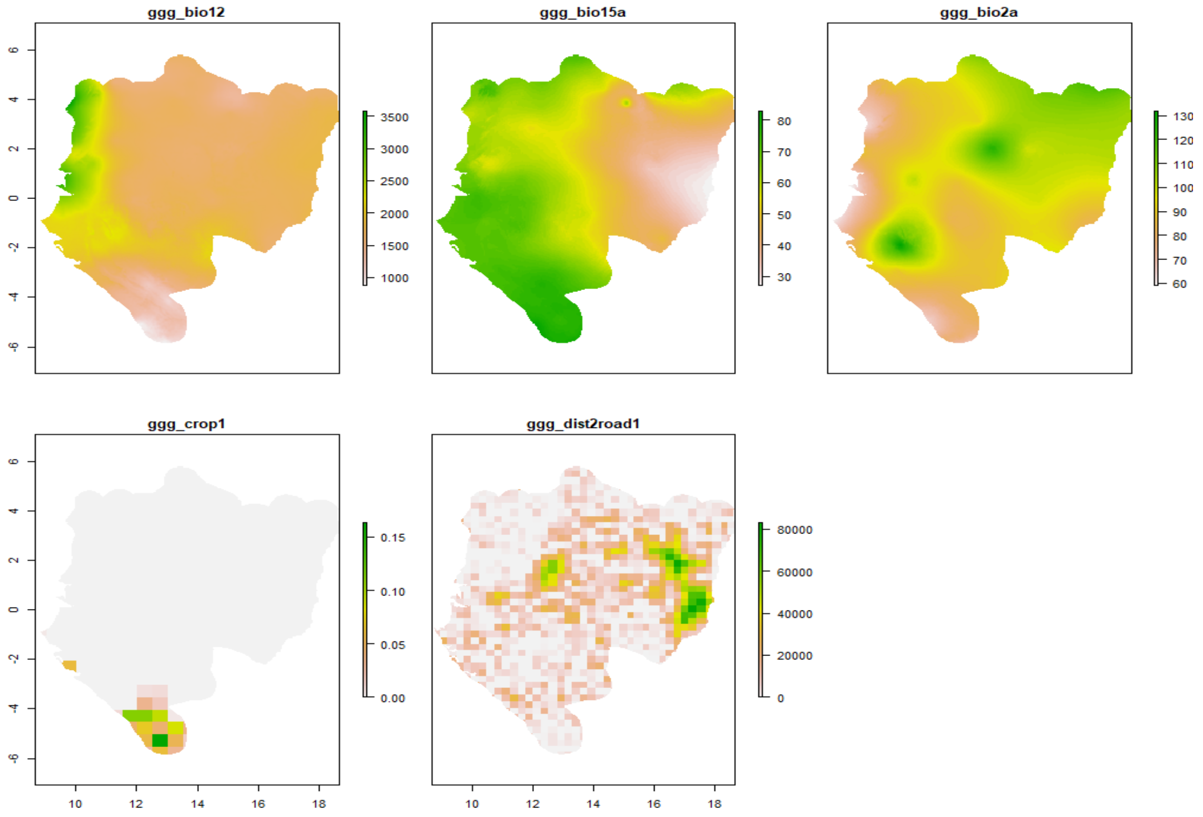


### Worst scenario

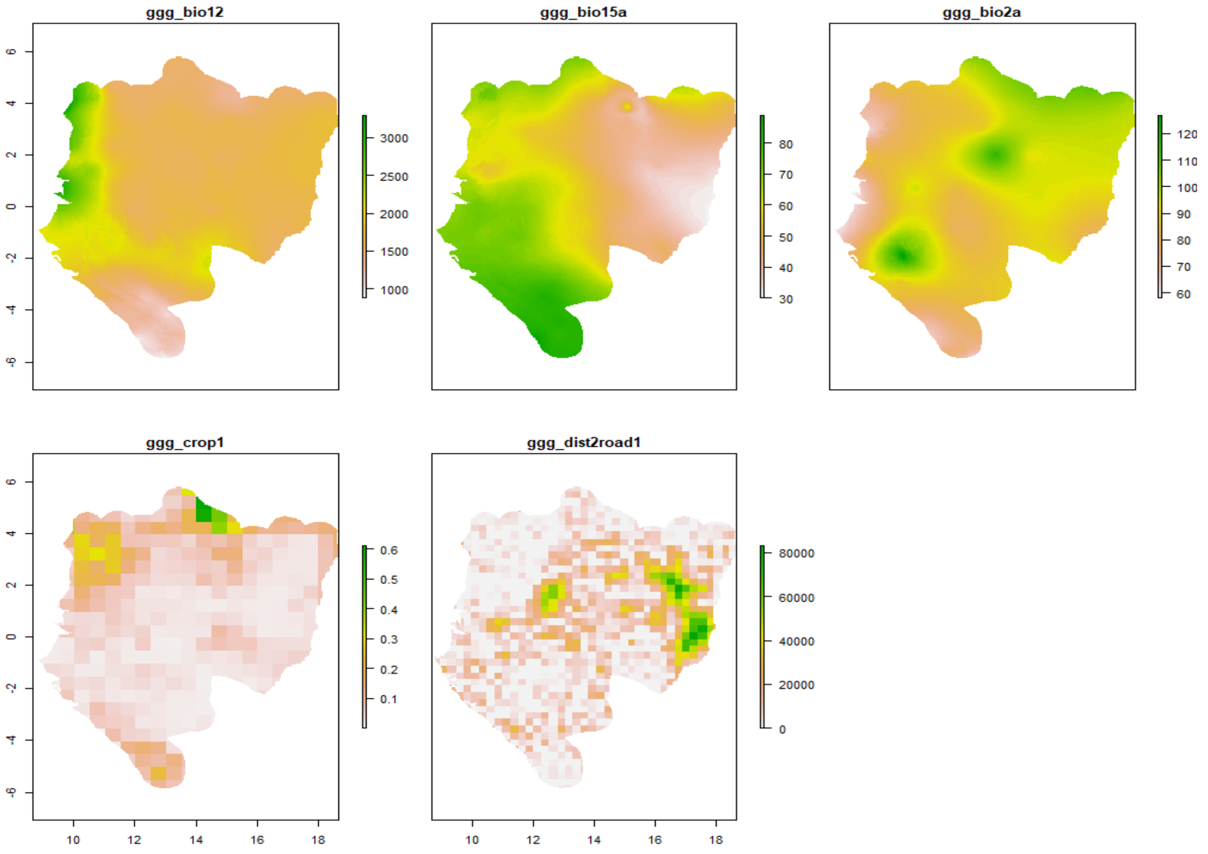

*Pan paniscus*

**Current**

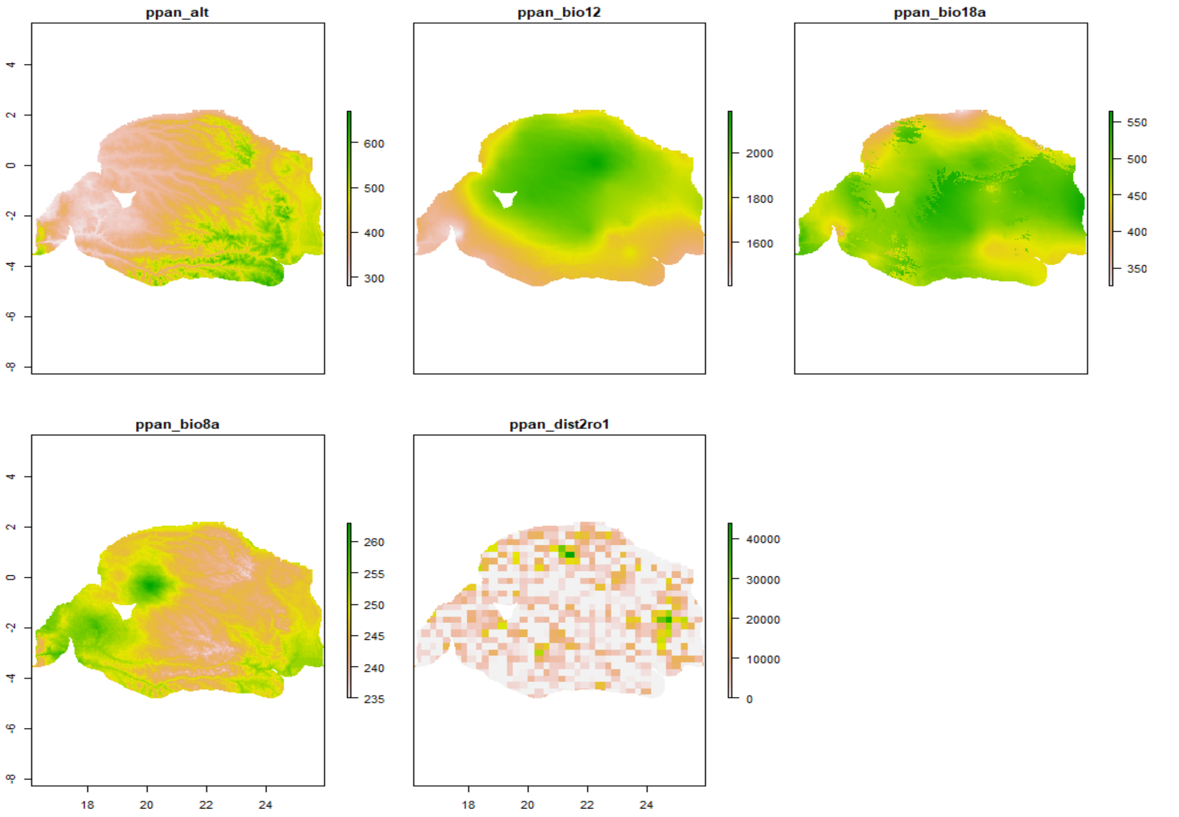


**Best scenario**

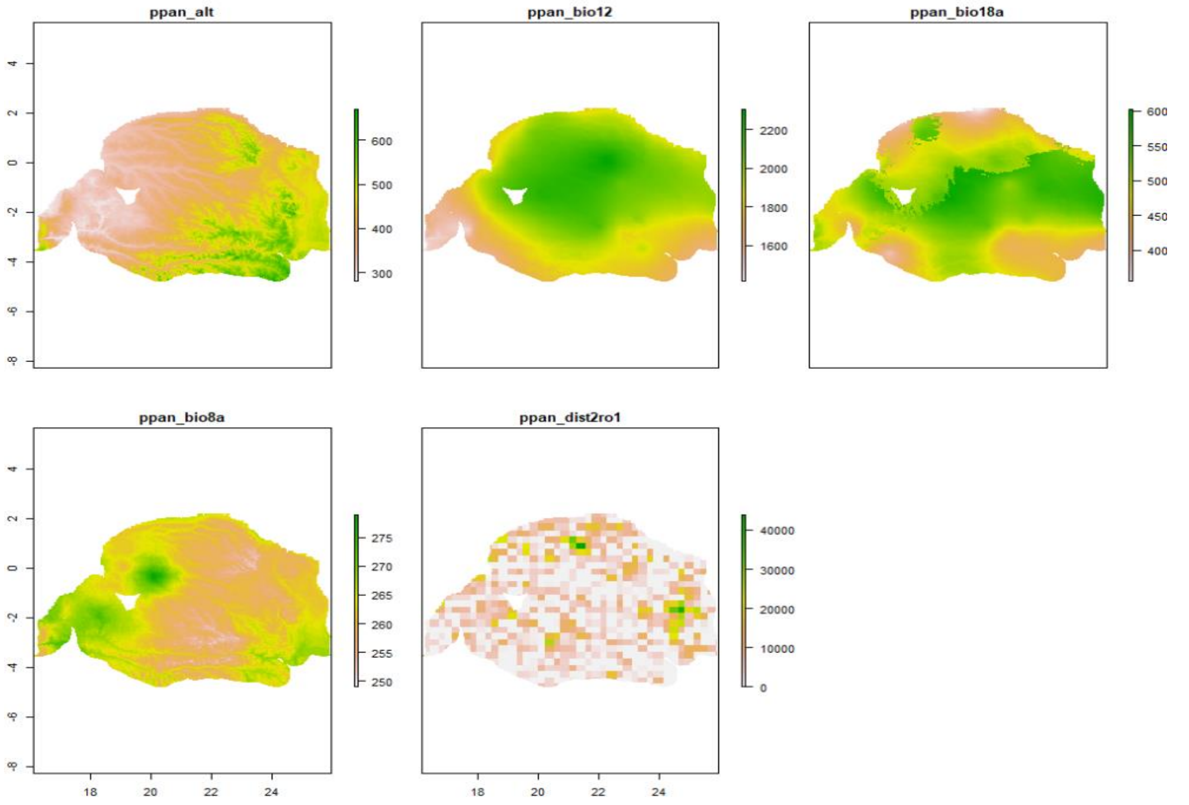

### Worst scenario

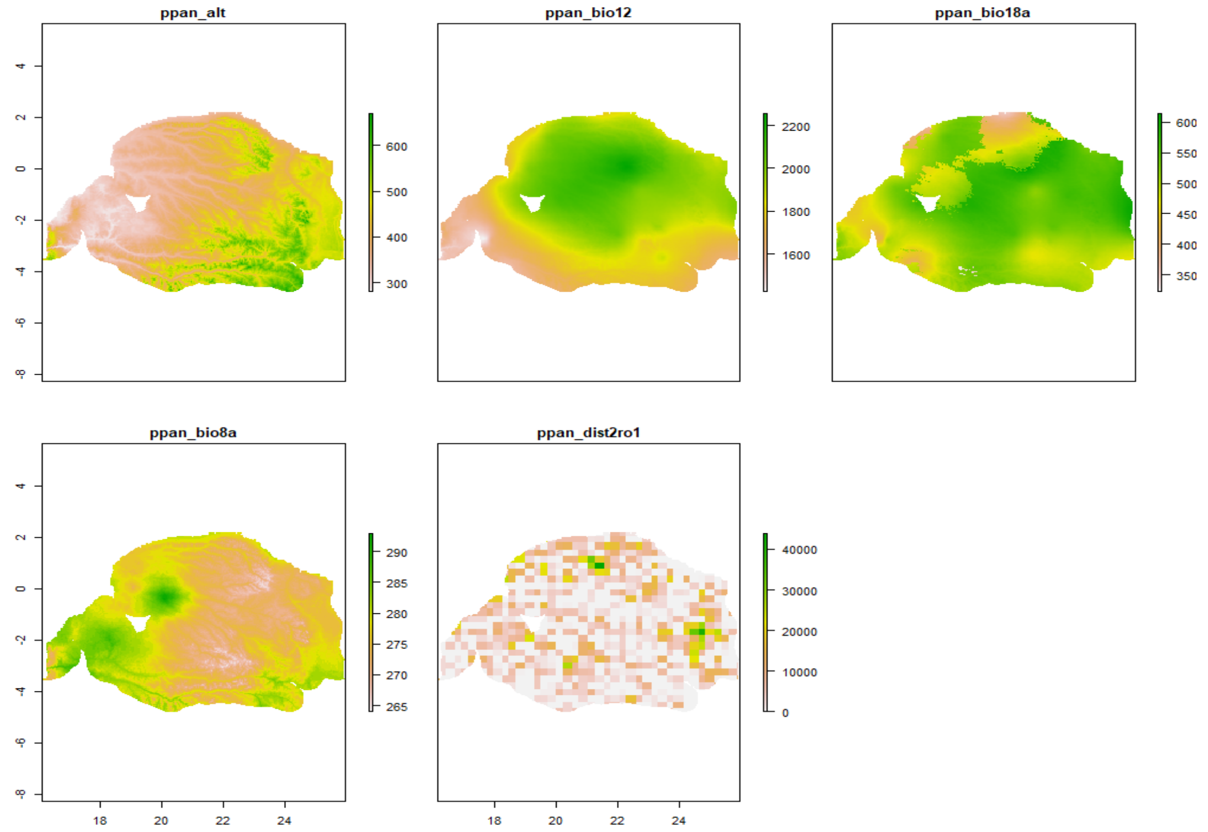


*Pan troglodytes ellioti*

### Current

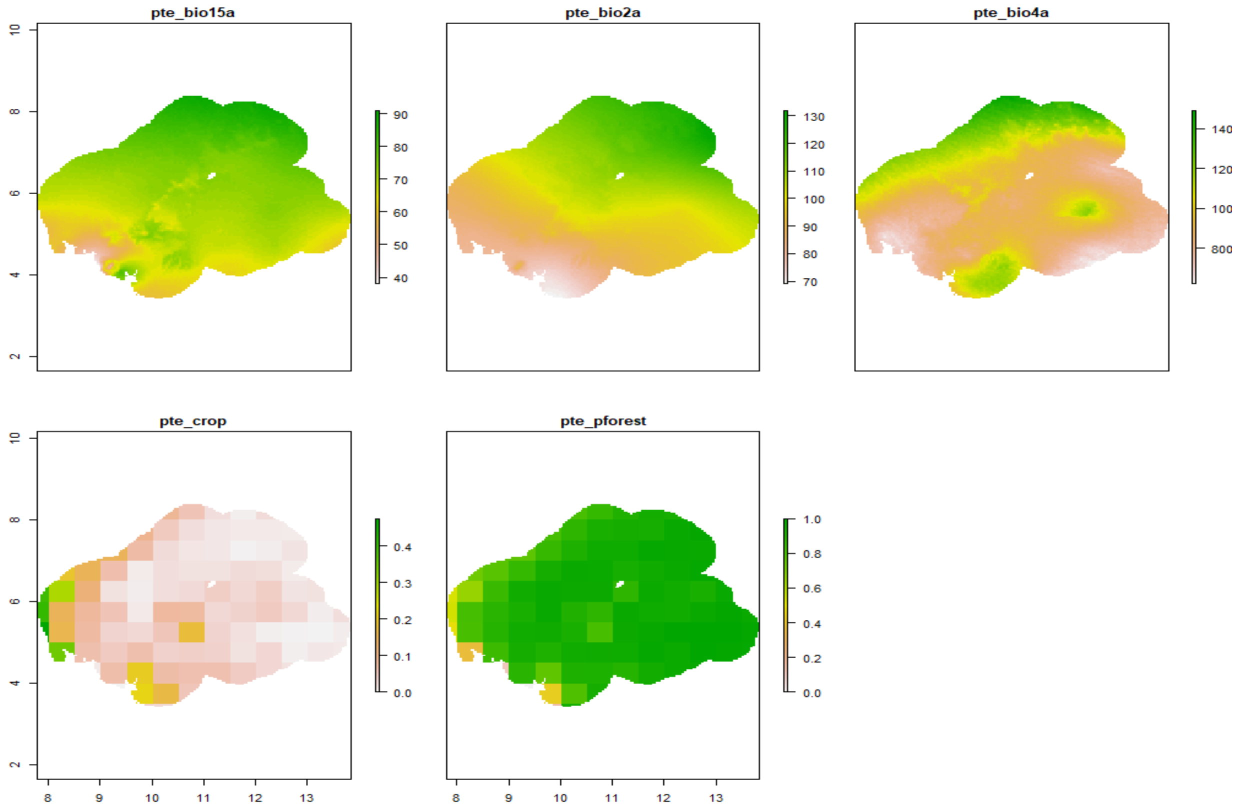

### Best scenario

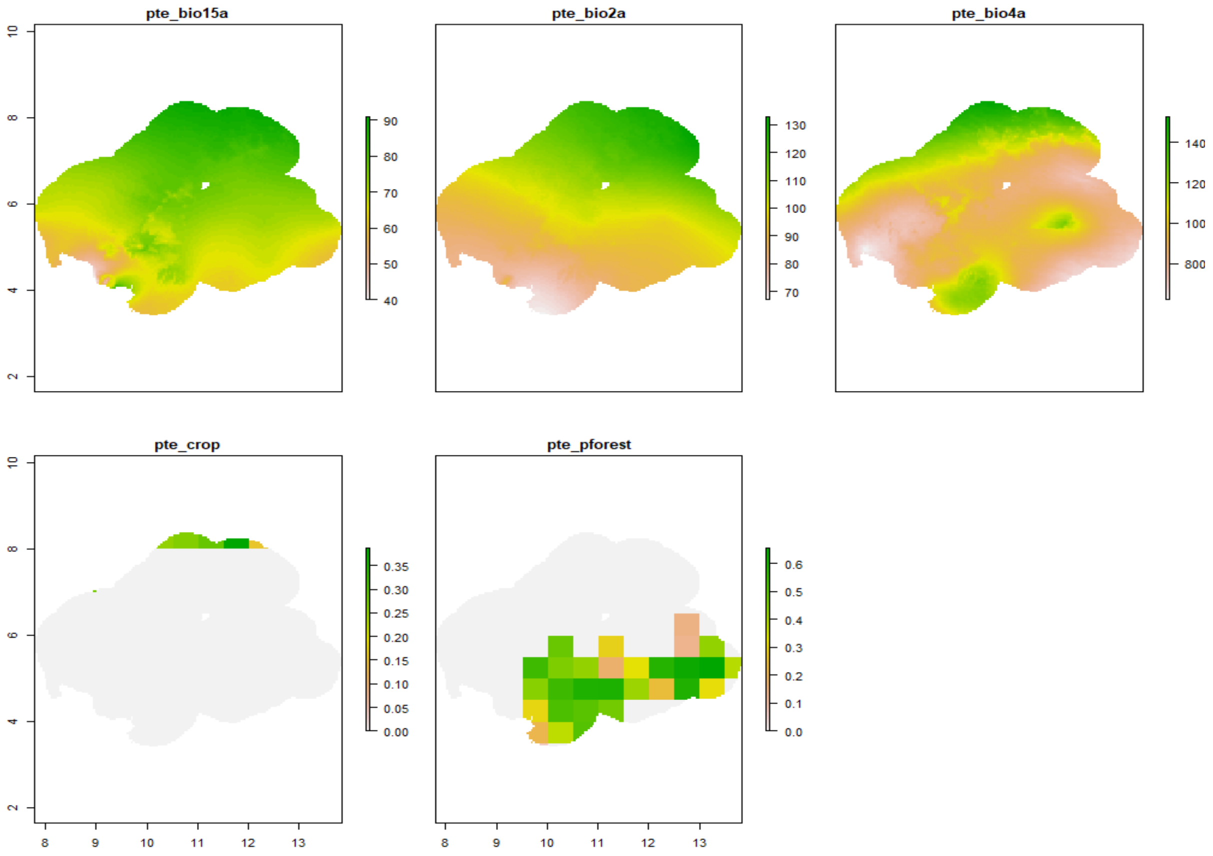


### Worst scenario

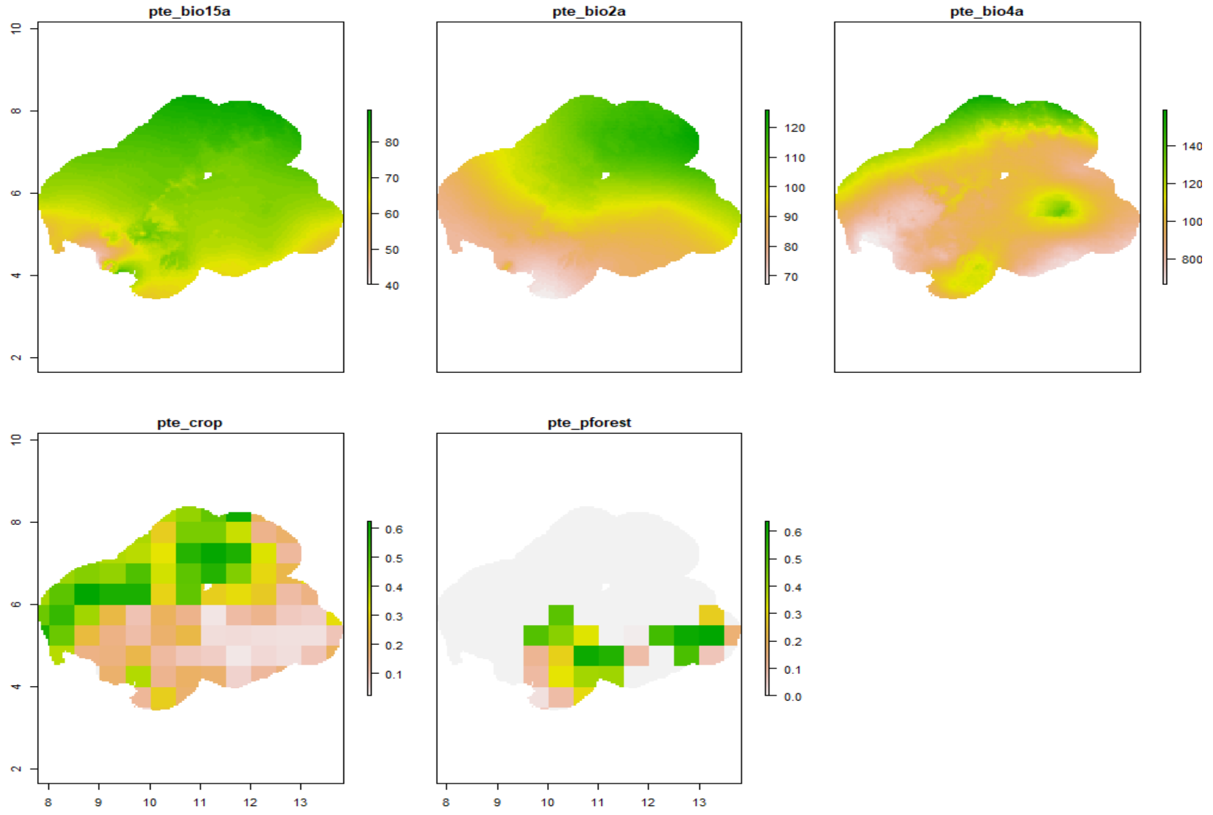

*Pan troglodytes schweinfurthii*

### Current

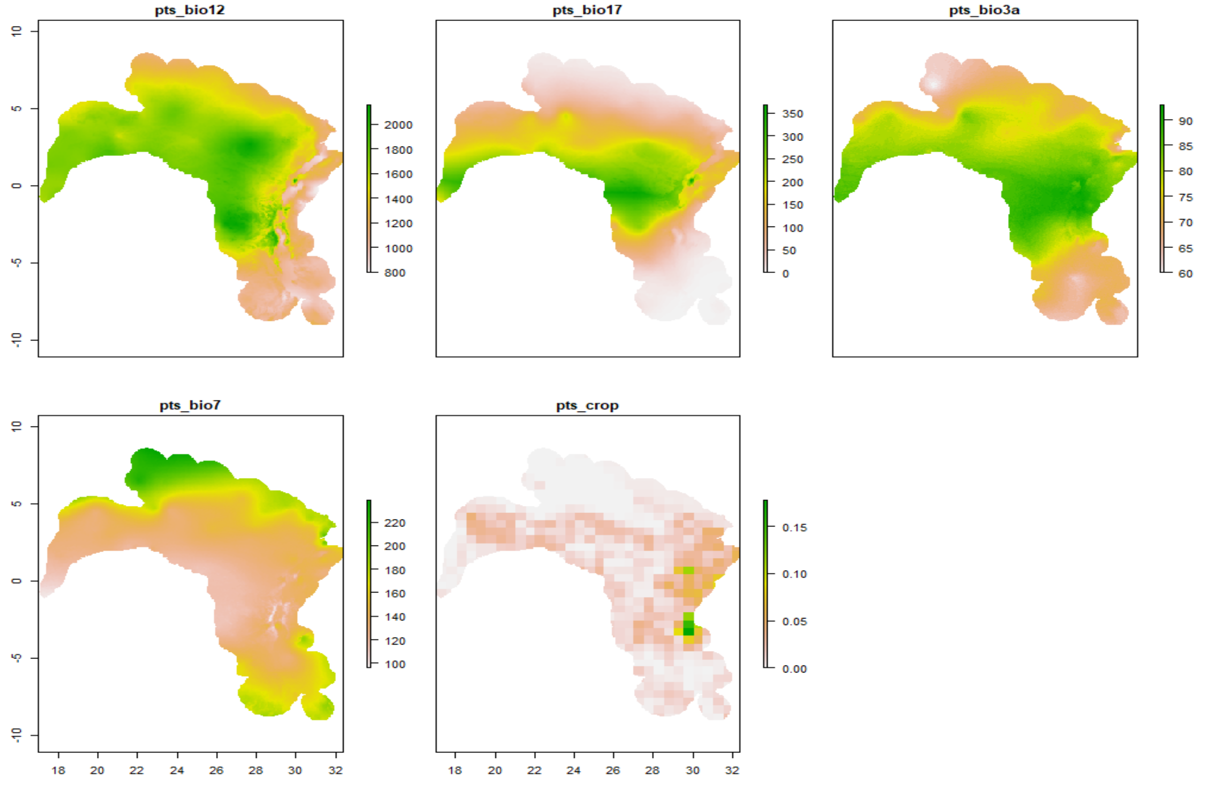


### Best scenario

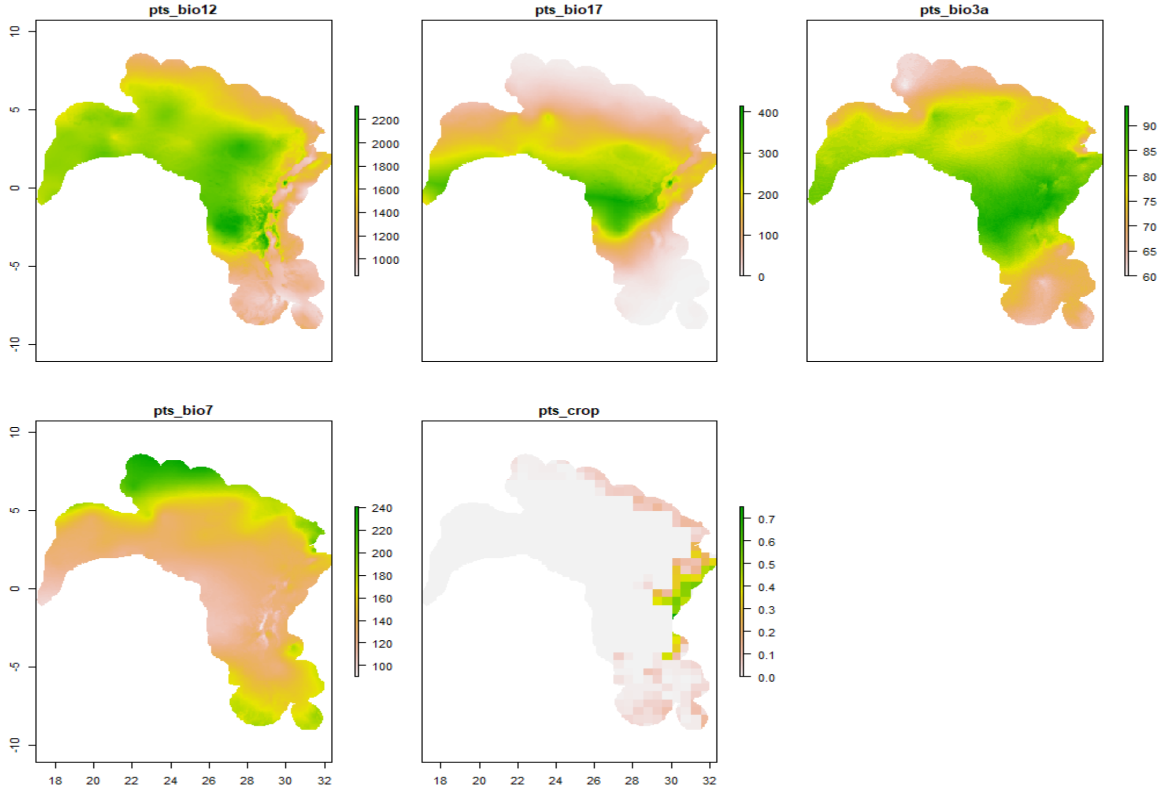

### Worst scenario

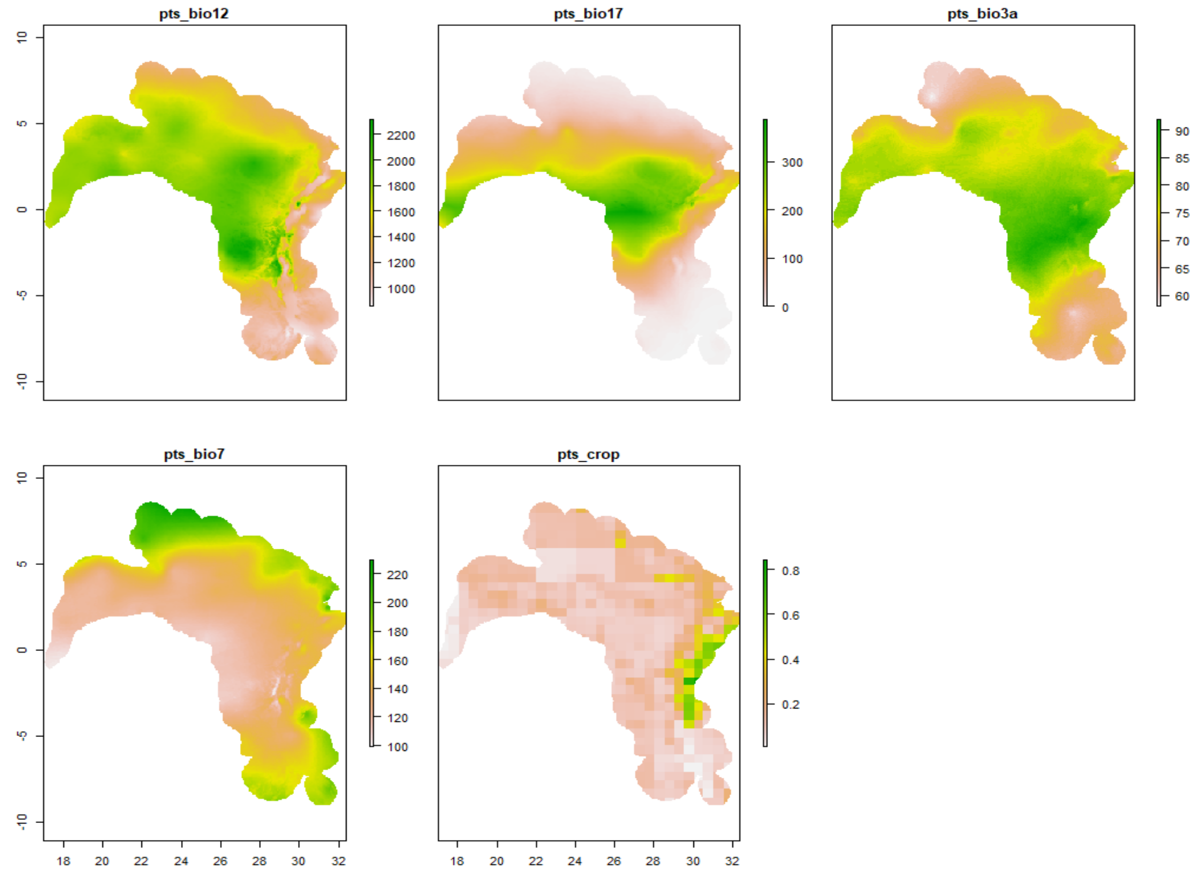


### *Pan troglodytes troglodytes*

### Current

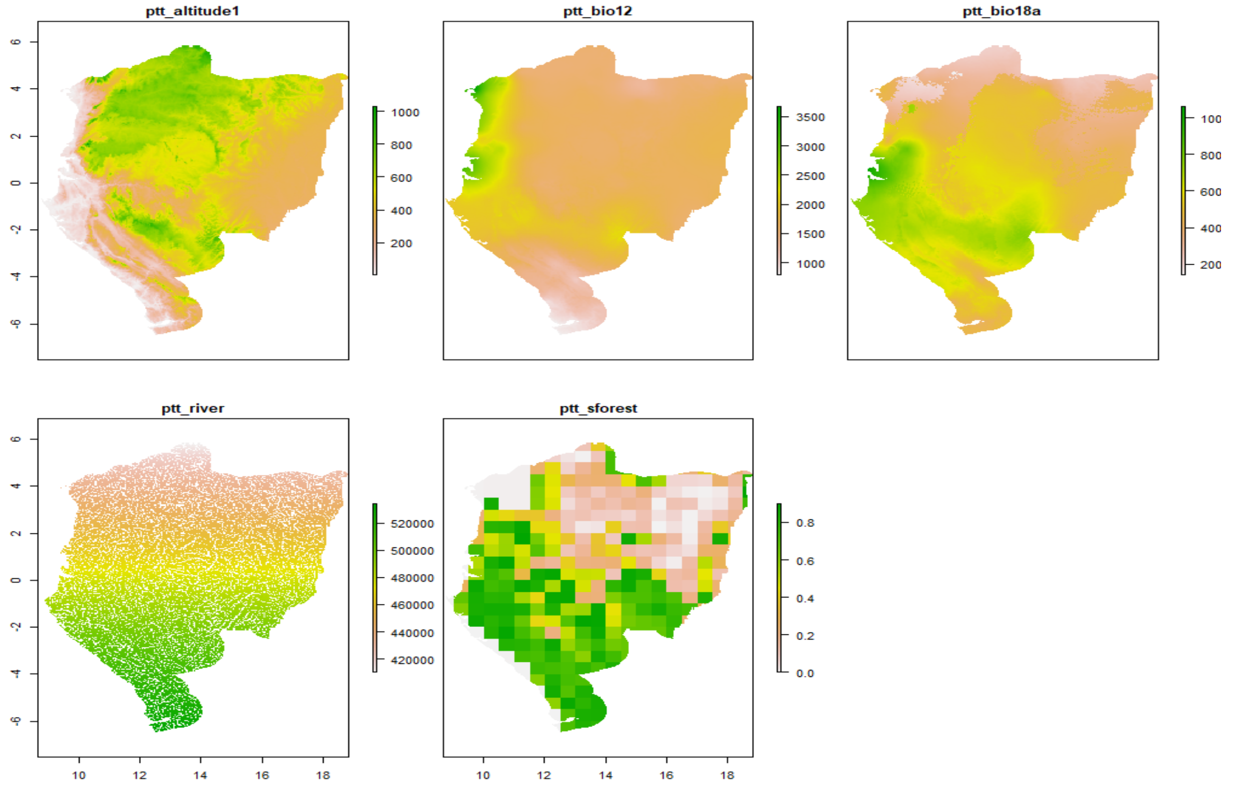

### Best scenario

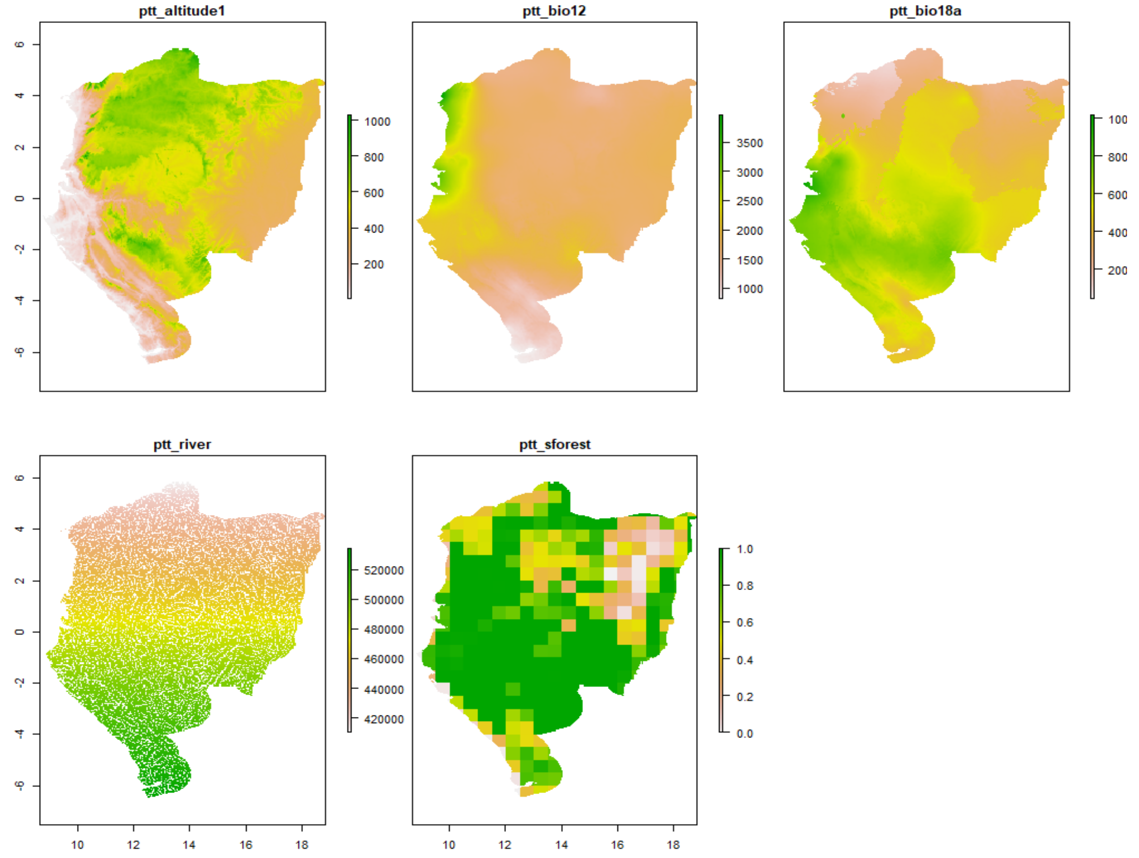


### Worst scenario

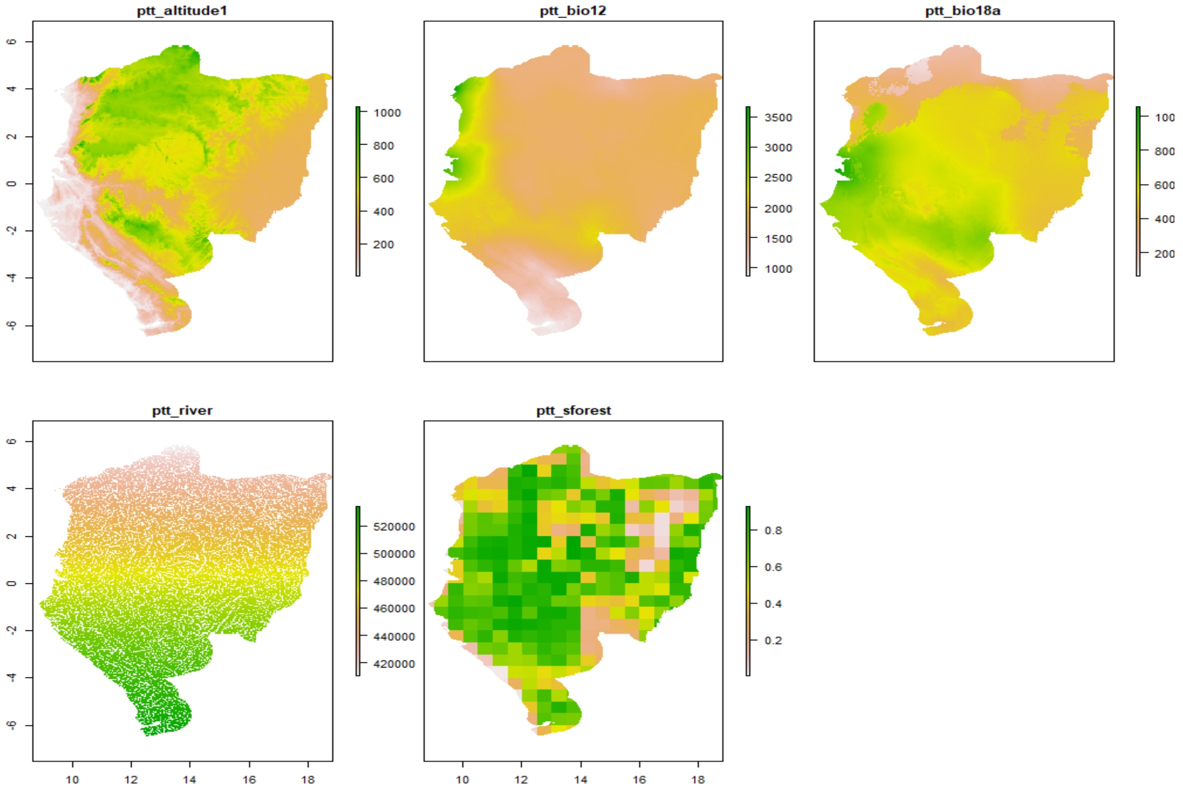

*Pan troglodytes verus*

**Current**

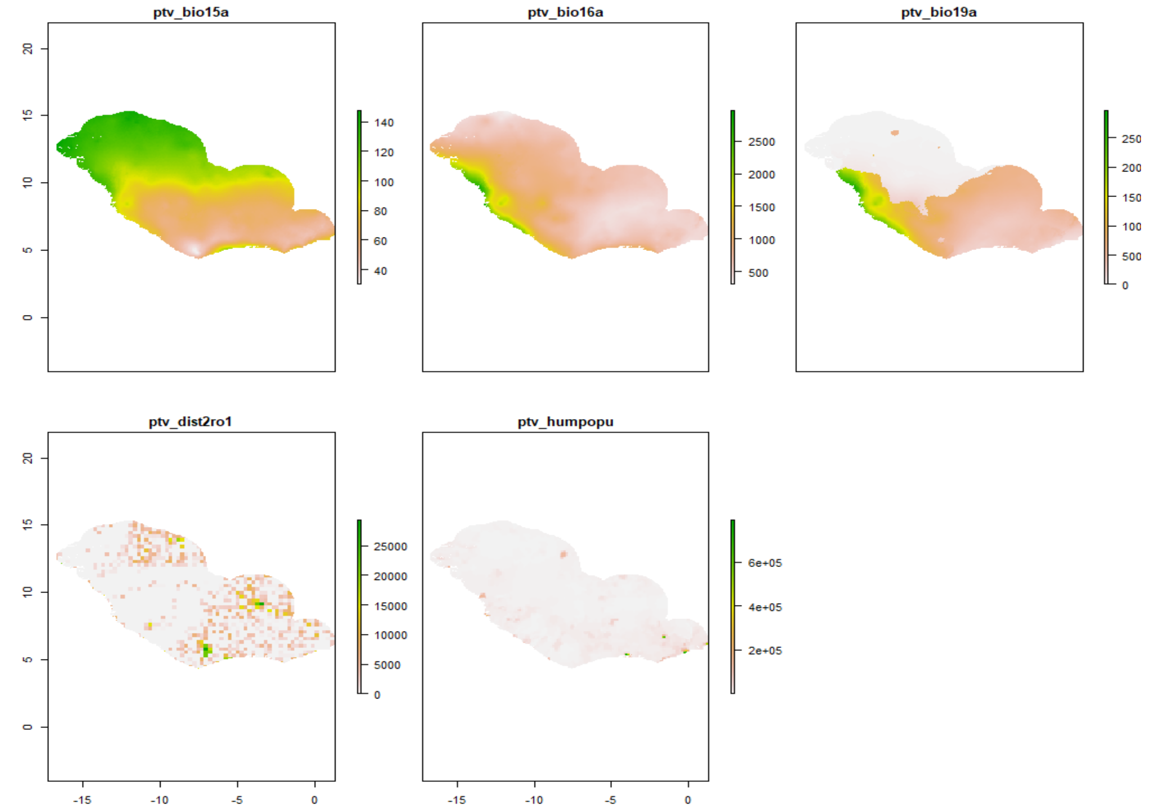


**Best scenario**

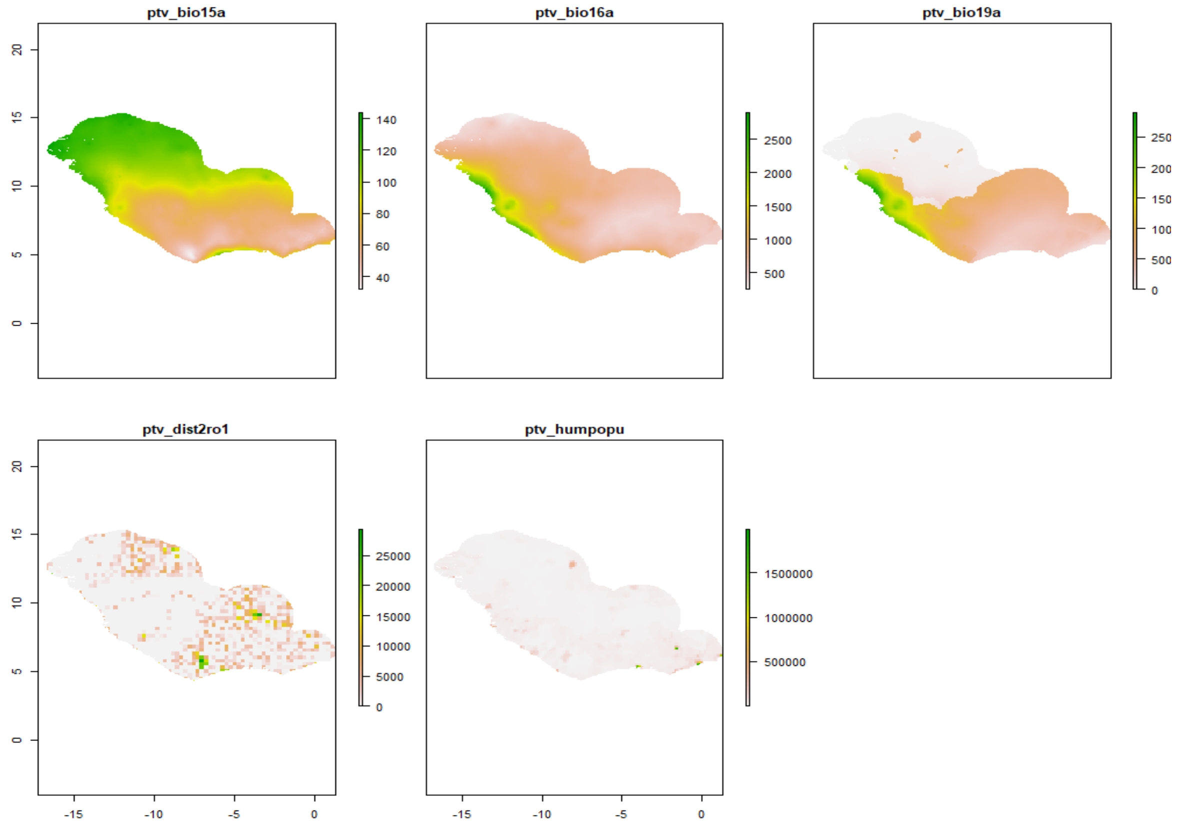

### Worst scenario

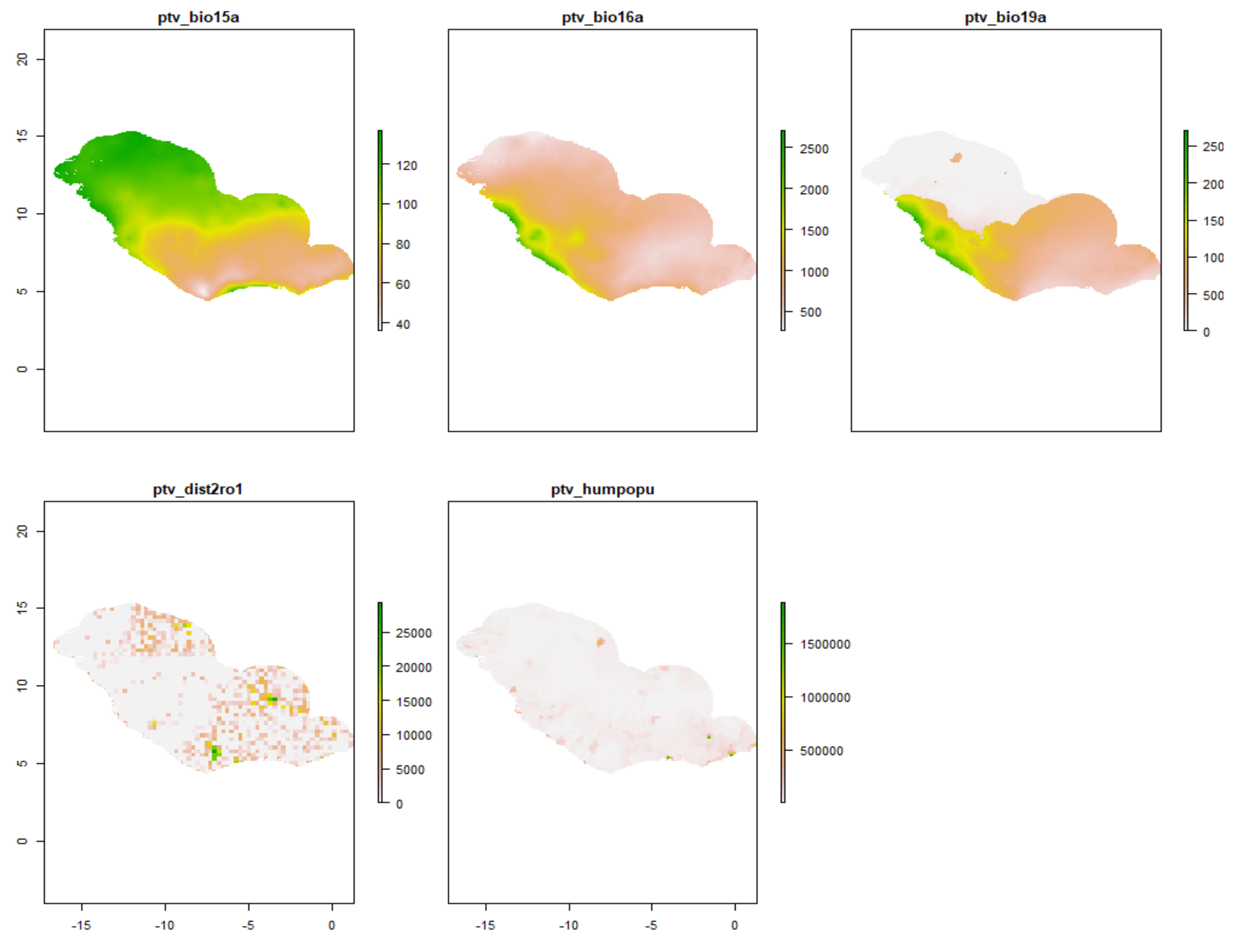


Figure S1.5. Variable predictors under current and future conditions used for predicting African ape distribution. Note that all temperature variables are plotted in their original scale (i.e. °C * 10 or °C *100 for the seasonal variation of temperature), as well as all land-use variables (i.e. % *100).

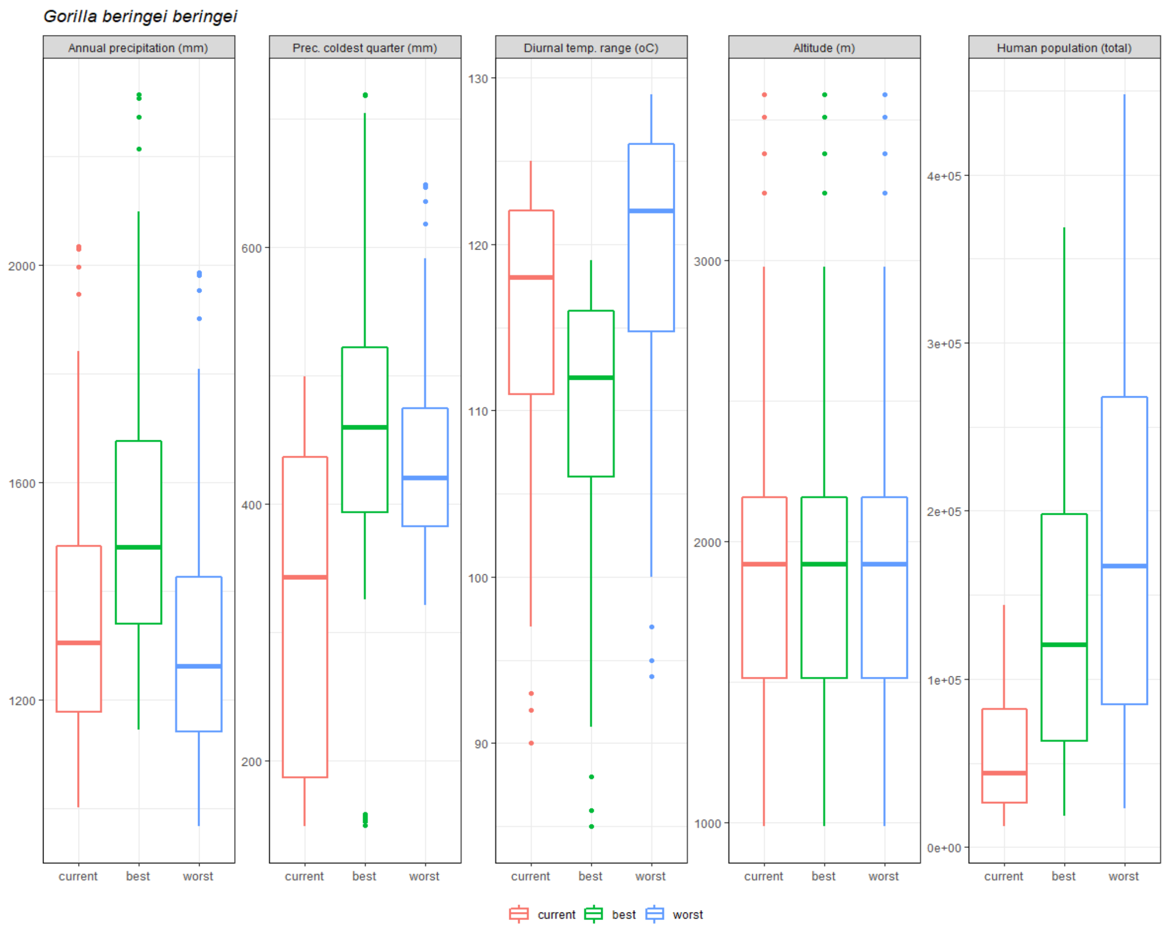
Gorilla beringei beringei
Annual precipitation (mm)
Prec. coldest quarter (mm)
Diurnal temp. range (oC)
Altitude (m)
Human population (total)
current
best
worst

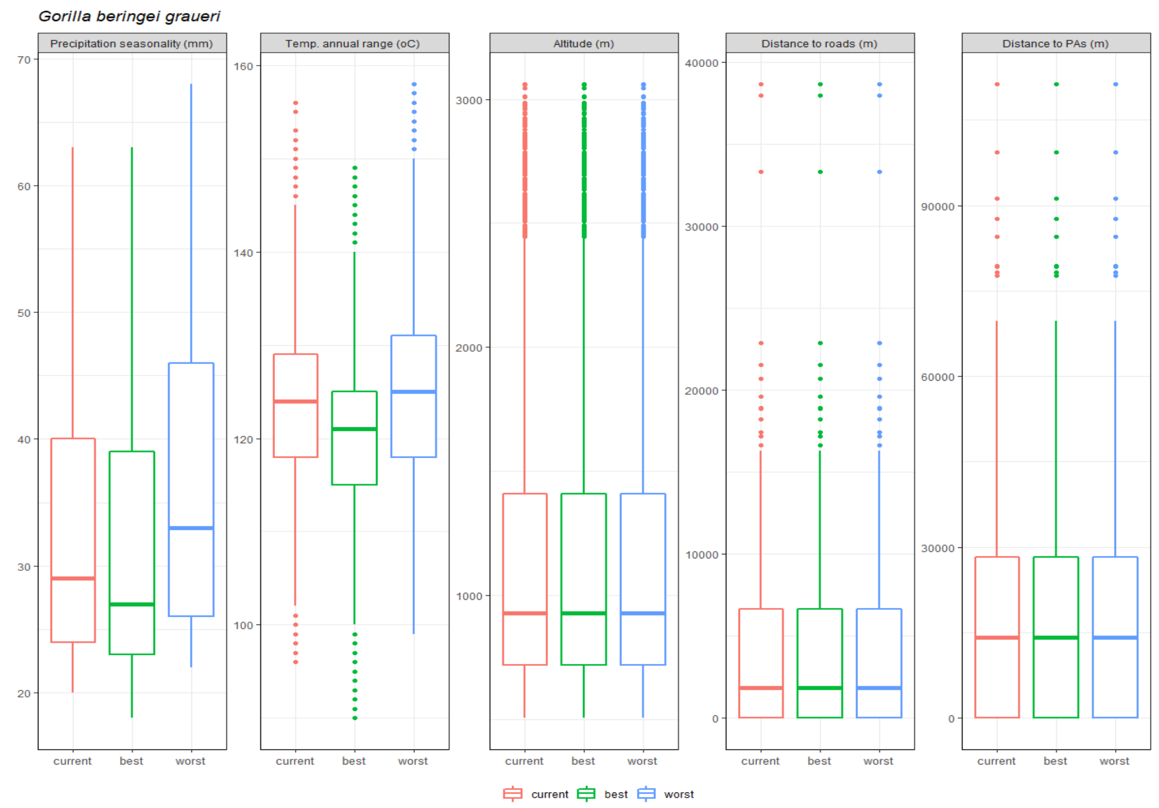
Gorilla beringei graueri
Precipitation seasonality (mm)
Temp. annual range (oC)
Altitude (m)
Distance to roads (m)
Distance to PAs (m)
current
best
worst

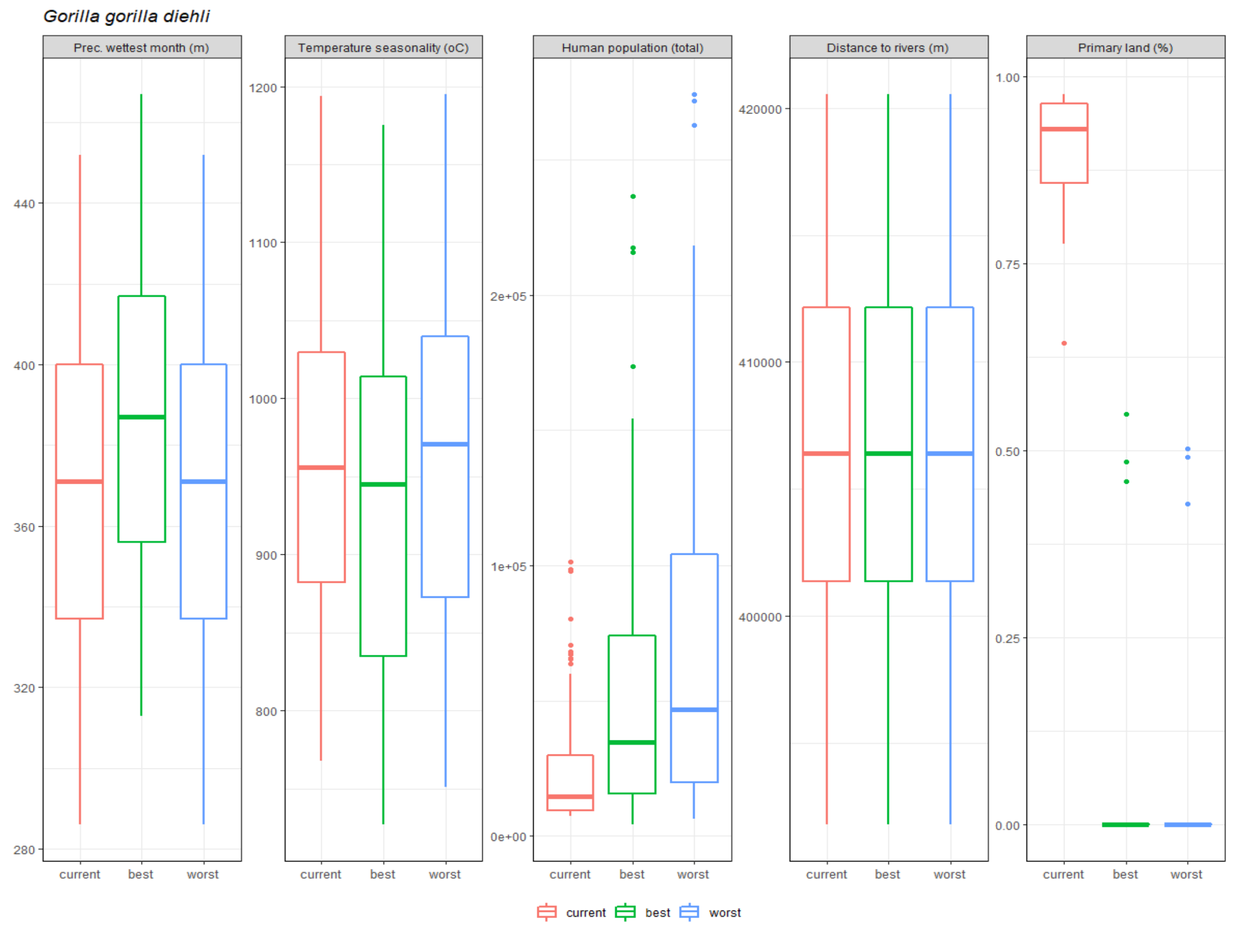
Gorilla gorilla diehli
Prec. wettest month (m)
Temperature seasonality (oC)
Human population (total)
Distance to rivers (m)
Primary land (%)
280
320
360
400
440
800
900
1000
1100
1200
0e+00
1e+05
2e+05
400000
410000
420000
0.00
0.25
0.50
0.75
1.00
current
best
worst

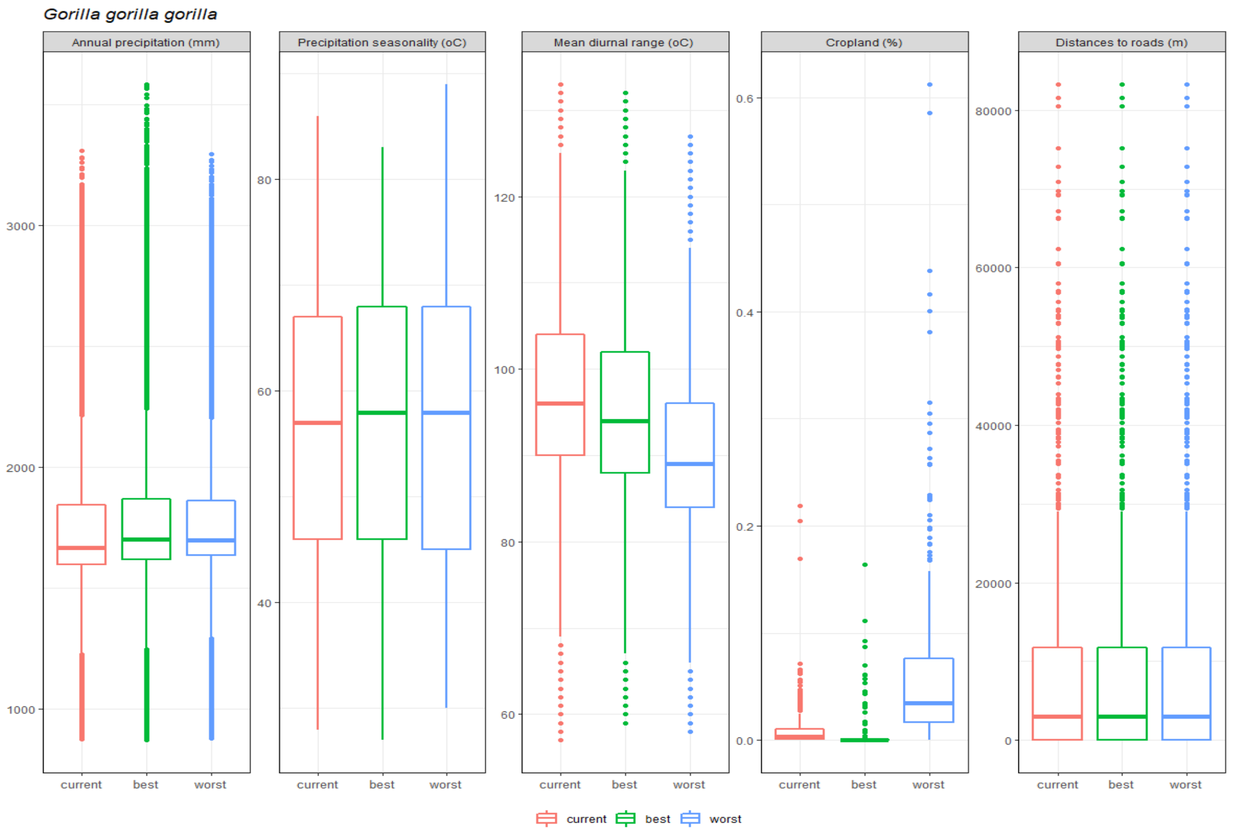
Gorilla gorilla gorilla
Annual precipitation (mm)
Precipitation seasonality (oC)
Mean diurnal range (oC)
Cropland (%)
Distances to roads (m)
1000
2000
3000
40
60
80
60
80
100
120
0.0
0.2
0.4
0.6
0
20000
40000
60000
80000
current
best
worst

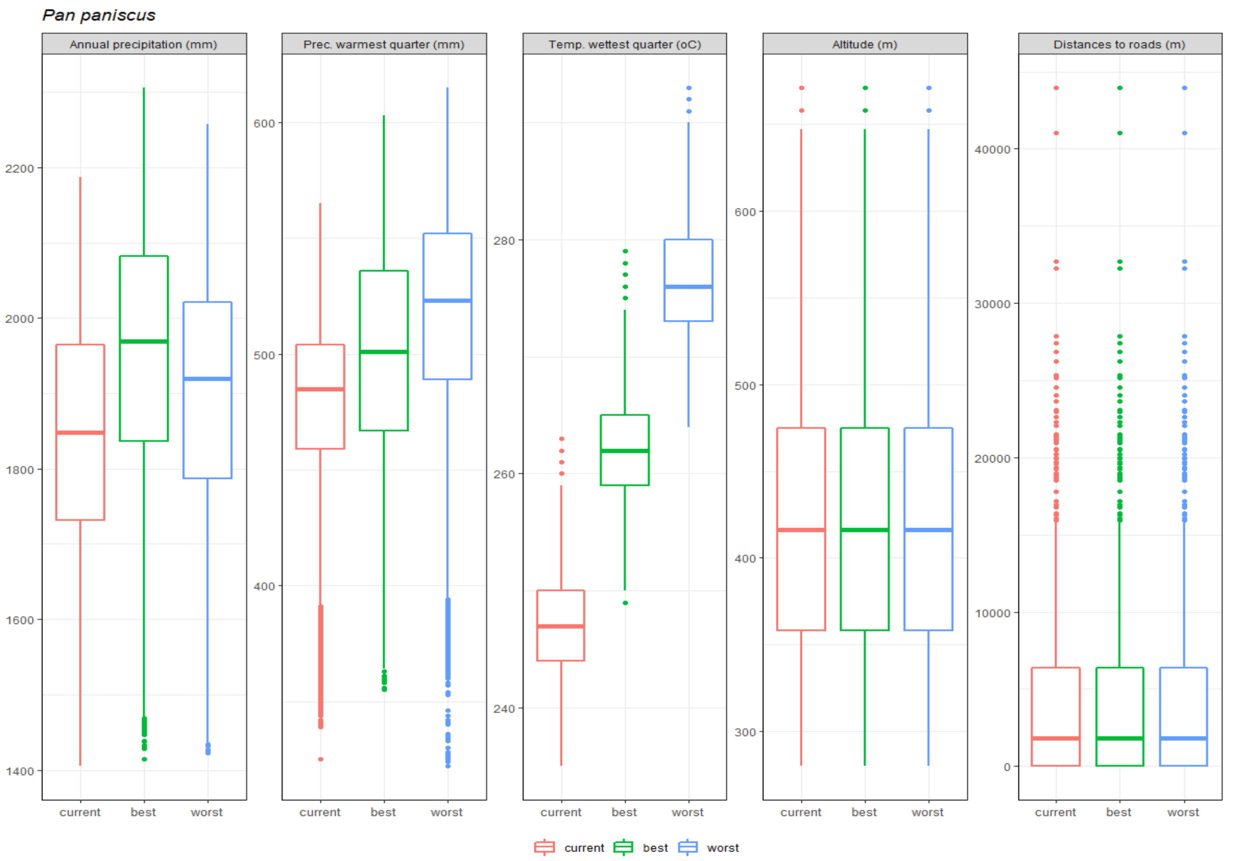
Pan paniscus
Annual precipitation (mm)
Prec. warmest quarter (mm)
Temp. wettest quarter (oC)
Altitude (m)
Distances to roads (m)
2200
2000
1800
1600
1400
600
500
400
280
260
240
600
500
400
300
40000
30000
20000
10000
0
current
best
worst
current
best
worst

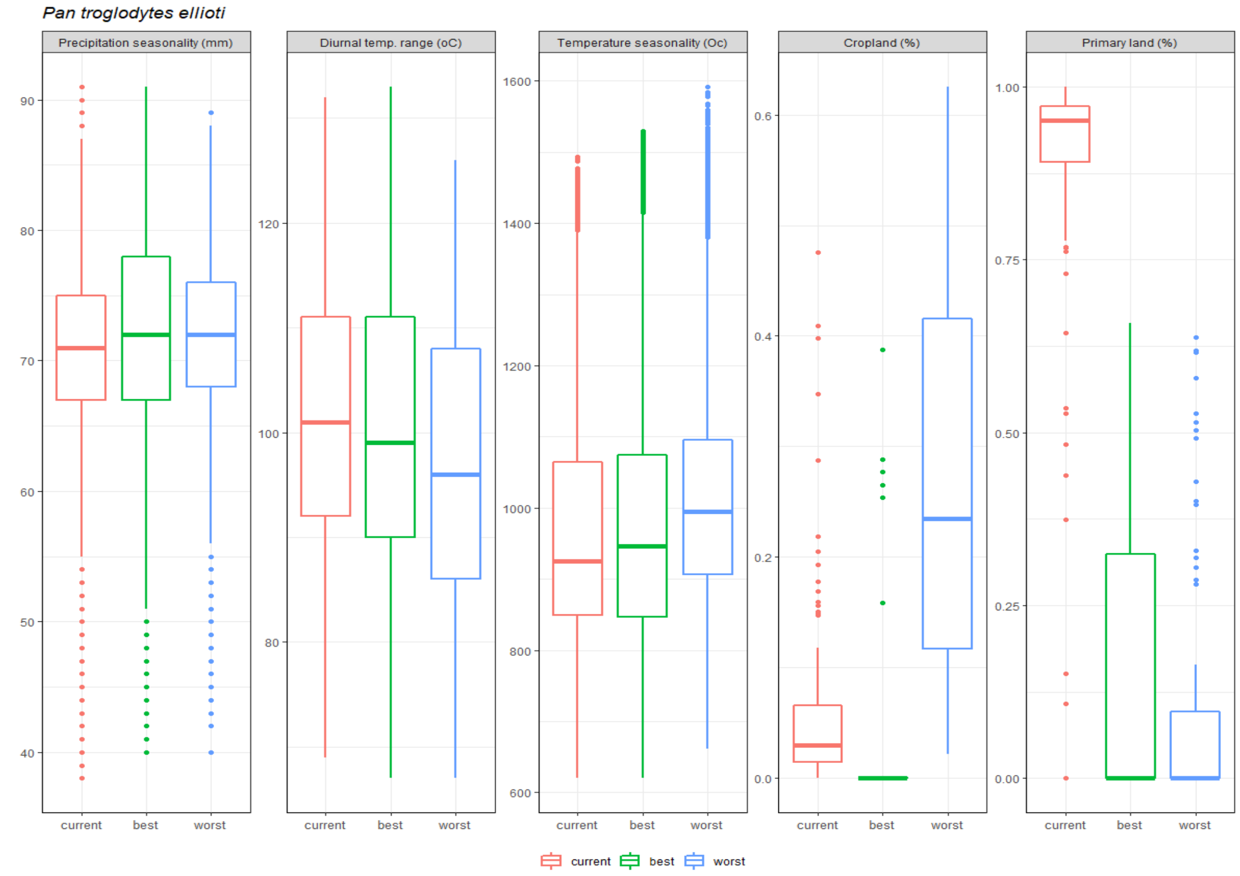
Pan troglodytes ellioti
Precipitation seasonality (mm)
Diurnal temp. range (oC)
Temperature seasonality (Oc)
Cropland (%)
Primary land (%)
90
80
70
60
50
40
120
100
80
1600
1400
1200
1000
800
600
0.6
0.4
0.2
0.0
1.00
0.75
0.50
0.25
0.00
current
best
worst
current
best
worst

*Pan troglodytes schweinfurthii*

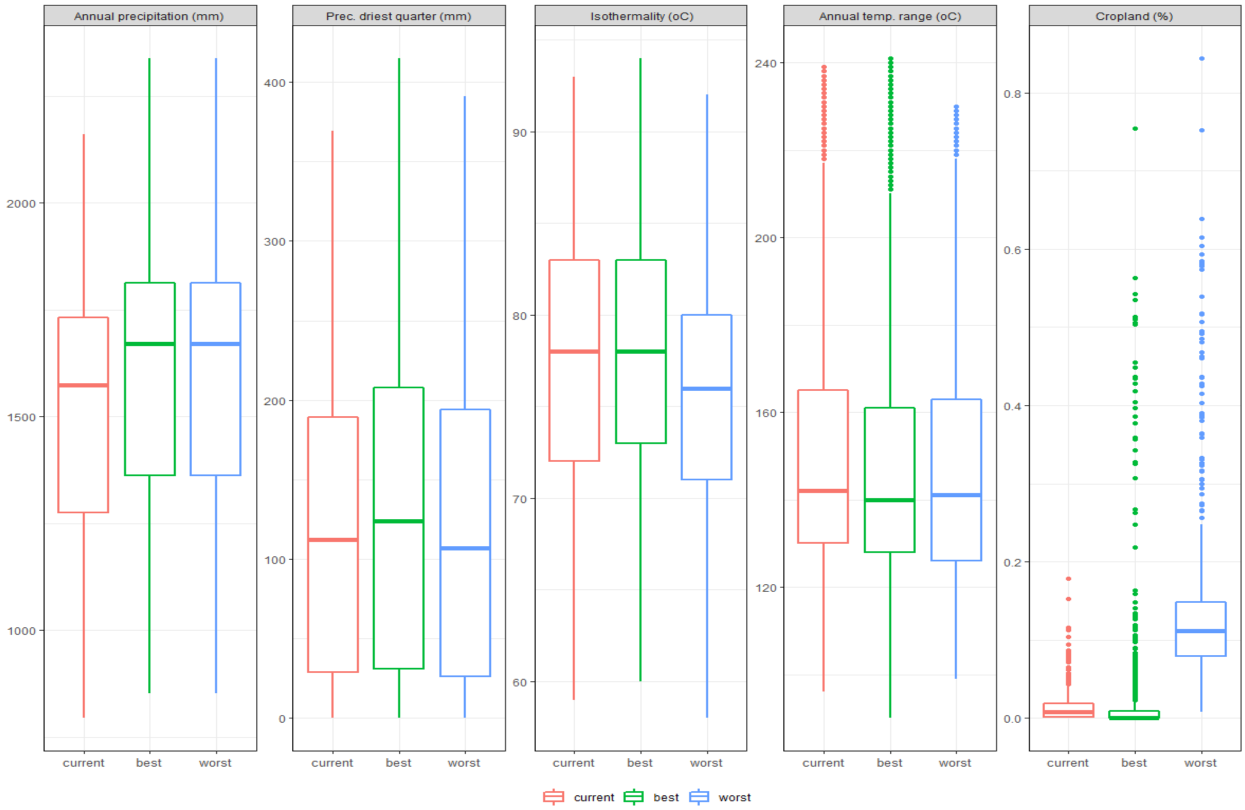


*Pan troglodytes troglodytes*

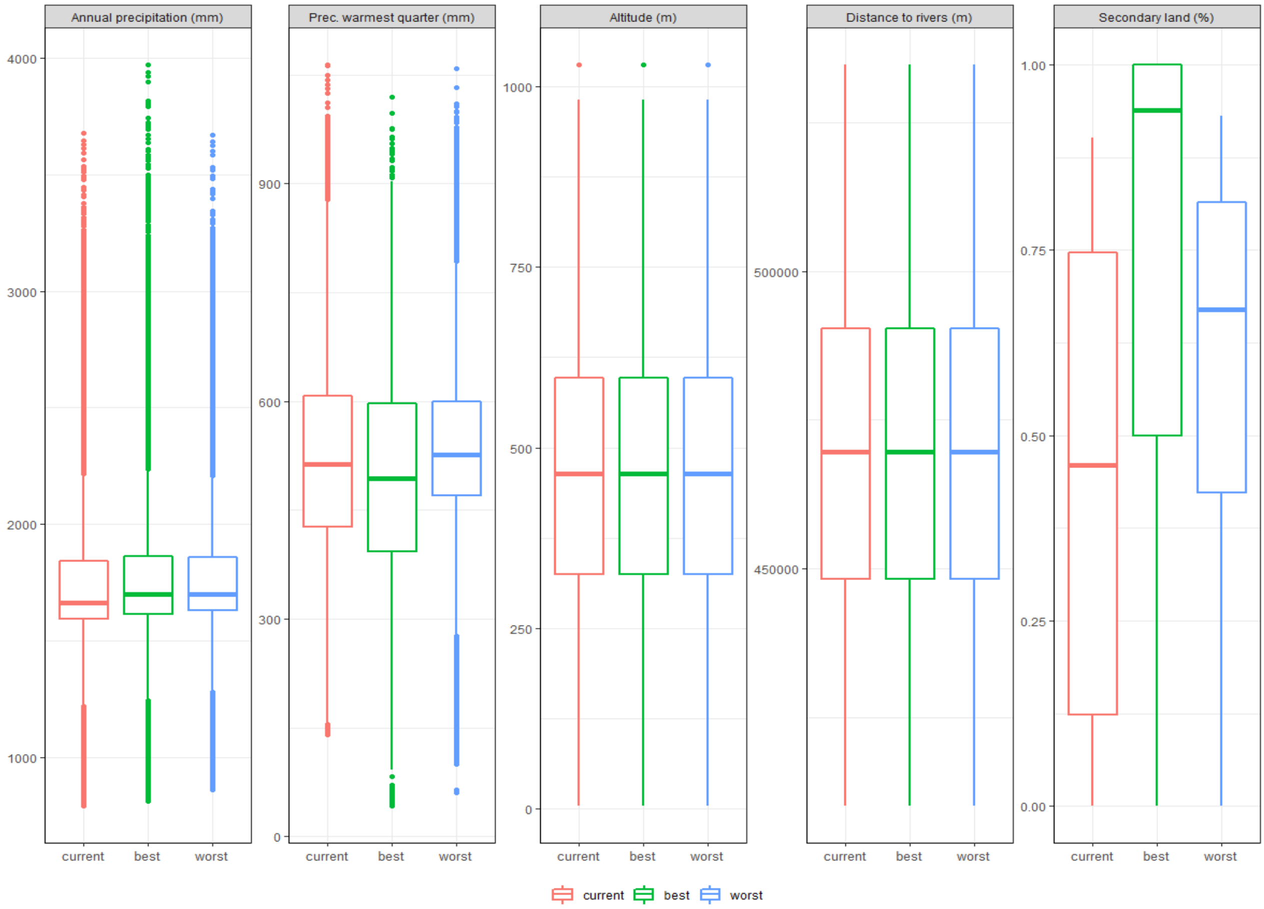

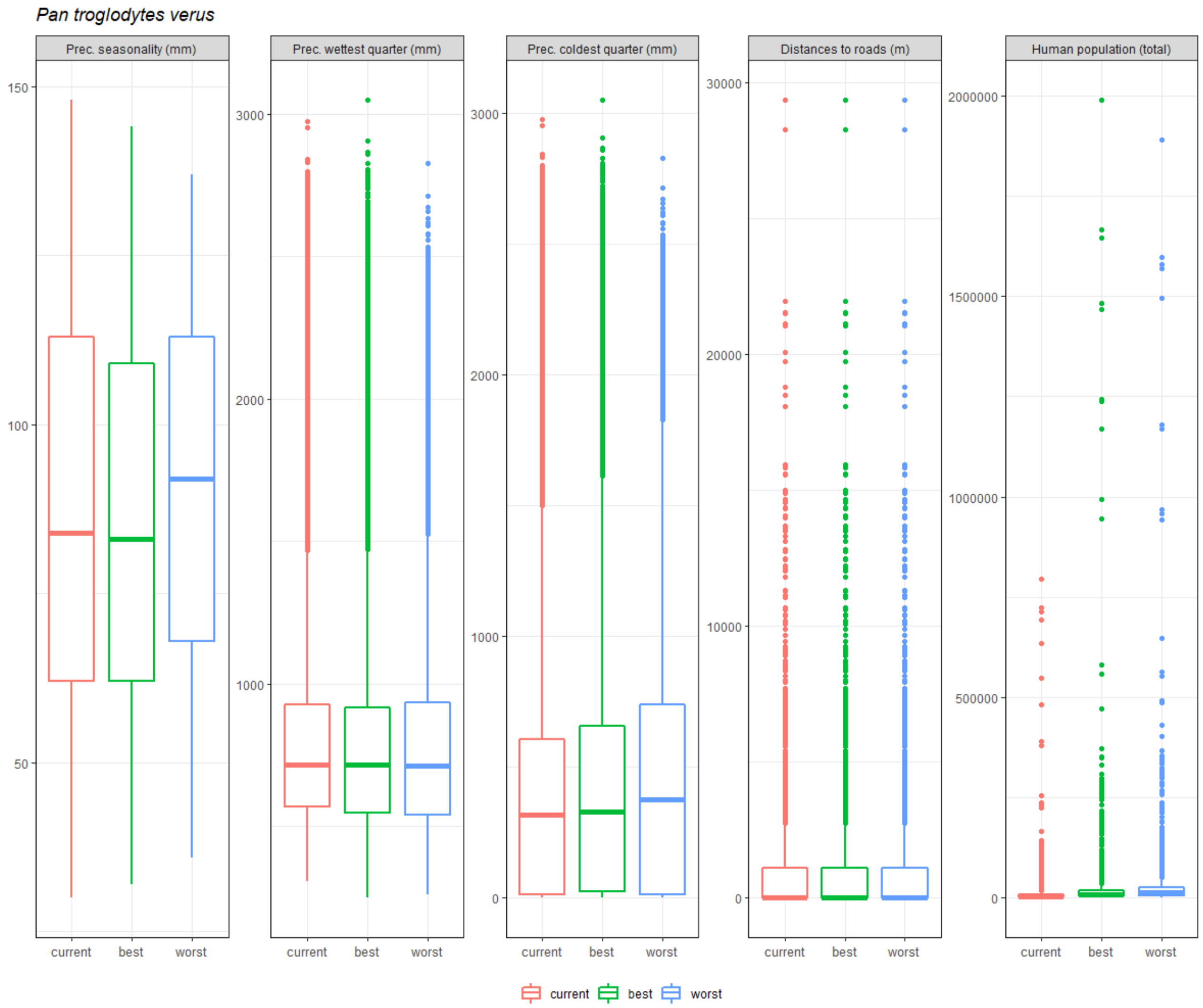


Figure S1.6. Range of values of the variable predictors under current and future conditions used for predicting African ape distribution. Note that all temperature variables are plotted in their original scale (i.e. °C * 10 or °C *100 for the seasonal variation of temperature), https://www.worldclim.org/), as well as all land-use variables (i.e. % *100).

*Gorilla beringei beringei*

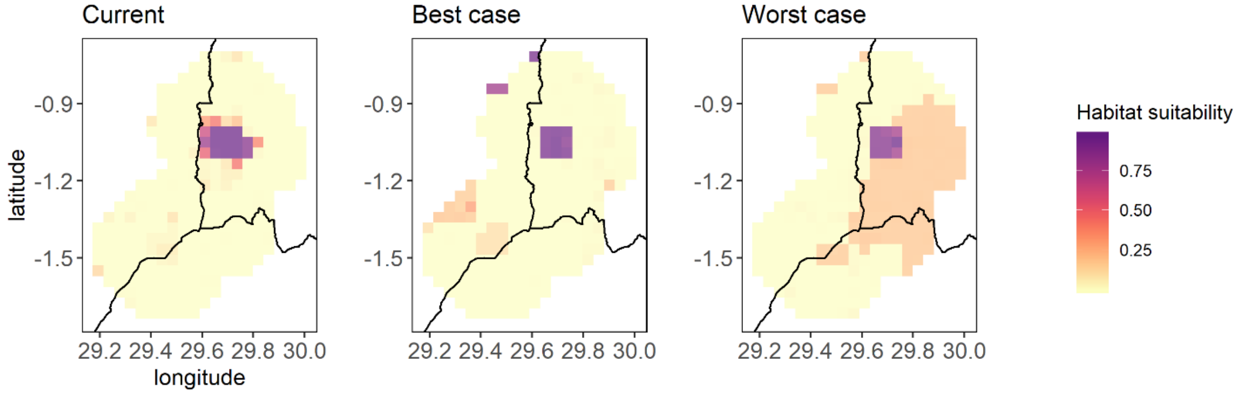


*Gorilla beringei graueri*

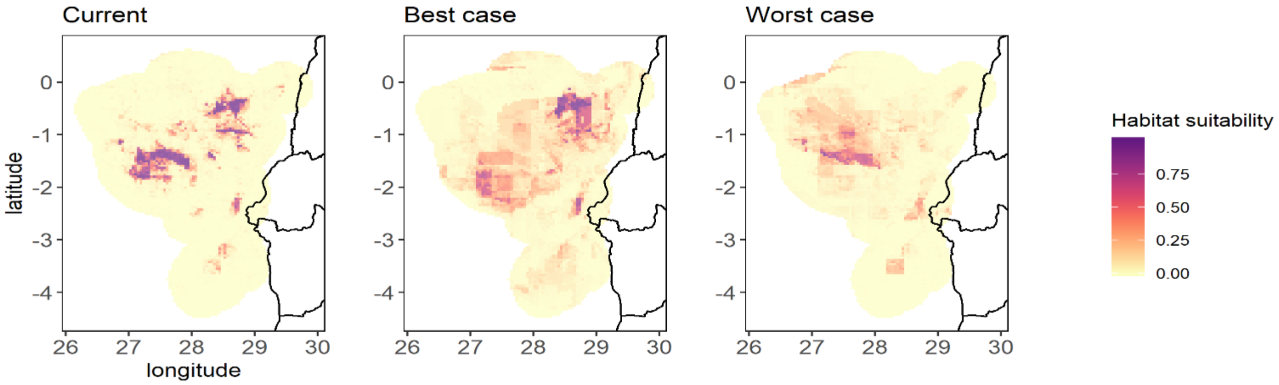


*Gorilla gorilla diehli*

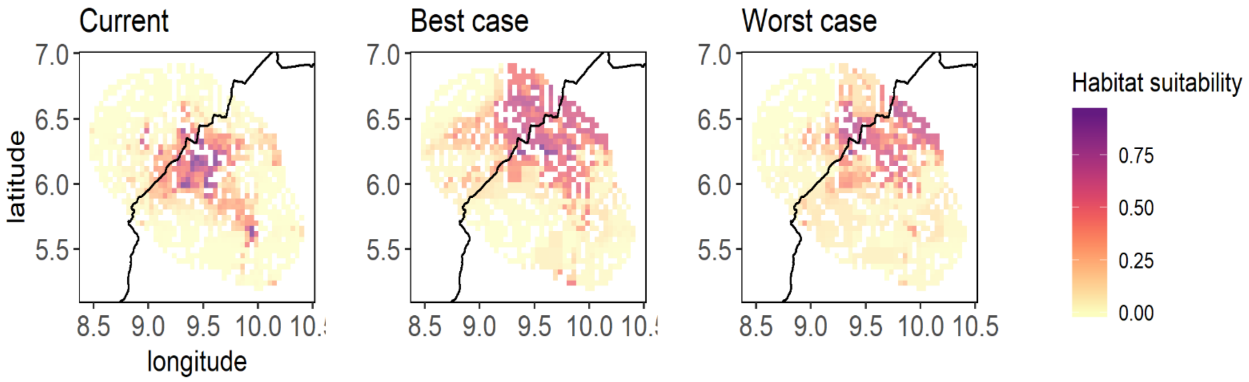

*Gorilla gorilla gorilla*

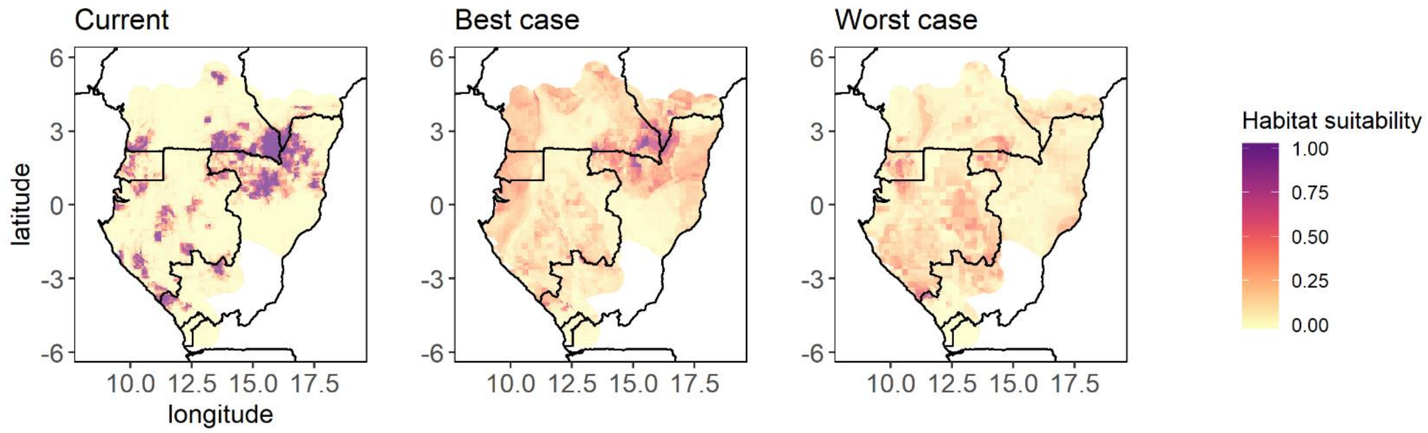


*Pan paniscus*

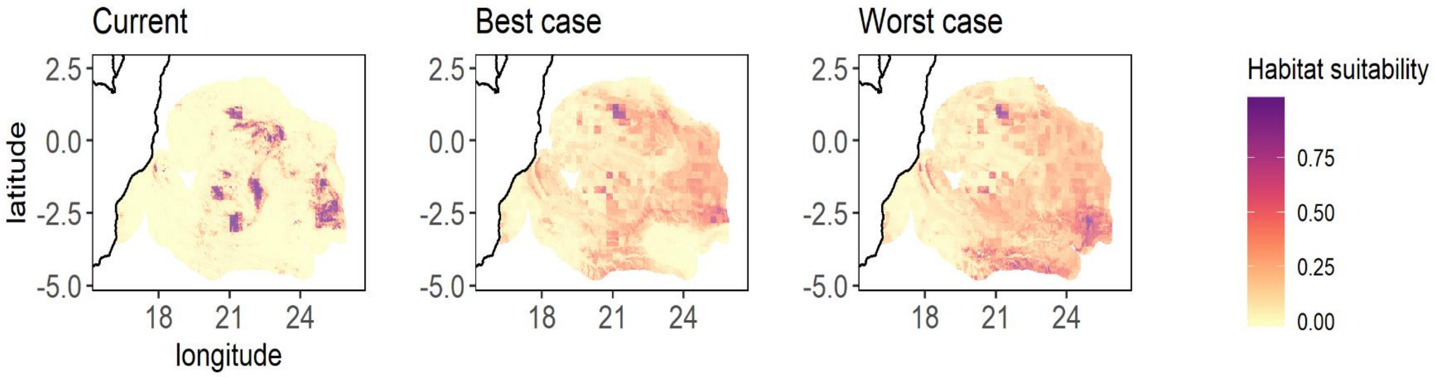


*Pan troglodytes ellioti*

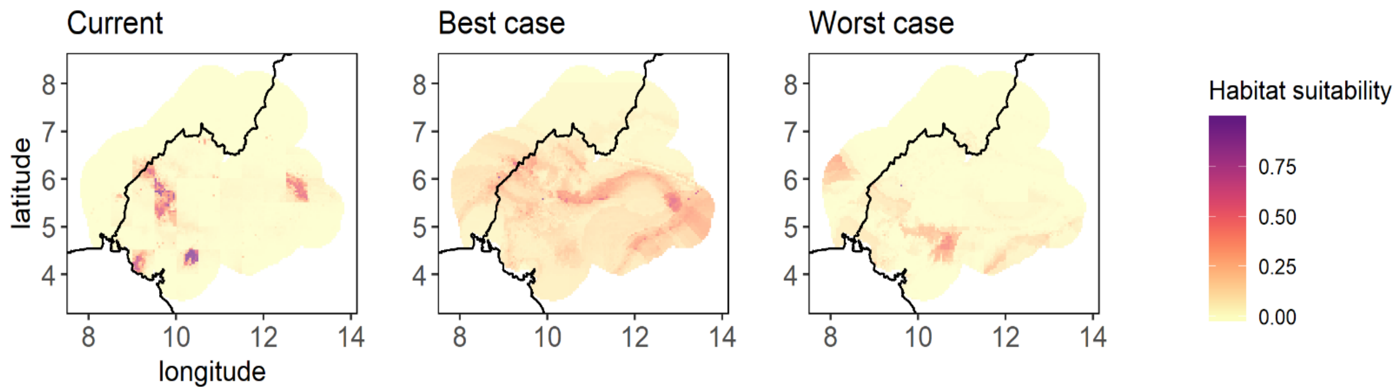

*Pan troglodytes schweinfurthii*

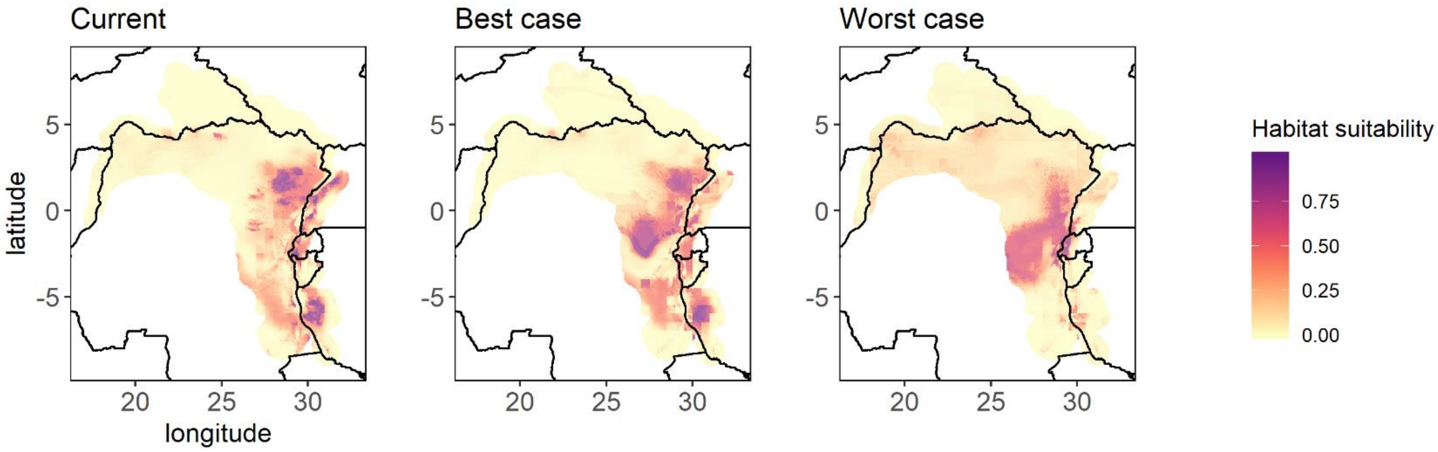


*Pan troglodytes troglodytes*

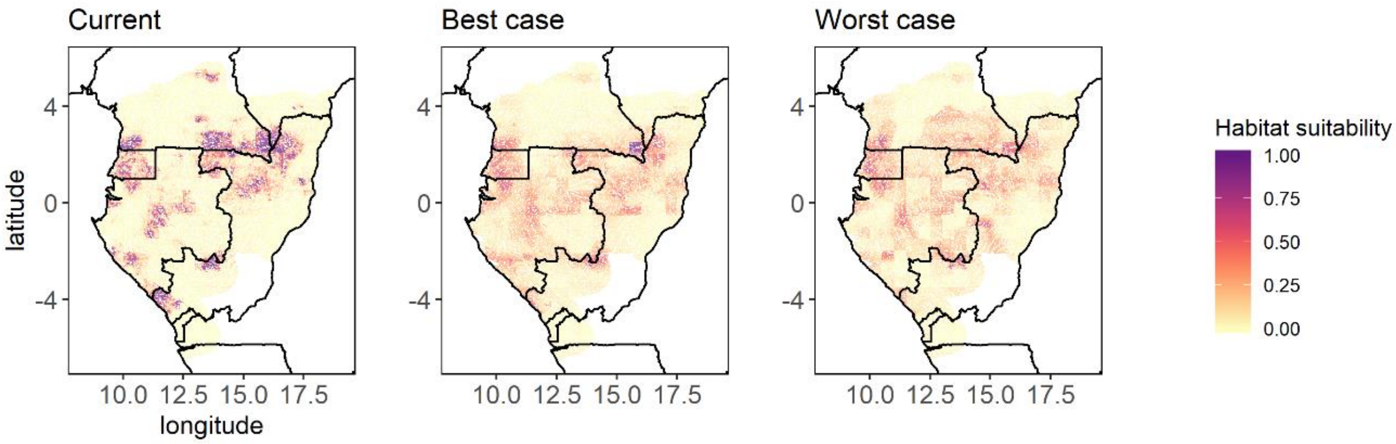


*Pan troglodytes verus*

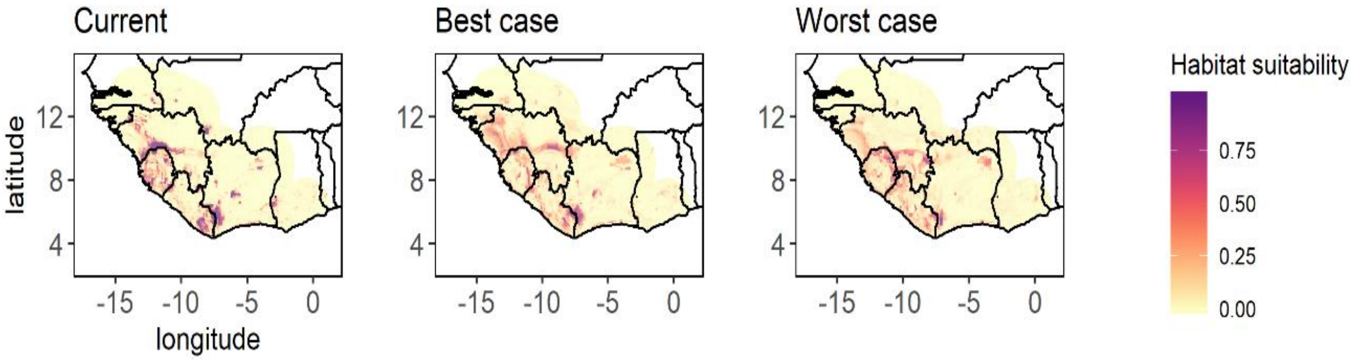


Figure S1.7. Ensemble forecasting of the current and future (best- and worst-case scenarios) habitat suitability for each taxon based on weighted mean and the true skill statistics (TSS). Note that only the no dispersal scenario is presented here.

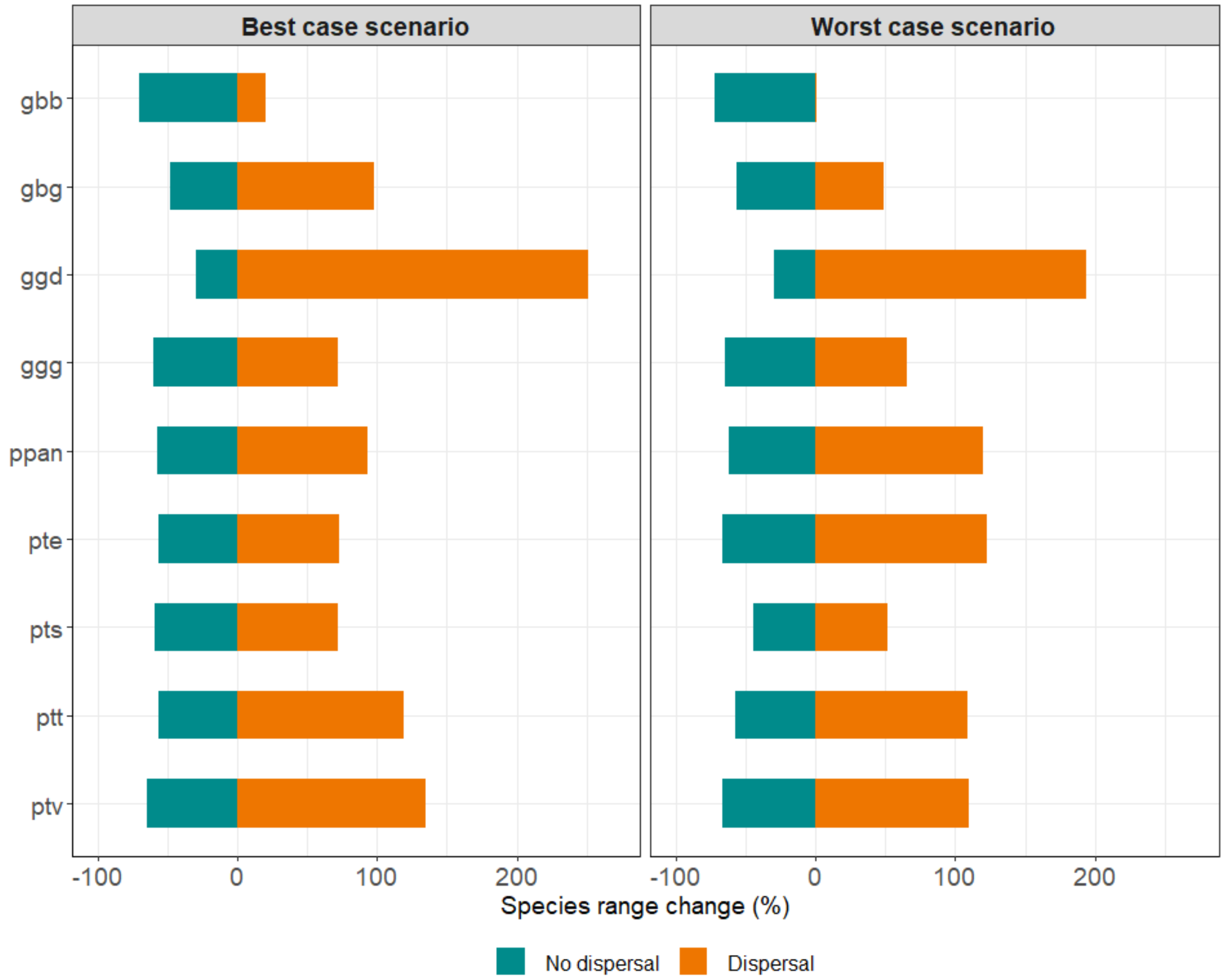


Figure S1.8. Predicted percentage change in African ape ranges by 2050 under the best- and the worst-case scenario, assuming either no dispersal (loss) and dispersal (gains) for the *default* approach.
