## Appendix 2 for "Predicting range shifts of African apes under global change scenarios"

### APPENDIX S2

##Example code for the modelling approach *non-default*, i.e. changes in the correlative algorithms GLM and GAM

*#Gorilla beringei beringei*

```
GLM = list(myFormula = formula("myResp ~ poly(gbb_bio12,2) + poly(gbb_bio19,2)
                     + gbb_bio2 + log(gbb_humpopu) + gbb_altitude1 +
                     gbb_bio12:gbb_bio19 + gbb_bio12:gbb_altitude1 +
                     gbb_bio2:gbb_altitude"),
           test = 'AIC',
           family = binomial(link = 'logit'),
           mustart = 0.5,
           control = glm.control(epsilon = 1e-08, maxit = 50, trace = FALSE))


GAM = list(algo = 'GAM_mgcv',
           type = 's_smoother',
           k = 4,
           interaction.level = 2,
           myFormula = NULL,
           family = binomial(link = 'logit'),
           method = 'GCV.Cp',
           optimizer = c('outer','newton'),
           select = TRUE,
           knots = 1,
           paraPen = NULL,
           control = list(nthreads = 1, irls.reg = 0, epsilon = 1e-06, maxit =
           100, trace = FALSE, mgcv.tol = 1e-07, mgcv.half = 15, rank.tol =
           1.49011611938477e-08, nlm = list(ndigit=6, gradtol=1e-05, stepmax=2,
           steptol=1e-04, iterlim=200, check.analyticals=0), optim =
           list(factr=1e+07), newton = list(conv.tol=1e-06, maxNstep=5,
           maxSstep=2, maxHalf=30, use.svd=0), outerPIsteps = 0, idLinksBases =
           TRUE, scalePenalty = TRUE, efs.lspmax = 15, efs.tol = 0.1, keepData
           = FALSE, edge.correct = FALSE))
```

*#Gorilla beringei graueri*

```
GLM = list(myFormula = formula("myResp ~ poly(gbg_bio7a,2) + gbg_dist2pa1 +
                      gbg_distroad + gbg_bio15a + gbg_alt +
                      gbg_dist2pa1:gbg_distroad + gbg_bio15a:gbg_alt +
                      gbg_bio7a:gbg_alt"),
           test = 'AIC',
           family = binomial(link = 'logit'),
           mustart = 0.5,
```

```
          control = glm.control(epsilon = 1e-08, maxit = 50, trace = FALSE))

GAM = list(algo = 'GAM_mgcv',
          type = 's_smoother',
          k = 4,
          interaction.level = 1,
          myFormula = NULL,
          family = binomial(link = 'logit'),
          method = 'GCV.Cp',
          optimizer = c('outer','newton'),
          select = TRUE,
          knots = 1,
          paraPen = NULL,
          control = list(nthreads = 1, irls.reg = 0, epsilon = 1e-06, maxit =
          100, trace = FALSE, mgcv.tol = 1e-07, mgcv.half = 15, rank.tol =
          1.49011611938477e-08, nlm = list(ndigit=6, gradtol=1e-05, stepmax=2,
          steptol=1e-04, iterlim=200, check.analyticals=0), optim =
          list(factr=1e+07), newton = list(conv.tol=1e-06, maxNstep=5,
          maxSstep=2, maxHalf=30, use.svd=0), outerPIsteps = 0, idLinksBases =
          TRUE, scalePenalty = TRUE, efs.lspmax = 15, efs.tol = 0.1, keepData
          = FALSE, edge.correct = FALSE))
```

*#Gorilla gorilla diehli*

```
GLM = list(myFormula = formula("myResp ~ ggd_bio4a + poly (ggd_bio13a,2) +
                     log(ggd_humpopu) + ggd_pforest + ggd_river+
                     ggd_humpopu:ggd_pforest"),
          test = 'AIC',
          family = binomial(link = 'logit'),
          mustart = 0.5,
          control = glm.control(epsilon = 1e-08, maxit = 50, trace = FALSE))

GAM = list(algo = 'GAM_mgcv',
          type = 's_smoother',
          k = 3,
          interaction.level = 1,
          myFormula = NULL,
          family = binomial(link = 'logit'),
          method = 'GCV.Cp',
          optimizer = c('outer','newton'),
          select = TRUE,
          knots = 1,
          paraPen = NULL,
          control = list(nthreads = 1, irls.reg = 0, epsilon = 1e-06, maxit =
          100, trace = FALSE, mgcv.tol = 1e-07, mgcv.half = 15, rank.tol =
          1.49011611938477e-08, nlm = list(ndigit=6, gradtol=1e-05, stepmax=2,
          steptol=1e-04, iterlim=200, check.analyticals=0), optim =
          list(factr=1e+07), newton = list(conv.tol=1e-06, maxNstep=5,
          maxSstep=2, maxHalf=30, use.svd=0), outerPIsteps = 0, idLinksBases =
          TRUE, scalePenalty = TRUE, efs.lspmax = 15, efs.tol = 0.1, keepData
          = FALSE, edge.correct = FALSE))
```

*#Gorilla gorilla gorilla*

```
GLM = list(myFormula = formula("myResp ~ ggg_bio15a + poly (ggg_bio12,2)+
                     ggg_bio2a + ggg_crop1 + ggg_dist2road1+
                     ggg_dist2road1:ggg_crop1"),
          test = 'AIC',
          family = binomial(link = 'logit'),
          mustart = 0.5,
          control = glm.control(epsilon = 1e-08, maxit = 50, trace = FALSE))


GAM = list(algo = 'GAM_mgcv',
           type = 's_smoother',
           k = 4,
           interaction.level = 1,
           myFormula = NULL,
           family = binomial(link = 'logit'),
           method = 'GCV.Cp',
           optimizer = c('outer','newton'),
           select = TRUE,
           knots = 1,
           paraPen = NULL,
           control = list(nthreads = 1, irls.reg = 0, epsilon = 1e-06, maxit =
           100, trace = FALSE, mgcv.tol = 1e-07, mgcv.half = 15, rank.tol =
           1.49011611938477e-08, nlm = list(ndigit=6, gradtol=1e-05,
           stepmax=2, steptol=1e-04, iterlim=200, check.analyticals=0), optim
           = list(factr=1e+07), newton = list(conv.tol=1e-06, maxNstep=5,
           maxSstep=2, maxHalf=30, use.svd=0), outerPIsteps = 0, idLinksBases
           = TRUE, scalePenalty = TRUE, efs.lspmax = 15, efs.tol = 0.1,
           keepData = FALSE, edge.correct = FALSE))
```

*#Pan paniscus*

```
GLM = list(myFormula = formula("myResp ~ poly(ppan_bio12,2) + ppan_alt +
                     poly(ppan_bio8a,2) + poly(ppan_bio18a,2) +
                     ppan_dist2ro1"),
          test = 'AIC',
          family = binomial(link = 'logit'),
          mustart = 0.5,
          control = glm.control(epsilon = 1e-08, maxit = 50, trace = FALSE))


GAM = list(algo = 'GAM_mgcv',
          type = 's_smoother',
          k = 3,
          interaction.level = 1,
          myFormula = NULL,
          family = binomial(link = 'logit'),
          method = 'GCV.Cp',
          optimizer = c('outer','newton'),
          select = TRUE,
          knots = 1,
          paraPen = NULL,
          control = list(nthreads = 1, irls.reg = 0, epsilon = 1e-06, maxit =
          100, trace = FALSE, mgcv.tol = 1e-07, mgcv.half = 15, rank.tol =
```

```
          1.49011611938477e-08, nlm = list(ndigit=6, gradtol=1e-05, stepmax=2,
          steptol=1e-04, iterlim=200, check.analyticals=0), optim =
          list(factr=1e+07), newton = list(conv.tol=1e-06, maxNstep=5,
          maxSstep=2, maxHalf=30, use.svd=0), outerPIsteps = 0, idLinksBases =
          TRUE, scalePenalty = TRUE, efs.lspmax = 15, efs.tol = 0.1, keepData
          = FALSE, edge.correct = FALSE))
```

*#Pan troglodytes ellioti*

```
GLM = list(myFormula = formula("myResp ~ poly(pte_bio2a,2) + pte_bio15a +
                     pte_bio4a + pte_crop + pte_pforest +
                     pte_bio15a:pte_bio4a + pte_crop:pte_pforest"),
          test = 'AIC',
          family = binomial(link = 'logit'),
          mustart = 0.5,
          control = glm.control(epsilon = 1e-08, maxit = 50, trace = FALSE))


 GAM = list(algo = 'GAM_mgcv',
           type = 's_smoother',
           k = 3,
           interaction.level = 1,
           myFormula = NULL,
           family = binomial(link = 'logit'),
           method = 'GCV.Cp',
           optimizer = c('outer','newton'),
           select = TRUE,
           knots = 1,
           paraPen = NULL,
           control = list(nthreads = 1, irls.reg = 0, epsilon = 1e-06, maxit =
           100, trace = FALSE, mgcv.tol = 1e-07, mgcv.half = 15, rank.tol =
           1.49011611938477e-08, nlm = list(ndigit=6, gradtol=1e-05,
           stepmax=2, steptol=1e-04, iterlim=200, check.analyticals=0), optim
           = list(factr=1e+07), newton = list(conv.tol=1e-06, maxNstep=5,
           maxSstep=2, maxHalf=30, use.svd=0), outerPIsteps = 0, idLinksBases
           = TRUE, scalePenalty = TRUE, efs.lspmax = 15, efs.tol = 0.1,
           keepData = FALSE, edge.correct = FALSE))
```

*#Pan troglodytes schweinfurthii*

```
GLM = list(myFormula = formula("myResp ~ pts_bio3a + pts_bio12+ pts_bio7 + poly
                     (pts_bio17,2) + pts_crop"),
          test = 'AIC',
          family = binomial(link = 'logit'),
          mustart = 0.5,
          control = glm.control(epsilon = 1e-08, maxit = 50, trace = FALSE))


GAM = list(algo = 'GAM_mgcv',
          type = 's_smoother',
          k = 4,
          interaction.level = 1,
          myFormula = NULL,
          family = binomial(link = 'logit'),
```

```
          method = ‘GCV.Cp’,
          optimizer = c(‘outer’,’newton’),
          select = TRUE,
          knots = 1,
          paraPen = NULL,
          control = list(nthreads = 1, irls.reg = 0, epsilon = 1e-06, maxit =
          100, trace = FALSE, mgcv.tol = 1e-07, mgcv.half = 15, rank.tol =
          1.49011611938477e-08, nlm = list(ndigit=6, gradtol=1e-05,
          stepmax=2, steptol=1e-04, iterlim=200, check.analyticals=0), optim
          = list(factr=1e+07), newton = list(conv.tol=1e-06, maxNstep=5,
          maxSstep=2, maxHalf=30, use.svd=0), outerPIsteps = 0, idLinksBases
          = TRUE, scalePenalty = TRUE, efs.lspmax = 15, efs.tol = 0.1,
          keepData = FALSE, edge.correct = FALSE))
```

*#Pan troglodytes troglodytes*

```
GLM = list(myFormula = formula("myResp ~ ptt_altitude1 + ptt_bio12+ ptt_river +
                     poly(ptt_bio18a,2) + ptt_sforest"),
          test = 'AIC',
          family = binomial(link = 'logit'),
          mustart = 0.5,
          control = glm.control(epsilon = 1e-08, maxit = 50, trace = FALSE))


GAM = list(algo = 'GAM_mgcv',
             type = 's_smoother',
             k = 4,
             interaction.level = 1,
             myFormula = NULL,
             family = binomial(link = 'logit'),
             method = 'GCV.Cp',
             optimizer = c('outer','newton'),
             select = TRUE,
             knots = 1,
             paraPen = NULL,
             control = list(nthreads = 1, irls.reg = 0, epsilon = 1e-06, maxit
             = 100, trace = FALSE, mgcv.tol = 1e-07, mgcv.half = 15, rank.tol
             = 1.49011611938477e-08, nlm = list(ndigit=6, gradtol=1e-05,
             stepmax=2, steptol=1e-04, iterlim=200, check.analyticals=0),
             optim = list(factr=1e+07), newton = list(conv.tol=1e-06,
             maxNstep=5, maxSstep=2, maxHalf=30, use.svd=0), outerPIsteps = 0,
             idLinksBases = TRUE, scalePenalty = TRUE, efs.lspmax = 15,
             efs.tol = 0.1, keepData = FALSE, edge.correct = FALSE))
```

*#Pan troglodytes verus*

```
GLM = list(myFormula = formula("myResp ~ poly(ptv_bio16a,2) +
                     poly(ptv_bio19a,2) + ptv_bio15a + log(ptv_humpopu) +
                     ptv_dist2ro1 + ptv_humpopu:ptv_dist2ro1 +
                     ptv_bio16a:ptv_bio19a"),
         test = 'AIC',
         family = binomial(link = 'logit'),
         mustart = 0.5,
```

```
          control = glm.control(epsilon = 1e-08, maxit = 50, trace = FALSE))

GAM = list(algo = 'GAM_mgcv',
           type = 's_smoother',
           k = 4,
           interaction.level = 1,
           myFormula = NULL,
           family = binomial(link = 'logit'),
           method = 'GCV.Cp',
           optimizer = c('outer','newton'),
           select = TRUE,
           knots = 1,
           paraPen = NULL,
           control = list(nthreads = 1, irls.reg = 0, epsilon = 1e-06, maxit =
           100, trace = FALSE, mgcv.tol = 1e-07, mgcv.half = 15, rank.tol =
           1.49011611938477e-08, nlm = list(ndigit=6, gradtol=1e-05, stepmax=2,
           steptol=1e-04, iterlim=200, check.analyticals=0), optim =
           list(factr=1e+07), newton = list(conv.tol=1e-06, maxNstep=5,
           maxSstep=2, maxHalf=30, use.svd=0), outerPIsteps = 0, idLinksBases =
           TRUE, scalePenalty = TRUE, efs.lspmax = 15, efs.tol = 0.1, keepData
           = FALSE, edge.correct = FALSE))
```
